# Evidence for kilometer-scale biophysical features at the Gulf Stream front

**DOI:** 10.1101/2023.09.25.559383

**Authors:** Patrick Clifton Gray, Ivan Savelyev, Nicolas Cassar, Marina Lévy, Emmanuel Boss, Yoav Lehahn, Guillaume Bourdin, Kate A. Thompson, Anna Windle, Jessica Gronniger, Sheri Floge, Dana E. Hunt, Greg Silsbe, Zackary I. Johnson, David W. Johnston

## Abstract

Understanding the interplay of ocean physics and biology, particularly at the submesoscale and below (<30km), is an ongoing challenge in oceanography. While poorly constrained, these scales may be of critical importance for understanding how changing ocean dynamics will impact marine ecosystems. Fronts in the ocean, regions where two disparate water masses meet and isopycnals become tilted towards vertical, are considered hotspots for biophysical interaction, but there is limited observational evidence at the appropriate scales to assess their importance. Western boundary currents like the Gulf Stream are of particular interest as these dynamic physical regions are thought to influence both productivity and composition of primary producers; however, how exactly this plays out, and at what scales, is not well known. Using satellite data and two years of detailed *in situ* observations across the Gulf Stream front near Cape Hatteras, North Carolina, U.S.A., we investigate how submesoscale frontal dynamics could affect biological communities associated with frontal regions and generate hotspots of productivity and export. In this analysis we assess the seasonality and phenology of the region, generalize the kilometer-scale structure of the front, and analyze 69 transects to assess two physical processes of potential biogeochemical importance: cold shelf filament subduction and high salinity Sargasso Sea obduction. We link these processes observationally to the meander phase of the Gulf Stream and discuss how cold filament subduction could be exporting carbon and how obduction of high salinity water from depth often leads to high chlorophyll-a. Finally, we report on phytoplankton community composition in each of these features and integrate these new observations into our understanding of frontal submesoscale dynamics.

**Plain Language Summary:** Phytoplankton move with large currents and are stirred by eddies with diameters ranging from 100s of kilometers down to the meter scale. Their growth is impacted by physical factors like light and temperature and also chemical and biological factors like nutrient availability, and their accumulation is also impacted by top down controls (zooplankton grazing, viral lysis) and competition with other phytoplankton. This interplay of physics and biology in determining the biomass and composition of phytoplankton communities is poorly understood and is key to understanding marine ecosystem resilience and structure in a changing ocean. In this work we investigated the impact of physics and biology on phytoplankton across scales focusing on the Gulf Stream front. Fronts in the ocean are where lines of equal density go from being horizontal to having a vertical tile, and because of this can enable nutrients and plankton to move from depth to the surface and vice versa. The objective of this work is to understand how physics might drive important changes in phytoplankton biomass and composition in the Gulf Stream front, which is amongst the sharpest gradients in temperature, density, and current speed in the global ocean. We find two frequent processes at the front, the apparent subduction of cold filaments down along the edge of the Gulf Stream, associated with meander troughs, and obduction of high salinity Sargasso Sea water into the front linked to meander crests. While ephemeral, these processes are frequent and could have a large impact on local phytoplankton biomass, phytoplankton composition, and the export of organic matter to depth.

**Key Points:**

- The frontal zone between the Gulf Stream and the shelf has an interface water mass which appears to have different origins with a range of biogeochemical impacts.
- Meanders appear to largely control the frontal interface: troughs lead to subduction of shelf filaments and crests lead to obduction of high salinity water.
- These two processes are common at the front and could lead to ephemeral kilometer- scale export and productivity.

## 1 Introduction

Connections between the well-lit upper ocean and nutrient rich interior fuel marine ecosystems. Regions where these connections are frequent can become hotspots of productivity, due to injections of nutrients from depth, and export, as organic matter is exported below the ocean surface for extended periods. While large-scale niches and gradients in productivity are set by environmental contrasts at the basin scale, they are rearranged by mesoscale O(30-300km) and submesoscale O(3-30km) flows (Barton et al., 2010; Lévy et al., 2015). Flows at all of these scales laterally stir existing gradients, but the submesoscale is especially efficient at driving vertical fluxes and linking the euphotic zone to the ocean’s nutrient rich interior (Mahadevan, 2016). These fluxes can then shape the composition and productivity of phytoplankton communities, cascading throughout the ocean’s food web (Lévy et al., 2018). Multiscale physical dynamics and biological responses produce the ocean’s characteristically extreme spatial and temporal heterogeneity. Yet exactly how and where these processes drive phytoplankton productivity, diversity, and export remain largely unresolved (Abraham, 1998; Klein & Lapeyre, 2009; Mackas et al., 1985; McGillicuddy, 2016).

Given the increased vertical transport at the submesoscale, submesoscale processes may help explain longstanding questions about patchy productivity and export. For example, global estimates of new production exceed the modeled nutrient availability (Klein & Lapeyre, 2009). Additionally, major effort has gone into quantifying the biological carbon pump(Siegel et al., 2016), with recent observational work suggesting an eddying flow field can drive hotspots of export - visible as small filaments of a few kilometers on eddy edges and fronts (Omand et al., 2015). However, disagreement exists with some modeling work indicating the total amount of this eddy-driven export is small, ∼5% of the annual budget (Resplandy et al., 2019). The disagreement in both cases could be due to the intermittent small scale dynamics which remain unresolved in both numerical models and most observation efforts (Couespel et al., 2021; Lévy et al., 2018; Mahadevan, 2016). For the forseeable future, it is unlikely we will be able to fully model this small scale in basin-wide or global climate models; however, understanding how to better parameterize models could vastly improve predictions in a warming world (Ferrari, 2011).

Strong persistent mesoscale fronts, such as the Gulf Stream’s north wall, are reliable hotspots of submesoscale dynamics (Mcwilliams et al., 2019) and the tilted isopycnals in these regions can provide a vertical path through the water column. These fronts are linked to increased productivity in phytoplankton, zooplankton, and fish, and represent a hotspot of ocean predator diversity (Lévy et al., 2015; Mann & Lazier, 2005; Uchida et al., 2020). The Gulf Stream front just downstream of the current’s separation point at Cape Hatteras, North Carolina, provides an ideal observatory for investigating questions of physical-biological interaction.

Although generally in geostrophic balance, western boundary currents (WBCs), such as the Gulf Stream, frequently exhibit significant ageostrophic circulation, enabling the vertical movement of nutrients and organisms at high velocities, unlike much of the rest of the ocean (Mcwilliams et al., 2019). A wide range of modeling work suggests that the ageostrophic secondary circulation at major oceanic fronts will lead to an enhancement of chlorophyll-a (chl-a) due to vertical transport of nutrients into the euphotic zone (Clayton et al., 2013; Lévy et al., 2015 and citations therein). Recent satellite observational work confirms this modeling result and suggests these mechanisms are at play (Haëck et al., 2023; Mangolte et al., 2022); however, satellite observations are restricted to the surface, at a resolution of ∼1km and are limited in their ability to mechanistically link physical processes to phytoplankton dynamics. Previous submesoscale surveys observed increased biomass and diversity at WBC fronts, but they are typically limited in temporal scope to a single short period or feature (Clayton et al., 2017). The impact of these processes on local ecology and biogeochemistry could be drastic, but they remain poorly understood due to their ephemeral nature and fine-scale. The combination of vertical injections of nutrients and lateral mixing of upstream populations that occurs at a WBC front may even lead to a unique biome at the front (Cavender-Bares et al., 2001; Clayton et al., 2014; Taylor et al., 2012). While potentially an ecologically important feature, it is poorly resolved by satellites, models, and most *in situ* sampling programs, hence motivating the present study.

This work addresses the following question: do the submesoscale dynamics associated with the Gulf Stream give rise to spatially distinct biological communities as well as localized export and high productivity patches? To assess the presence of such features we analyze the physical and biological mesoscale context across the Gulf Stream front along seasonal progression and link them to the phase of the front’s meanders.

## 2 Methods

### 2.1 Study Area

This study focuses on the Gulf Stream front, a region of oceanographic interest for centuries (Franklin & Folger, 1786) and still a frequent area for study (Andres, 2021; Muglia et al., 2022; Seim et al., 2022). The Gulf Stream is a dominant feature of the North Atlantic, the western expression of the subtropical gyre and a surface limb of the Atlantic meridional overturning circulation (AMOC). It has a major impact on climate and ocean biogeography (Longhurst, 2007; Stommel, 1965) and is associated with a nutrient core at depth thought to sustain ecosystems in the North Atlantic (Pelegrí et al., 1996). The Gulf Stream is notable for its intense submesoscale dynamics (Gula et al., 2015), making it an ideal observatory for investigating the role of submesoscale frontal dynamics on biogeochemistry and phytoplankton community composition (PCC).

We focus particularly on the region just off Cape Hatteras, North Carolina (Figure 1). This is a highly dynamic area at the confluence of the warm rapidly flowing Gulf Stream, the relatively slower, cooler, and fresher Slope Sea and mid-Atlantic Bight shelf water, spanning productive continental shelf waters out to the deep oligotrophic open ocean with a sharp change in water depth from 100m to 1000m in just a few dozen kilometers laterally. The Gulf Stream separates from the continental shelf at this point to become a meandering jet. The front on the cyclonic side of the Gulf Stream, often called the North Wall, is a region of intense gradients in temperature, salinity, chl-a, and current speed (Stommel, 1965).

**Figure 1.**
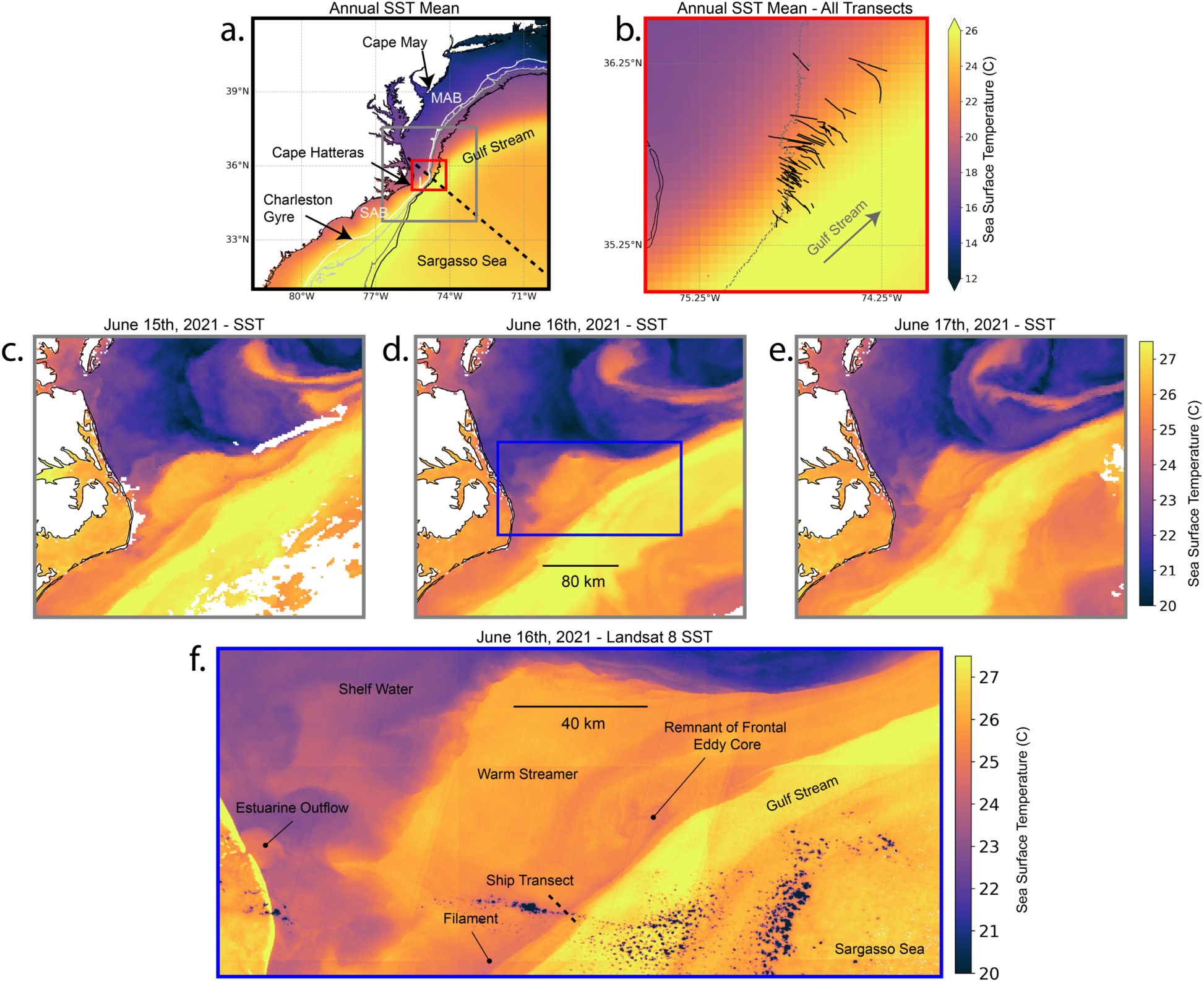
Annual mean sea surface temperature (SST) at the study area (a). The red inset is a zoom in with all transects done during this study overlaid in black (b). Examples of daily maps of SST in June of 2021 within the area indicated by the gray square of a (c-e). These days show a large meander and warm streamer passing by Cape Hatteras that was previously a frontal eddy. A higher resolution Landsat 8 SST image of the frontal region from June 16th 2021 and covering the area indicated by the blue rectangle in d (f). The annual mean SST is from the Group for High Resolution Sea Surface Temperature Level 4 Global 0.05° Nighttime Foundation Sea Surface Temperature Analysis product. The daily SST is from the Advanced Clear-Sky Processor for Ocean Global 0.02° Gridded Super-collated SST and Thermal Fronts from Low-Earth-Orbiting Platforms product. The Landsat SST is from USGS’ Earth Explorer and is derived from Band 10 at 100m resolution.

### 2.2 Satellite Analysis

We used data collected by multiple ocean-observing satellites in this analysis. Chl-a data is derived from the European Space Agency’s merged Ocean Colour Climate Change Initiative (OC-CCI)’s v5.0 daily composite product at 4km spatial resolution (Sathyendranath et al., 2019).

SST is derived from NOAA’s Advanced Clear-Sky Processor for Ocean Global daily 0.02° Gridded Super-collated SST and Thermal Fronts from Low-Earth-Orbiting Platforms product. Given the intense temperature gradients on the front, many products erroneously flag this gradient as a cloud pixel and this product was found to have the fewest erroneously flagged pixels. Landsat 8 Operational Land Imagery Surface Temperature is used for illustration purposes (Figure 1) and was downloaded from https://earthexplorer.usgs.gov/. OC-CCI chl-a was acquired from https://www.oceancolour.org/ and SST was acquired from https://coastwatch.pfeg.noaa.gov/erddap/griddap/noaacwLEOACSPOSSTL3SCDaily.html.

### 2.3 Wind Data

Wind speed data were collected via the NOAA National Data Buoy Center station 44014 as 10-minute averages from the buoy’s location at 36.603 N 74.837 W (Figure 2d).

**Figure 2.**
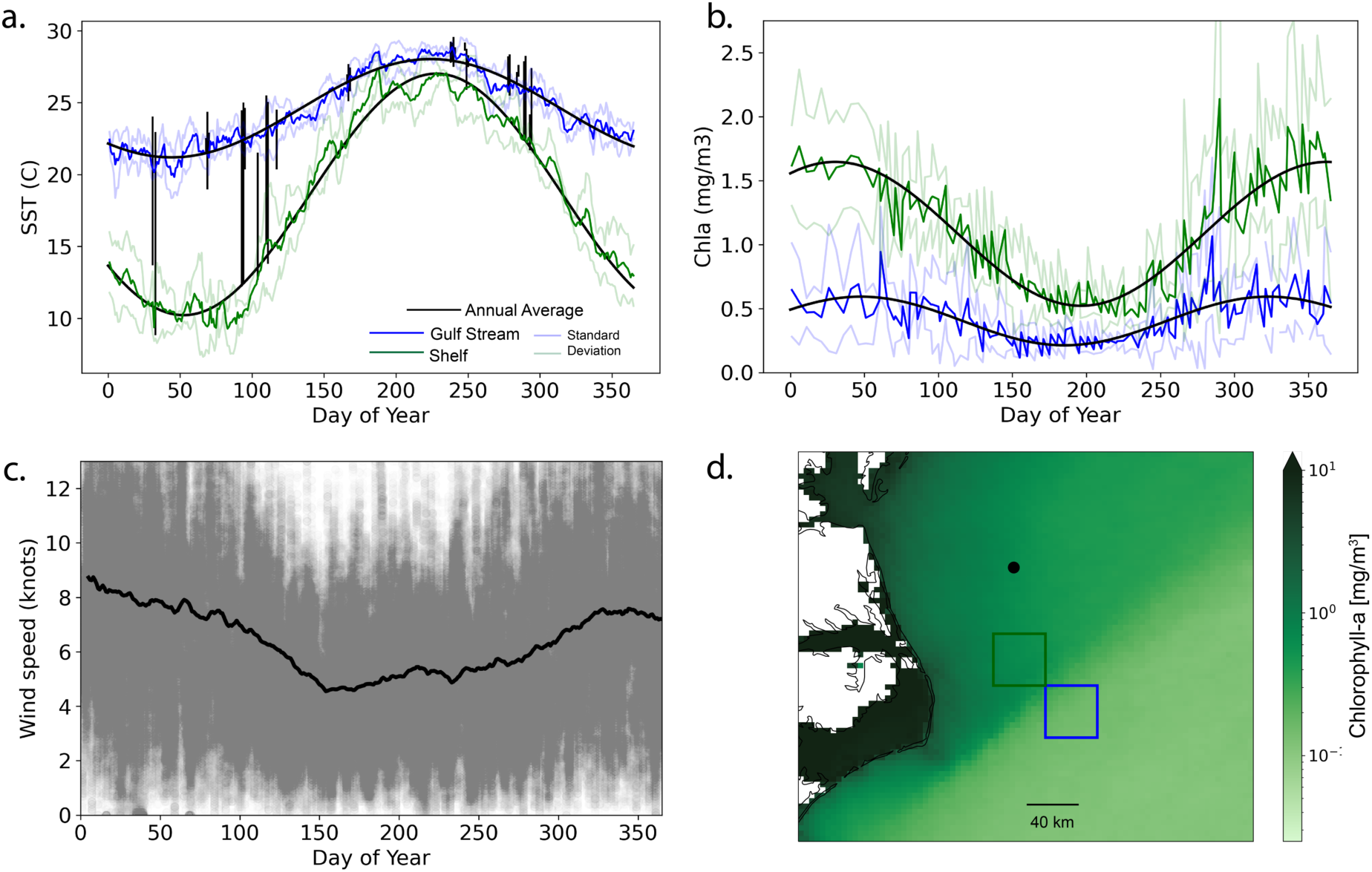
Time series of sea surface temperature (SST, Panel a) and chlorophyll-a (chl-a, Panel b) from 2018-2021 averaged from a 40 by 40 km box on the coastal side (green) and ocean side of the front (blue) delineated in panel d. Shaded lines in panels a-b show the standard deviation of all pixels within the boxes during each time step. Black vertical lines in Panel a show the range of SST measured during ship transects on those days and correspond to all the transects in Figure 1b. Panel c shows wind from the NOAA buoy indicated by the black marker in Panel d averaged from 2010-2020. Panel d shows the mean satellite-based chl-a from 2020 along with the locations for the data in the other panels.

### 2.4 Ship transects

Transect data were obtained from expeditions on-board the F/V *Instigator*, a 17 m sportfishing boat, and the R/V *Shearwater*, a 24 m regional-class research catamaran. On the F/V *Instigator,* only underway profiling and acoustic doppler current profiling were conducted. We typically transited the Gulf Stream front in approximately 5 to 20 km transects. Before beginning data collection, we would detect the front as sharp gradients in temperature and salinity with a real-time visualization from the ship’s thermosalinograph SST and visible surface roughness changes and then steam 3-8 km offshore of this point to begin the survey from that location moving inshore perpendicular to the front.

### 2.5 Underway Profiling

Vertical profiles were conducted with a Rockland Scientific VMP-250 coastal-zone profiler which collects salinity, temperature, chl-a fluorescence, turbidity (via optical backscatter at 880nm), and turbulent shear. This instrument was operated in a tow-yo mode on a winch which allowed it to freefall and then be quickly reeled back in a repeated pattern. The instrument fell to an approximate 100 m depth every 500-800m along the track. Only undisturbed measurements collected by sensors located at the front end of the instrument during down-casts were used in analyses.

### 2.6 Acoustic Doppler Current Profiler (ADCP)

Ocean velocity data was collected using a Nortek Signature 500 VM Acoustic Doppler Current Profiler which was mounted off the side of the vessel in ∼1 m depth. This instrument operates at 500kHz to collect horizontal current vectors with the 5^th^ beam measuring 1mHz acoustic backscatter profiles. From this data we calculated current speed, direction, and vertical shear along with acoustic backscatter strength. Vertical profiles of these measurements were binned in 1 m resolution from ∼2 to 60 m depth.

### 2.7 Temperature and Salinity

Surface temperature and salinity data were collected with a SeaBird SBE45 Thermosalinograph collected via the R/V *Shearwater*’s flow-through system from an intake at approximately 1.5 m depth. This data was logged once per second. The F/V *Instigator* cruises had no flow-through setup however, the ADCP unit was equipped with an in-water temperature sensor, measuring temperature transects at 1 m depth on both vessels.

### 2.8 Flow-through Optical Properties

Hyperspectral absorption and attenuation (400 to 735 nm at ∼4 nm spectral resolution, AC-s, Seabird Sci.), and the volume scattering function (VSF, Eco-BB3, Seabird Sci., installed in a 4.5 L box) at 120 deg and 470, 532, and 650 nm were measured continuously for a subset of transects. A 0.2 *μ*m filter cartridge was connected to the system and we manually redirected the flow to measure the properties of filtered seawater in both the ACS and BB3 for about 10 minutes before and after each transect, whereas total (“normal”) seawater was flowing the rest of the time. Absorption, attenuation, and backscattering measurements recorded during the filtered periods were linearly interpolated across the transect and subtracted from the total seawater measurements to obtain an estimate of the particulate absorption (ap), attenuation (cp), and backscattering (bbp). This setup allows retrieval of particulate optical properties independently from the instrument drift and biofouling (Slade et al., 2010), and assumes dissolved properties vary linearly across the front (more detaisl in Text S1 and Table S1).

These inherent optical properties were used as proxies for a range of particulate properties (Table S1). These include chl-a line height, a chl-a estimate derived from the absorption peak at 676 nm (Boss et al., 2007; Roesler & Barnard, 2013), gamma, which is a robust proxy for mean particle size with a higher gamma indicating smaller average particle sizes and lower gamma indicating larger average particles (Boss et al., 2001), and phytoplankton pigment concentrations derived from a gaussian decomposition of the particulate absorption spectra (Chase et al., 2013). This approach uses a series of gaussians placed at the same location as expected for various pigment absorption peaks and minimizes the difference between a spectrum constructed of these gaussians with the measured spectrum. This gives the approximate concentration of a range of pigments that can be used as a proxy for various phytoplankton groups.

This data was collected using Inlinino (Haëntjens & Boss, 2020) an open-source logging and visualization program, and processed using InlineAnalysis (https://github.com/OceanOptics/InLineAnalysis) following (Boss et al., 2019), further details in Text S1.

### 2.9 Surface Front Delineation

Daily Gulf Stream SST was modeled by fitting a sine curve to the satellite SST time series in Figure 2A (blue line) derived from the blue box in Figure 2D. This model is shown as the top black line in Panel 2A. This model provided daily Gulf Stream SST estimates and anything within a 0.5° C buffer of this SST was considered Gulf Stream surface water. Shelf water was modeled the same way via the green box in Figure 2D. In between these two thresholds was considered interface water.

### 2.10 Identification of High Salinity Sargasso Sea Water

We used a salinity threshold of 36.35 PSU to identify deeper Gulf Stream and Sargasso Sea water. This value is higher than the salinity we see all year in the Gulf Stream core.

### 2.11 Identification of Shelf Filaments

We identified shelf filaments based on a non-monotonic SST gradient when moving from the Gulf Stream towards the shelf.

## 3 Results

### 3.1 Seasonality and phytoplankton phenology at the front

Chl-a and SST generally follow a seasonal subtropical pattern common at this latitude of the Atlantic Ocean with highest chl-a in mid-February and lowest in mid-August (Figure 2). This matches the pattern in wind speed with its minima in mid-August and maxima in mid-February (Figure 2c). SST is opposite in phase (Figure 2b). While coastal waters in this region have more diverse origins (shelf, slope, estuarine outflow) they follow a similar pattern in both chl-a and SST as the Gulf Stream, but with a more intense seasonality. Salinity also has a slight seasonal dependence with higher salinity in winter (Figure S3). While nutrient data is limited to only two cruises (one each in September 2021 and March 2022, Figure S4), it suggests that both sides of the front that in late summer nitrate (NO3) is limiting (<0.1*μ*M, N:P ratio of 1:1), and in late winter there appears to be a NO3 surplus for most phytoplankton (∼0.5*μ*M, N:P ratio of 50:1) however, growth may be limited by phosphate (∼0.01*μ*M; often at or below the detection limit) (extended discussion in Text S4).

### 3.2 Multiple Scales of Biophysical Interaction

Layered on top of this seasonal pattern are mesoscale Gulf Stream meanders and frontal eddies which both impact SST and chl-a at the front through advection and vertical processes (Lee et al., 1991). When looking at a cross-section of the front through time, the interweaving of this seasonality and mesoscale features is seen as periods of cooler higher chl-a water displaced onto the left of the front line and warmer lower chl-a water displaced to the right of the front as a function of time (Figure 3). Vertical profiles collected down to ∼80 m at sub-km lateral resolution demonstrate a large spread beyond these events, indicating importance of submesoscale and fine-scale processes.

**Figure 3.**
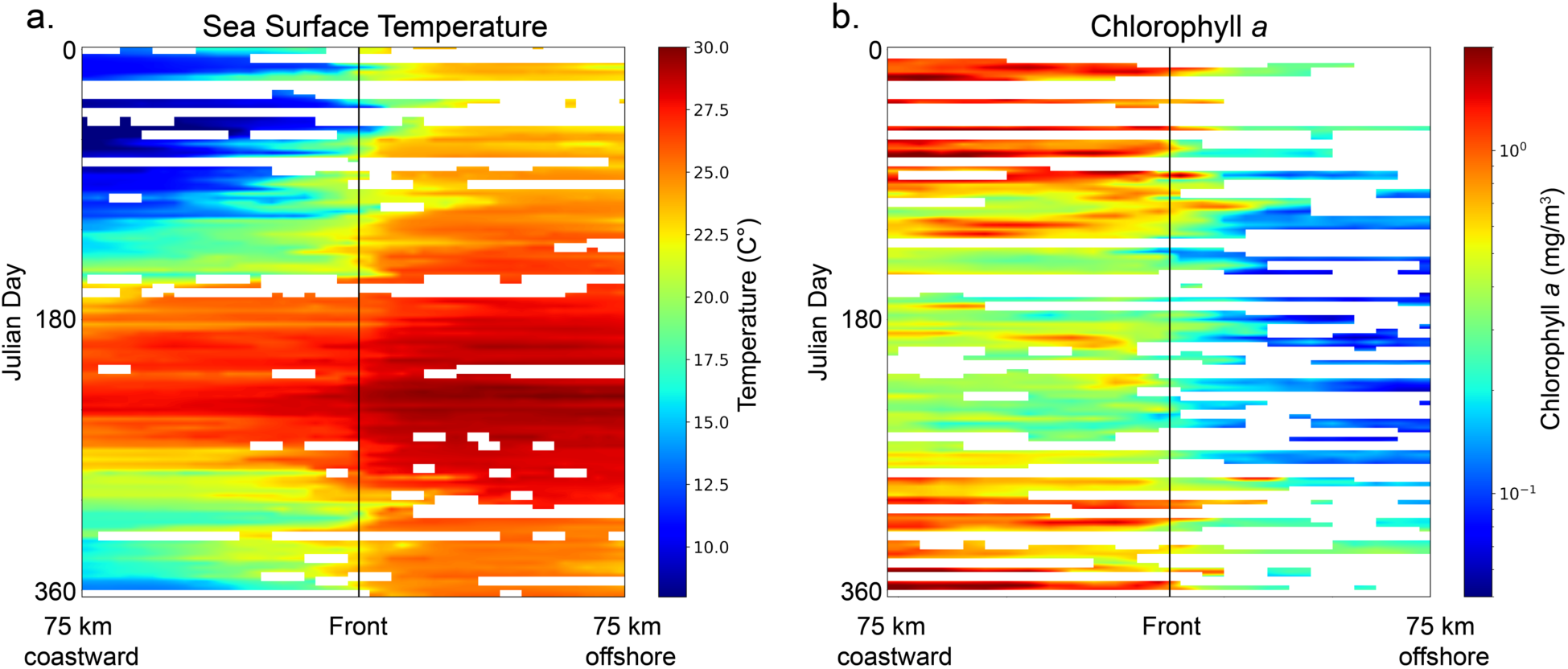
Hovmöller plot for satellite sea surface temperature (SST, a) and for chlorophyll-a (chl-a, b) from 2021. This is calculated via a 150 km line perpendicular to the Gulf Stream (x-axis) and shown through time (y-axis; DOY- day of year). The black line is the average Gulf Stream front centered on the location of our surveys (35.626, -74.738). It shows both the seasonality of SST and the meanders and eddies that propagate through the region on a regular basis. These can be identified as the daily and weekly variability around the black line indicating the average front. On the right is a Hovmöller plot for chl-a from 2021 calculated via the same approach. Cloudy pixels were masked and are shown as white in these plots. Both products were resampled to two-day averages.

### 3.3 Processes at the front

The Gulf Stream surface front is a complex interface (Figure 1), with fronts not always aligned in all parameters (e.g. the SST front, current front, and ocean color front are not always in the same location). Not only is there a decoupling between different parameters, but the Gulf Stream is rarely directly adjacent to shelf waters in this region. Instead, there is a water mass at the interface, effectively the front itself, that appears to be formed by a range of processes and with different biogeochemical implications. All transects used in this analysis can be seen in Figure S5.

In 20 of our 45 front-crossing transects, the SST gradient is non-monotonic across the front with a minima near the peak current gradient (Figure 4a). Depth profiles show a cooler, fresher, generally higher biomass filament being entrained at the front (Figure 4, Panels e-i). Often, these cold filaments are visible at depths >30 m, though in the summer they may only penetrate the top 10 m of the water column.

**Figure 4.**
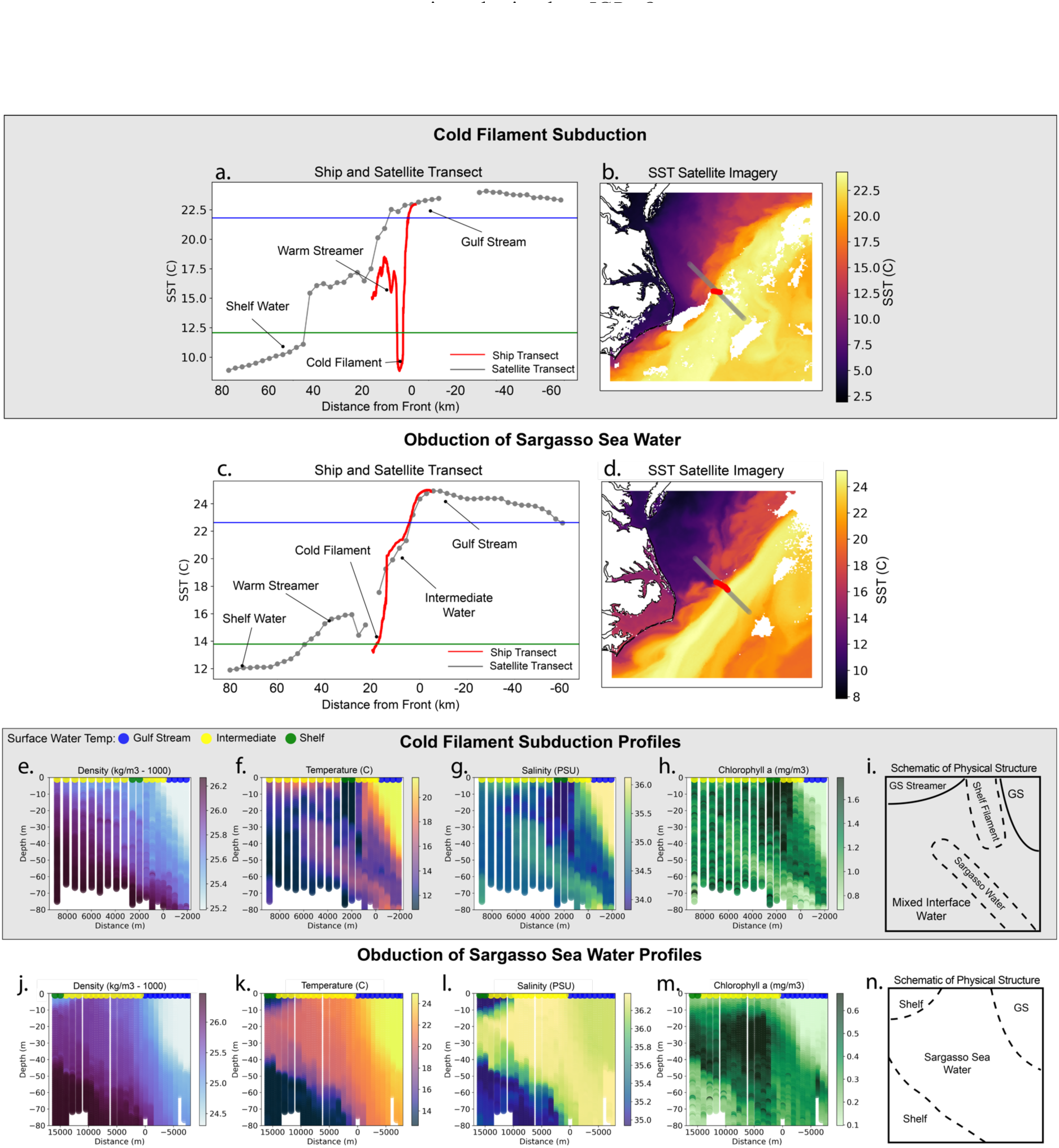
Examples of a cold filament likely being entrained and subducted at the Gulf Stream front and a period of obduction of Sargasso Sea water. The 1-dimensional transect views (Panels a and c) show the ship transect in red and a longer satellite derived transect in grey. The blue and green lines indicate the Gulf Stream and shelf water temperature thresholds respectively. The 2- dimensional surface satellite views (Panels b and d) show the mesoscale context. The 2- dimensional depth view is shown first as a schematic (Panels i and n) and in measured parameters corresponding to the red surface transect shown (in Panel a and b).

In 32 of our 45 front-crossing transects, a higher salinity water than the Gulf Stream core is present in the top 60 meters of the water column (Figure 4, Panels j-n). Typically, this higher salinity water has higher chl-a than the Gulf Stream and at times higher chl-a than the shelf water. In all cases where there is available satellite data this higher salinity water occurs near the crest of the meander, though sometimes extending a few kilometers along the upstream edge of the trough (upstream relative to the crest).

### 3.4 Phytoplankton communities across individual transects

Continuous measurements of absorption, attenuation, and backscattering of light and derived optical proxies reveal changes in PCC and particle properties across the Gulf Stream front in three different cases: cold filament entrainment (Figure 5, left column) and obduction of Sargasso Sea water (Figure 5, right column), as well as case with a frontal eddy (Figure 5, center column)

**Figure 5.**
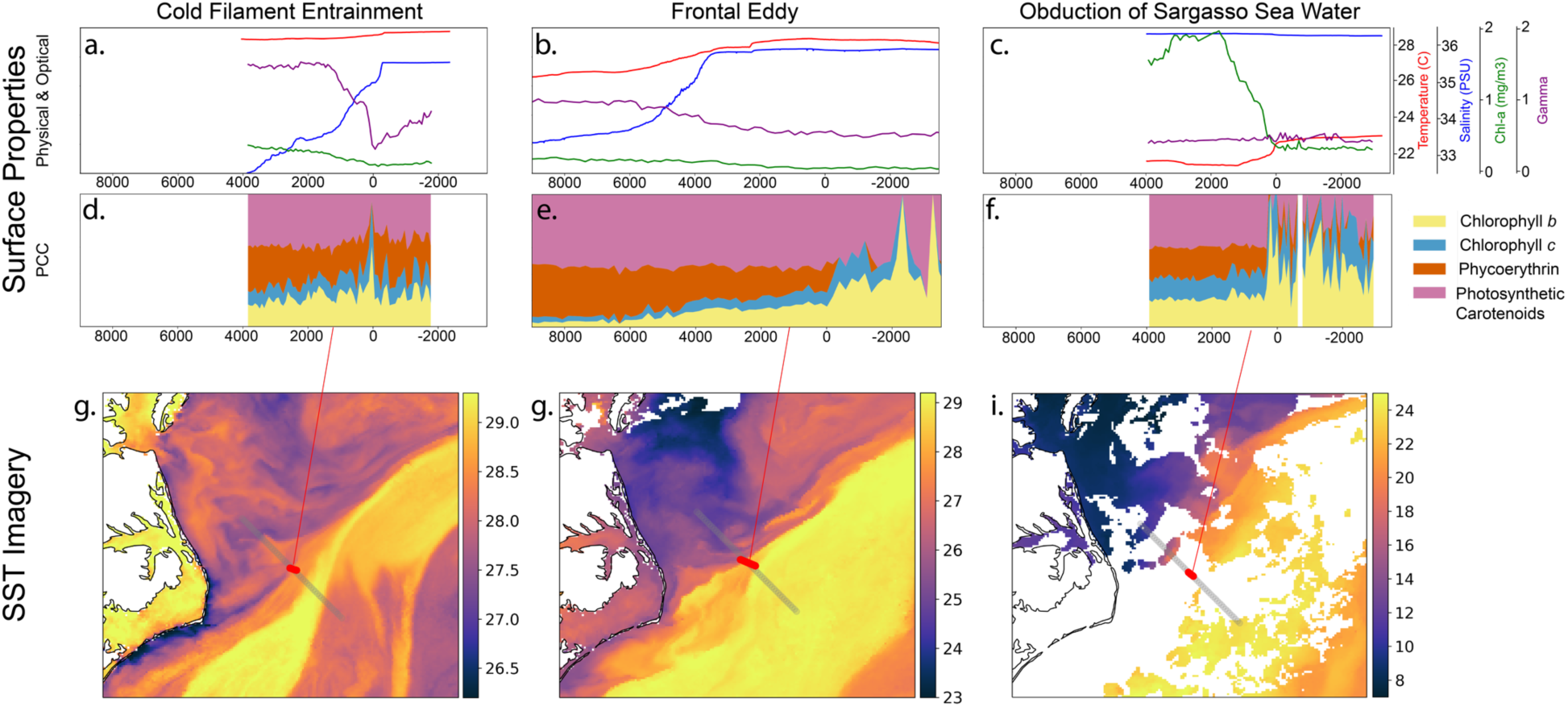
Relative pigment-based surface phytoplankton community composition (PCC) during three different frontal transects affected by different processes across the front. Surface properties are shown on the top row (a-f). Satellite SST is shown in the bottom row (g-i). The left column is crossing a cold filament and into the Gulf Stream, the middle column is crossing from a shelf filament into a frontal eddy, and the right column is crossing from a warm streamer into upwelled Sargasso Sea water and finally into Gulf Stream water. See Figure S2 for 2D context in the form of vertical profiles and a schematic.

In the cold filament case (Figure 5, Panels a,d,g) we observe low surface chl-a and relative PCC is mostly stable except for an anomaly directly at the front. Gamma (optically-based particle size proxy) has its minima at the front indicating a peak in particle size while the relative phycoerythrin concentration (likely *Synechococcus,* (Kramer & Siegel, 2019)) and the relative photosynthetic carotenoid pigment concentrations have a minima. These patterns are consistent with the concept of a converging front subducting, with larger buoyant particles collecting on the surface at the front.

In the high salinity obduction case (Figure 5, Panels c,f,i) as we move from this high salinity water to the Gulf Stream we see a large decrease in chl-a and a major shift in relative PCC. The change in this transect is not gradual or a single anomaly, but rather a step change directly at the front. In the Gulf Stream core there is more chlorophyll-b (chl-b, possibly prasinophytes, chlorophytes, or *Prochlorococcus*, (Kramer & Siegel, 2019)), less phycoerythrin (likely *Synechococcus*), and higher variability. The chl-a in this obducted Sargasso Sea water (∼2 mg/m^3^) is nearly the highest observed in all two years of frontal transects and from our flow cytometry it appears much of this is in the larger nano-plankton group (Figure S4). From SST this could appear as a shelf-based cold filament, but the surface salinity is higher and vertical profiles indicate it is actually Gulf Stream or Sargasso Sea derived (Figure S2).

In the frontal eddy case (Figure 5, Panels b,e,h) we traversed between the meander crest and trough, closer to the crest. Our measurements show a large gradual change in PCC across this transect with a decrease in phycoerythrin (likely *Synechococcus*) and increase in chl-b (possibly prasinophytes, chlorophytes, or *Prochlorococcus*). Gamma indicated an increase in particle size moving from the shelf into the eddy water, the opposite of our expectations. An increase in non-algal particles such as detritus or mineral particles could explain this finding.

## 4 Discussion

While typical observational approaches obscure the front, we report on an entire subfrontal scale where isopycnals that lead deep into the Sargasso Sea often outcrop. It may be this subfrontal region where much of the important biogeochemical role of major fronts is occurring. We focus on two processes driven by the meander phase: subduction and obduction - which could have major importance for net primary productivity, plankton community composition, and carbon export.

The present study links concepts from previous work studying WBC meanders and vertical movement (Olson et al., 1994), recent insight into the fine-scale frontal interface (Klymak et al., 2016), and observations of the Gulf Stream’s nutrient sublayer (Csanady & Hamilton, 1988; Pelegrí et al., 1996; Stefánsson & Atkinson, 1971) with combined high resolution *in situ* sampling and satellite analyses.

### 4.1 Meander Driven Entrainment and Detrainment

Ecologically, the Gulf Stream is often assumed to be a barrier at scales larger than 150 km (Bower et al., 1985), but instabilities of the front and departures from a geostrophic balance not only inject nutrients, but also drive the creation of submesoscale filaments, eddies, and streamers on both sides of the Gulf Stream (Gula et al., 2015). One mechanism that has been suggested to drive frequent vertical fluxes on the Gulf Stream front is the phase of the Stream’s meandering (i.e. crest vs trough) (Bower, 1989; 1991). The meander structure is thought to lead to convergence on the upstream side of a meander (trough) along with frontogenesis and entrainment of shelf water, which due to the converging frontal structure can be quickly subducted along isopycnals (Olson et al., 1994). The opposite process is described to occur on the downstream side of the meander (crest) where divergence occurs along with frontolysis, often associated with a warm streamer, allowing for positive vertical velocities, possibly from the nutrient core of the Gulf Stream. This framework has been demonstrated via modeling as well as observations (Mcwilliams et al., 2019). Recent work just south of our study region shows meanders driving positive vertical velocities up to 10 cm/s at the crest and downward velocities of the same magnitude in the trough (Muglia et al., 2022). This process increases horizontal and vertical gradients, likely increasing diapycnal mixing during periods of frontogenesis (Olson et al., 1994).

We connect our observations to the meander phase and find that they often help explain and contextualize the major processes observed in our transects, namely troughs associated with obduction of high salinity water and crests associated with entrainment of cold filaments (Figure 6).

**Figure 6.**
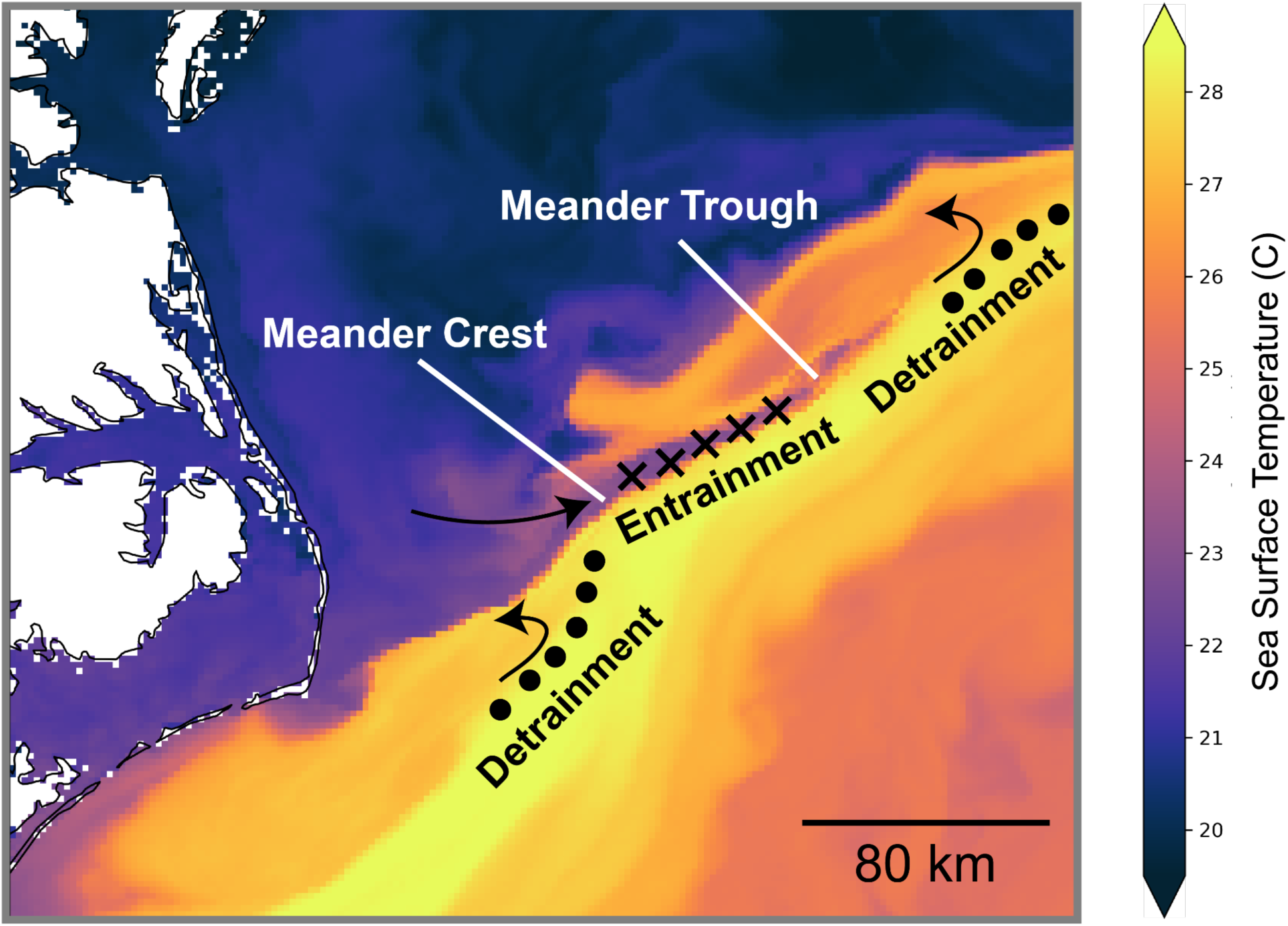
Gulf Stream meander driven convergence and divergence. Dots indicate upwelling and crosses indicate downwelling. Figure inspired by Olson (1994). Sea surface temperature (SST) is from Sept 15^th^, 2020 and is from the Advanced Clear-Sky Processor for Ocean Global 0.02° Gridded Super-collated SST and Thermal Fronts from Low-Earth-Orbiting Platforms product.

### 4.2 A Conceptual Model for Gulf Stream Frontal Structure

The frontal interface of the Gulf Stream, often considered a singularity, can have a width of ∼5-10 km across, and has been described physically in previous work (Klymak et al., 2016). This interface water and small-scale mixing within it, have been suggested as the dominant mechanism for exchange between the Sargasso Sea and the subpolar North Atlantic (Klymak et al., 2016) but it is not fully understood or resolved in most models (Wenegrat et al., 2020). Water within this interface is often trapped long enough within the meander to be completely mixed (Klymak et al., 2016) with symmetric instability possibly playing a role (Thomas et al., 2013).

Klymack et al. (2016) used observations and simulations to describe frequent streamers and intrusions exchanging water at the Gulf Stream front. The authors associated this process with the meander phase of the front. High salinity streamers mix with shelf water and are carried away from the Gulf Stream at the crests, and entrainment of fresher water occurs at the trough. This helps to maintain the density gradient of the front itself and explains why the front interface does not widen. McWilliams et al. (2019) argue this process leads to a sharper front in the trough and just upstream of the trough, with a wider front at the crest that is more prone to submesoscale perturbations.

In the context of our observations and the growing literature on the frontal interface, we present a generalized conceptual model of the front structure based on all 45 frontal transects (Figure 7). We highlight the two front-associated features observed frequently in our work connected to meander phase: subduction of shelf filaments entrained at the front and upwelling of apparent Sargasso Sea water on the coastal side of the Gulf Stream associated with the detrainment of warm streamers.

**Figure 7.**
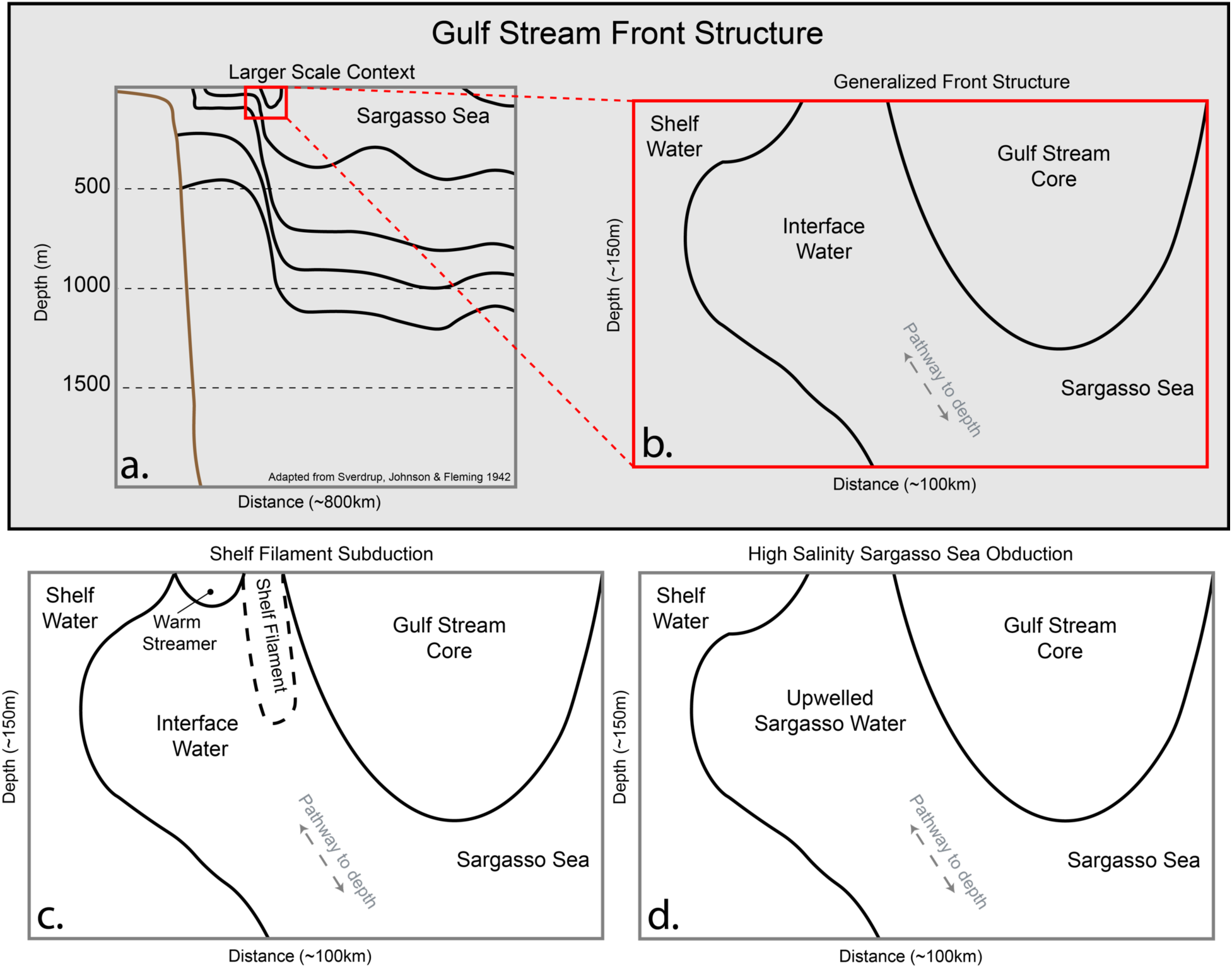
Based on all transects across the front, the shelf and Gulf Stream waters are rarely directly adjacent. An interface water mass is typically at the front with highly variable origins and properties. This interface water is in some ways a representation of the front itself, this is generally where the isopycnals are nearly perpendicular to the ocean surface and provides a density pathway for waters from depths under the Gulf Stream to the surface. a) larger scale context from the coast out into the middle of the Sargasso Sea. b) A diagram of this general structure. c) schematic of transect during cold filament subduction. d) schematic of transect during the upwelling of Sargasso Sea water at the interface. Panel a is adapted from (Sverdrup et al., 1942).

### 4.3 Obduction of Sargasso Sea Water

Previous work has revealed a nutrient core, sometimes termed the “nutrient bearing strata”, of the Gulf Stream and Sargasso Sea in the isopycnal range between 26.5 and 27.3 σT (σT = ρ(S,T) - 1000 kg m^−3^) (Pelegrí et al., 1996). There has been frequent speculation that this must be a principal factor in the high productivity of the subpolar Atlantic (Pelegrí et al., 1996) with seasonal outcroppings of this density layer bringing this water mass into the euphotic zone leading to patchy high productivity. This has been thought to occur above 45° N with reports that this isopycnal is a few hundred meters deep off Cape Hatteras, North Carolina (Csanady & Hamilton, 1988). The Gulf Stream and Sargasso Sea waters off the North Carolina coast have been shown to increase dramatically in NO3 and PO4 around 25.5 σT (Stefánsson & Atkinson, 1971). Directly off Cape Lookout, North Carolina NO3 and PO4 concentrations peak ∼100 m depth in February and ∼200 m depth in November (Stefánsson & Atkinson, 1971), though these observations were not related to Gulf Stream meanders which could have a large influence on the depth of the nutrient bearing layer in addition to seasonal density changes. While, to our knowledge, this has not previously been connected, the steep isopycnals in the Gulf Stream in lower latitudes (<40 N) where there is sufficient light all year for phytoplankton growth, could be a major source of local productivity.

This expression of dense high salinity water that we observe and assume to be part of the Gulf Stream and Sargasso Sea’s nutrient strata has, to our knowledge, not been documented to occur near this latitude. The outcropping of these isopycnals is typically thought to occur >45° N (Pelegrí et al., 1996). We observationally link this feature occurring within the top 20-60 m of the water column to the detrainment of Gulf Stream water at the crest of the meander which leads to local divergence, and thus obduction of water from depth. This warm streamer typically separates from the meander and can be left stranded on the shelf (Lee et al., 1991) or re-entrained in the trough of the meander. Increasing summer stratification may prevent this deeper Sargasso water from reaching the surface, though it is often still visible at >50 m depth in the summer. While this may limit the impact of this process on phytoplankton growth, we do still observe high chl-a in this water mass even when ∼50 m depth.

There is the possibility that enhancement of chl *a* in obducted water could come from the chlorophyll max, but we suggest the deep chlorophyll max is likely sustained and enhanced in this region by the linkage to the nutrient strata and that it would be challenging to disentangle the two.

Given the lateral extent of this high salinity feature (∼5-10 km) this would not be visible via satellite microwave salinity measurements which typically have a 25-40 km resolution. Often, this feature is below the surface and thus not visible at all to satellites or even *in situ* surface salinity measurements. At 50 m depth, where we see it in the summer, the chl-a enhancement from this process would mostly be hidden from passive ocean color remote sensing as well. While SST satellites have sufficient spatial resolution, there is no temperature signature because this water mass would appear as a mixture of shelf and Gulf Stream water origins rather than deeper Gulf Stream or Sargasso Sea water.

This conduit to the Gulf Stream and Sargasso Sea nutrient strata could enhance primary production at the front and could be a large component of increased productivity at the front seen in models and satellite data (Haëck et al., 2023; Mangolte et al., 2022). To constrain the total impact of these events, further work is needed to pinpoint the along-stream extent of this process, over what duration the high vertical velocities exist, and under what conditions it leads to growth enhancement. If the proper *in situ* work is done to gain this understanding we may be able to leverage satellite-based detection of the meander phase to accurately estimate their role in regional biogeochemical cycles.

### 4.4 Subduction of Cold Filaments

Our observations suggest all cold filaments visible at the surface are likely connected to the meander trough entraining the filament and convergence at the front pulling the cold filaments along layers of equal density (Figure 4a and Figure S5). Gravitational export can be enabled when isopycnals tilt towards the vertical and submesoscale eddies appear to contribute a large amount of export in the North Atlantic (Omand et al., 2015). Given the intense submesoscale vertical motion and nearly vertical isopycnals, this is an obvious candidate for export, but it is not clear how regularly the shelf water is subducted to depth. Even if these filaments have the opportunity to travel to depth, long-term export is not the only outcome as filaments could be sheared apart by and mixed into the surface core of the Gulf Stream. The entrainment of cold filaments at the front has been observed previously (Gula et al., 2014; Todd, 2020), and one study in this region that observed subduction of a cold filament inferred this was a vertical circulation in only the top 200 m of the water column using the omega equation (Thomas & Joyce, 2010). Seasonality is likely to play a large role in the durability of export, our summer example of a cold filament shows no subduction and instead does not penetrate below 10 m (Figure S2).

It is unclear how the export from convergence of MAB and SAB waters at the Hatteras Front (Todd, 2020), export via cascading events due to atmospheric cooling (Text S5) (Han et al., 2021), and the entrainment of (and possible export) shelf water into meanders interact, or how important each is for export in this region. It appears some of the glider observations in Todd (2020) could possibly be explained by the meander-based entrainment and detrainment. All three of these processes could combine to export substantial carbon and may well sustain mesopelagic ecosystems that in turn support the large populations of deep feeding top predators such as beaked whales and pilot whales known to aggregate in this region.

### 4.5 Connection to Meander Phase

The meander phase in this region can be challenging to identify because of the shift in Gulf Stream dynamics as it separates from the continental shelf and moves north of Cape Hatteras (Gula et al., 2015; Muglia et al., 2022). Notably, there is a decrease in meander amplitude and shearing apart of frontal eddies and warm streamers at this point (Andres, 2021). Additionally, detraining streamers and entraining filaments can be layered upon each other (Figure 1c).

Despite this local complexity, in nearly all of our observations the entrainment of cold filaments and detrainment appear to be associated with troughs and crests respectively. We do not quantify the exact proportion of crests with high salinity Sargasso Sea water and troughs with cold filaments because they may not be captured during *in situ* sampling; however, where our data spans the full front interface this connection exists. Meanders are reported to move through this region every 5-15 days (Andres, 2021; Glenn & Ebbesmeyer, 1994a) and given the continued detrainment and entrainment this could be a large and even dominant conduit between the well-lit surface and deeper nutrient rich water in this region.

Further study is needed, in particular, to identify cold filament subduction depths. This could possibly be done with a density following float. Quantifying nutrient input by the obduction of high salinity water is another priority, ideally with nutrient samples along the same isopycnal from the surface to below the euphotic zone.

A large body of observational and modeling studies have shown that meander crests and troughs have a strong impact on the ageostrophic circulation in the Gulf Stream (Bower et al., 1985; Mcwilliams et al., 2019) though impacts on chl-a are not always obvious from satellite and model output (Gaube & McGillicuddy, 2017). Varying stratification, meander asymmetry, wind events, and lags in biological response may make it hard to identify these features from satellites and this combined with low spatiotemporal resolution may be why previous work did not identify consistent chl-a patterns along with meander structure. Adding to the challenge, the depth at which we see the enhancement occurring in the summer and the cloudiness during winter when the obducted high chl-a water is at the surface, may prevent this from being evident in satellite ocean color. Further work connecting meander crests to chl-a enhancement may need to use daily 1 km resolution ocean color to prevent obscuring the reported submesoscale meander crest driven enhancement and an ocean color LiDAR could be a key piece of unraveling this puzzle.

While frontal eddy formation may include substantial upwelling as indicated by a cold dome and uplifted isotherms, from a Eulerian perspective off Cape Hatteras, the majority of positive vertical velocities in the top 150 m appear to occur at the crest of the meander (Muglia et al., 2022), which may be cause to reconsider assumptions and applicability of previous work further upstream in the SAB to this region and downstream (Glenn & Ebbesmeyer, 1994b; Lee et al., 1991; Yoder et al., 1981). The physical interplay and amount of upwelling driven by divergence at the crest and cyclonic eddies should be further investigated and this balance may shift from the Charleston gyre to further downstream from Cape Hatteras (Andres et al., 2023).

### 4.6 Phytoplankton Community Composition

The frontal interface could represent a distinct habitat even in the highly dynamic and turbulent Gulf Stream. In particular, the obduction of nutrient-rich Sargasso Sea water could allow a distinct microbiome to quickly grow. Our limited data shows higher salinity water containing different PCC from the Gulf Stream water. We expect the entrainment of shelf filaments will lead to PCC that is similar to the shelf though it could appear as an anomaly when entrained between the Gulf Stream and a warm streamer.

In previous work, the increase in phytoplankton biomass and presence of a unique community at frontal zones has been variously attributed to lateral mixing (bringing diverse groups into the same place (Barton et al., 2010)), vertical circulation (injecting new nutrients (Clayton et al., 2014)), complementary nutrient compositions in the adjacent water masses (e.g. mixing iron-poor nitrate-rich and iron-rich nitrate-poor waters (Cassar et al., 2011; Ribalet et al., 2010)), or a combination of stirring and biotic processes (Mangolte et al., 2023). Short-lived submesoscale fronts that are common around larger geostrophic fronts can also structure phytoplankton diversity (Mousing et al., 2016), though they have unknown impacts on predator prey relationships given their ephemeral nature (Greer et al., 2015).

Will we be able to remotely sense these PCC changes across the front with upcoming 1 km resolution hyperspectral satellites such as the Plankton, Aerosol, Cloud, ocean Ecosystem (PACE) mission? In the frontal eddy scenario (Figure 5, middle column) the changes are fairly drastic and it is likely that we could distinguish different communities with PACE. In the cold filament scenario (Figure 5, left column) the frontal anomaly in gamma and PCC is very fine-scaled and would likely be missed at 1 km resolution. In the high salinity obduction scenario (Figure 5, right column) PCC shifts in a step change at the front so 1 km resolution would not miss any spatial dynamics, except the true sharpness of that shift.

### 4.7 Caveats and New Questions

#### 4.7.1 Defining and finding the front - What is a front?

While ocean fronts are often described as a “line in the sea” (Yoder et al., 1994) and do sometimes appear so, we found that with continuous data, the front can exhibit phenomenal complexity and present an entire sub-front scale that is often missed by coarser approaches or averaged representations of fronts (Figure S1). Importantly, for comparisons across observational approaches, the temperature, salinity, current, and ocean color fronts are often spatially offset, sometimes by kilometers. This appears to be largely controlled by the phase of the meander in this region. The definition of the front, whether density, current, temperature, salinity, or ocean color and an analysis based on this definition could lead to very different conclusions. It should be noted that no matter what definition(s) are used to define the front, they are not consistent through depth and their structure can change dramatically in short periods of time and space.

While it may be appropriate in some investigations, and at least a best viable option in others, it does call into question the use of satellite SST to define WBC fronts given the frequency of a 2-15 km offset between the temperature and density or current fronts. While SSH via altimetry provides a reliable metric for delineating the front (Andres, 2016; Gray & Johnston, 2021) the 0.25 degree resolution is not suitable for looking at submesoscale or fine-scale dynamics. The recently launched Surface Water and Ocean Topography (SWOT) mission will help with this substantially moving forward, improving our ability to spatially resolve ocean features by nearly 10 times (Morrow et al., 2019).

We found that surface properties are generally divergent from the properties at depth near the front and using satellite data from the top ∼1 cm of the water column in the case of longwave infrared SST, or even the first optical depth for ocean color (1/Kd, ∼10 m in our study area), may not allow proper investigations of dynamics. This has been found to be true of the vertical structure of the Gulf Stream more broadly (Meinen & Luther, 2016) and we find it to be the case for the front at finer scales.

Despite these challenges, linking PCC to the meander phase of the Gulf Stream from satellites will be substantially improved with PACE-based PCC metrics and SWOT-based physical structure of the front. The inclusion of HF radars where the Gulf Stream is close to the coast [sensu (Muglia et al., 2022)] could also help substantially to connect these processes.

#### 4.7.2 Capturing the full breadth of phytoplankton community composition

Measuring high spatial resolution PCC and differentiating phytoplankton species is methodologically challenging and expensive. Our optical flow-through measurements capture the integrated contribution of all groups to the inherent optical properties of the water and are binned into averages of ∼100 m. For most questions, this is an appropriately high spatial resolution, but the measurement technique is more variable and less discriminative than chemical, imaging-based, or molecular methods. A limitation to all of our PCC data is that it is surface only. Paired with additional techniques (e.g. continuous flow cytometry and automated imaging microscopy) we could observe a broader size range of plankton in the ocean but standardizing across methods can be challenging (Lombard et al., 2019). Meta-analyses such as that conducted by (Mangolte et al., 2023) in the California Current Ecosystem highlight a path towards addressing these challenges.

To better understand the impact of frontal dynamics on biogeochemical cycles and ecosystem diversity, we recommend future work use optical proxies and automatic imaging tools to measure PCC and nutrients across a wider size range and across depth. This could be done with a tow-yo (Hales & Takahashi, 2002) system undulating instruments across the front or a pump towed up and down the water column bringing water to instruments in the ship’s wet lab. Given the need for high spatial resolution instruments with a fast sampling rate such as the ACs, flow cytometers, and optical nitrate sensors could provide the most insight.

#### 4.7.3 Local complexities

Undoubtedly in this area, where the Gulf Stream separates from the continental shelf, the observed complexity arises from a combination of the geostrophically driven current front and interactions between this and myriad local features such as frontal eddies from upstream (Gray et al., 2023), reintegration of warm-core rings from the slope sea, major storms, movement of the Hatteras Front where the MAB and SAB waters meet (Seim et al., 2022), and outflows from the Chesapeake Bay and Albemarle-Pamlico Sound system. This study area was chosen due to the reliable location of the current front, but this local mixture of water masses makes definitively attributing chl-a or PCC changes to frontal processes more challenging. While this complexity makes this area inherently interesting - and possibly accounts for the high local biodiversity - to mechanistically understand frontal impacts on productivity and diversity an area with simpler water mass interactions is likely preferable. One possible location for such work could be further upstream in the SAB or in the Kuroshio Current. WBC fronts are not simple even in the best case though, and given nonlinear ecosystem feedbacks, extrapolating mean properties from simplified areas may not be representative of their impact on marine ecosystems.

The Ocean Observatories Initiative’s Pioneer Array will be relocated to this area in 2024 and provides an excellent opportunity to investigate these questions in more depth (Text S6).

## 5 Conclusions

In this work we investigate kilometer-scale processes, a scale that has rarely been observed, and our observations provide evidence for coherent niches and biomass enhancement at this fine-scale. We suggest that intermittent meander driven vertical motion is connecting the euphotic zone and the Sargasso Sea nutrient strata and subducting cold filaments. This work integrates and advances previous studies by 1) suggesting that the Sargasso Sea nutrient bearing strata shoals at this latitude, 2) demonstrating the front is an interface with a sub-frontal scale rather than a singularity, and 3) indicating that submesoscale processes at this interface, especially meander dynamics, can link the well-lit surface with the deeper ocean.

Ocean fronts have long interested mariners, fishers, and oceanographers for good reason. The aggregations of fish, marine mammals, and seabirds are incredible to witness and the sharpness of the interface between water masses brings some of the immensity of the ocean down to a human scale – colors and currents are hard to gauge by eye without comparison and fronts provide that rare contrast. The present study focuses on the lower end of the submesoscale and provides evidence that a range of important biogeochemical processes may be occurring here which warrant further study and will undoubtedly require advances in observational and modeling approaches.

## Supporting information

Supplemental Figure 5 (all transects visualized)

## Acknowledgments

Funding support was provided by National Aeronautics and Space Administration (NASA) Future Investigators in NASA Earth and Space Science and Technology (FINESST, #80NSSC19K1366, Ocean Biology and Biogeochemistry program) and the Zuckerman STEM Leadership Program to PCG. R/V Shearwater ship time was supported primarily by ONR-NRL program element 61153N, WU 72-1R25 to IS and partially by the Nicholas School of the Environment and Duke University Marine Lab donors through a student research grant to PCG. The authors thank the crew of the R/V *Shearwater*, Matt Dawson, Tina Thomas, Zach Swaim, and John Wilson, and the captain of F/V *Instigator*, Josh Wentling. We thank Ali Chase for helpful discussions of her gaussian decomposition algorithm for phytoplankton pigments.

Additional funding support was provided by NSF Award OCE-1829905 to SAF. We thank Minghao Li for assistance with flow cytometry sample analyses and Wake Forest University for student support provided to ML.

## Open Research

The code to recreate this analysis are available at https://doi.org/10.5281/zenodo.8377865 (Gray, 2023a) and all data used in this study is available at https://doi.org/10.5281/zenodo.8377770 (Gray, 2023b). All code is shared with an MIT License for free reuse. We have provided multiple Jupyter Notebooks that go from raw or initially processed data to the nearly complete figures shown in the paper. The python environment used can be easily and exactly reproduced using the pangeo-notebook a Docker image https://github.com/pangeo-data/pangeo-docker-images/tree/master/pangeo-notebook as detailed in the Github repo.

## Notes

https://doi.org/10.5281/zenodo.8377770

https://doi.org/10.5281/zenodo.8377865

### Competing Interest Statement

The authors have declared no competing interest.

