## Supplemental Text and Figures for "Evidence for kilometer-scale biophysical features at the Gulf Stream front"

*JGR: Oceans*

### Contents of this file

Text S1 to S6
Figures S1 to S5
Table S1

### Additional Supporting Information (Files uploaded separately)

Figure S6

### Introduction

This supporting material covers extended methods for processing optical flow-through, nutrient, and flow cytometry data. It includes additional discussion of nutrient results, phytoplankton community composition results, and a section on future work. Figures include demonstrations of scale, a 2D view of the water column from Figure 5, the physical environment via a T-S diagram, and a detailed look at the nutrient and flow cytometry data. Finally included is an additional file with visualizations of every cruise transect used in this study.

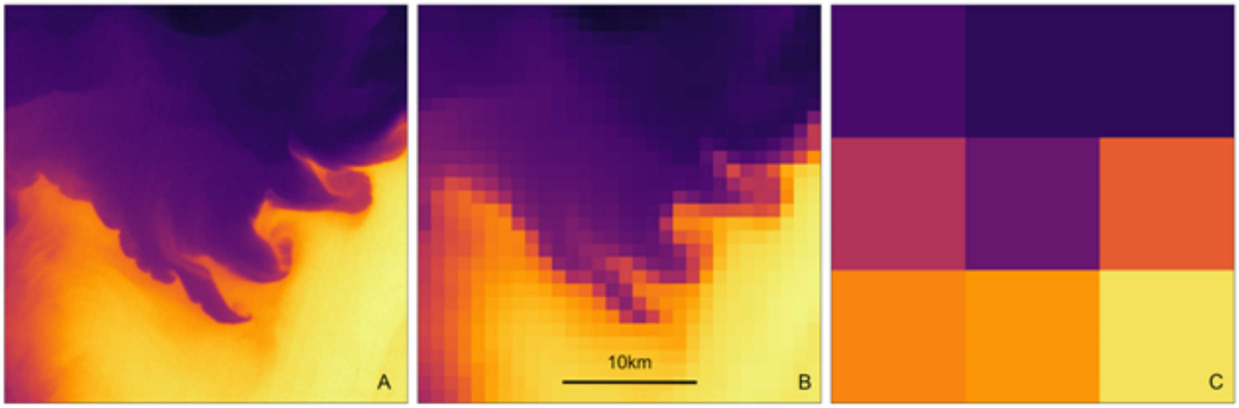


**Figure S1.** Thermal imagery from Landsat 8 off Cape Lookout, North Carolina, USA, showing A) 100m Landsat imagery of the Gulf Stream front and an array of submesoscale filaments and eddies, B) imagery resampled to 1km representing the typical ocean observing satellite resolution, and C) 10km imagery representing the typical "high resolution" global or basin scale biogeochemical model resolution.

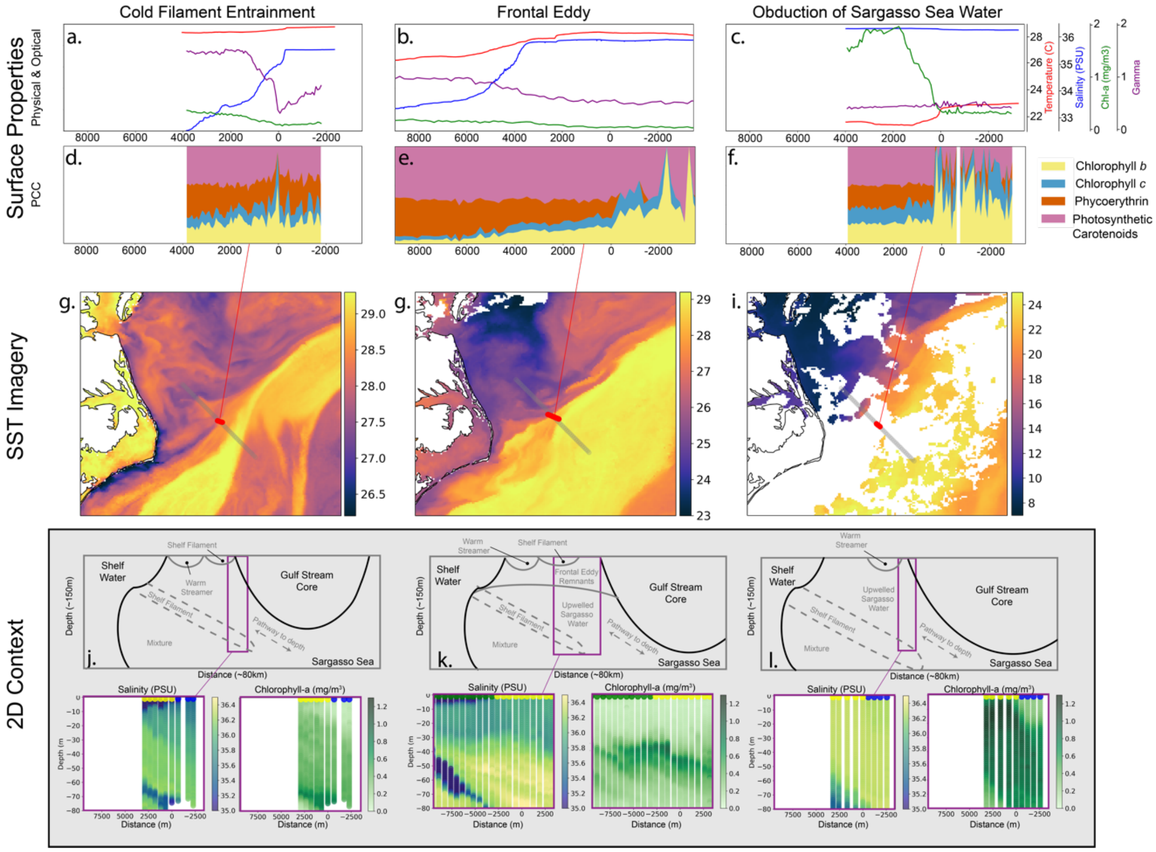


**Figure S2**. More detailed version of Figure 5. Surface phytoplankton community composition (PCC) during three different processes on the front. Surface properties are shown on the top row. Satellite SST is shown in the middle row. Bottom row shows the 2D context both in the form of vertical profiles and a schematic with solid lines indicating isotherms. The left column is crossing a cold filament and into the Gulf Stream, in the middle column we cross from a shelf filament into a frontal eddy, and on the right column we cross from a warm streamer into upwelled Sargasso Sea water and finally into Gulf Stream water.

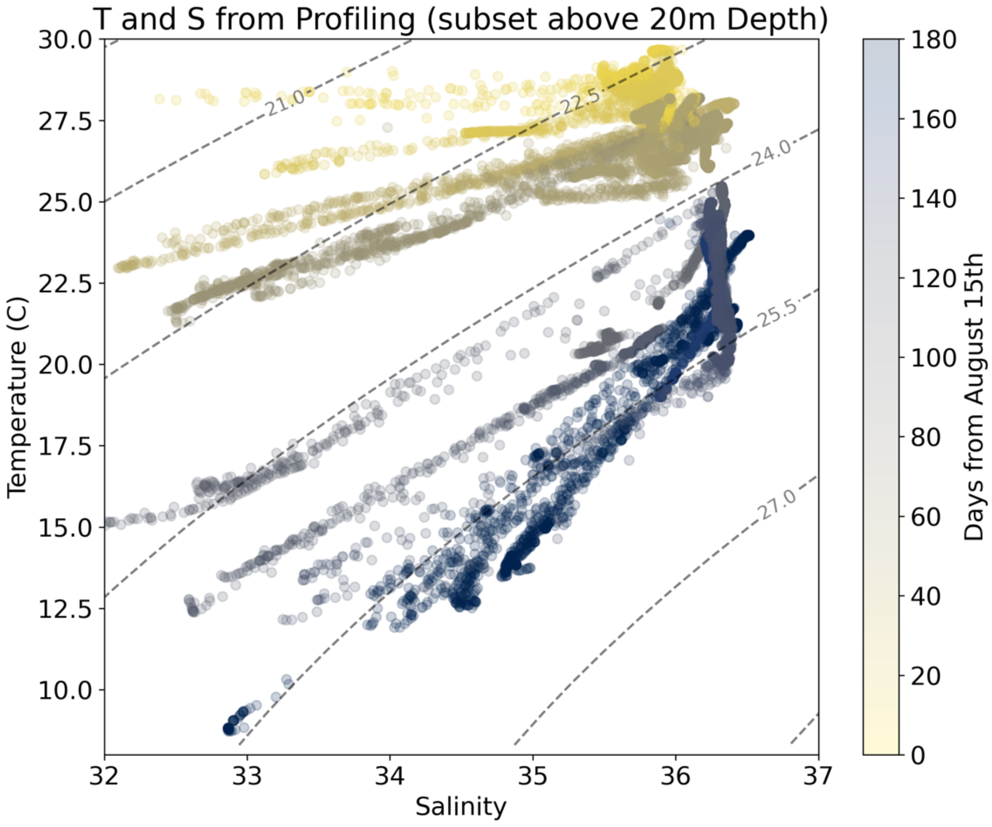


**Figure S3.** Temperature and salinity diagram showing surface data (top 20m) from the VMP-250 colored by the days between the sample and August 15$^{th}$. This shows the slight seasonal dependence of Gulf Stream water salinity (highest salinity in winter) and the more extreme seasonal temperature changes in both Gulf Stream and coastal water.

**Text S1. Flow-through Optical Properties Methods Continued**

For each minute of the total seawater measurement, the signal between the 2.5th and 97.5th percentiles were averaged, and their standard deviation was used to quantify uncertainty. Dropping the 2.5th to 97.5th percentiles filters out noisy spikes from bubbles which can be a major problem in optical measurements. All spectra were manually checked and quality controlled for obviously bad measurements (e.g. bubbles, bad filtered seawater measurements). All $a_p$ and $c_p$ spectra were unsmoothed following the method in (Chase et al., 2013) and a residual temperature and scattering correction (Slade et al., 2010) was applied using the temperature and salinity dependencies from (Sullivan et al., 2006). Following (Sullivan et al., 2013), a $\chi$=1.097 is used to convert from measurement of scattering around a single angle (nominal angle 120°) to the particulate backscattering coefficient ($b_{bp}$).

**Table S1**. Optical measurements and biogeochemical properties they are proxies for.

| Parameter | Definition | Proxy |
|---|---|---|
| Absorption, $a_p(\lambda)$ | Absorption coefficient of light due to particles | Total pigment concentration |
| Scattering, $b_p(\lambda)$ | Scattering coefficient due to particles | Total particle composition and concentration |
| Attenuation, $c_p(\lambda)$ | Extinction of light due to particles (Absorption + Scattering) | Total particle concentration |
| Backscattering, $b_{bp}(\lambda)$ | Scattering in the backward direction due to particles (>90 degrees) | Total particle composition and concentration |
| Backscattering Ratio, $ƀ_p(\lambda)$ | Particulate backscattering / particulate scattering | Refractive index (composition) and size |
| Chlorophyll-a line height | Chl-a estimate based on the absorption line height at 676nm | Chl-a |
| Gamma, $\gamma$ | Spectral slope of the particulate beam attenuation | Mean particle size |
| Gaussian decomposition of absorption | Decomposition of absorption spectra into Gaussian components | Concentration of accessory pigment, proxies of phytoplankton groups. |

**Text S2. Nutrient Methods**

March 2022 nutrients were collected by filtering duplicate ~10 mL through 0.22 $\mu$m PES membrane Acrodisc filters and the filtrate was stored at 4° C until analysis was facilitated at Bigelow Laboratory for Ocean Sciences (https://www.bigelow.org/services/nutrient-analysis/) where a Seal Analytical continuous-flow AutoAnalyzer-3 was used to measure silicate, nitrate, nitrite, phosphate, and ammonia. September 2021 nutrients were collected by filtering duplicate ~50mL samples through 0.22 $\mu$m Sterivex filters and the filtrate was stored frozen at -80° C until analysis at the UCSD Nutrient Analysis facility (https://scripps.ucsd.edu/ships/shipboard-technical-support/odf/chemistry-services/nutrients) where a Seal Analytical continuous-flow AutoAnalyzer-3 was used to measure silicate, nitrate, nitrite, phosphate, and ammonia.

**Text S3. Flow cytometry (FCM) Methods**

Duplicate whole seawater samples were collected from the ship's flow through system fixed with 0.125% glutaraldehyde and stored at -80° C until processing. Summer prokaryotic and eukaryotic phytoplankton populations were enumerated using a Becton Dickinson FACSCalibur Flow Cytometer, while winter populations were enumerated using a Beckman Coulter CytoFLEX Flow Cytometer (Johnson et al., 2010). Summer bacterioplankton were quantified using SYBR Green-I on the Attune NxT acoustic flow cytometer (Life Technologies).

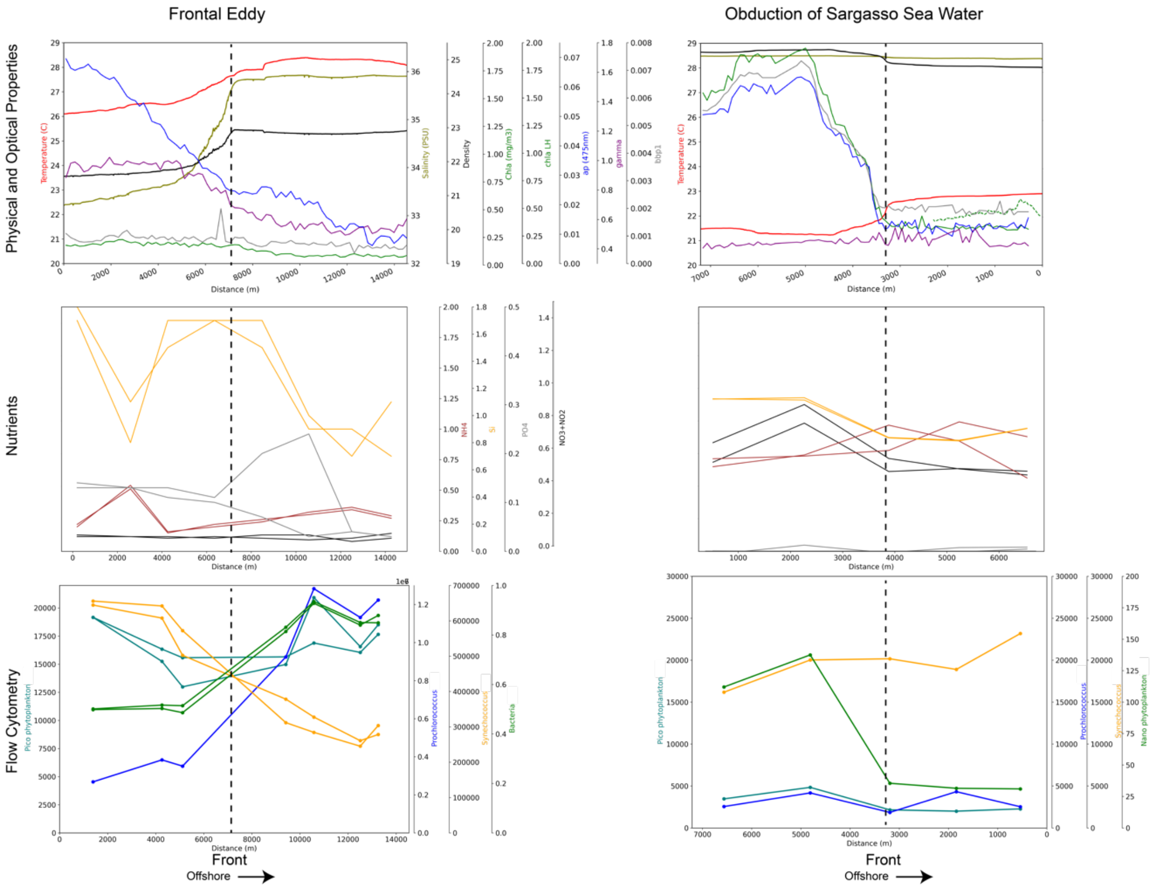


**Figure S4**. Comparison of optical, physical, chemical, and flow cytometric properties during a period with obduction of sargasso sea water and during a frontal eddy. The top row shows physical and optical properties, the second row nutrients, and the third row flow cytometry.

**Text S4. Nutrient Discussion**

It was not expected at the beginning of this work that we would see phosphate limitation this close to shore or in the Gulf Stream. Given that we only have nutrient data over one cruise in September and one in March we cannot fully determine if this nutrient limitation has a seasonal pattern, but that is our initial intuition with N limiting in the summer and P limiting in winter. Though we cannot be confident without nutrient addition experiments and more data. Based on the Redfield ratio, P is limiting in the Sargasso Sea (Wu et al. 2000) though work at this latitude in the core of the Sargasso Sea using bioassays suggests N is limiting for some species (Prochlorococcus) while N and P are co-limiting for others (Synechococcus and eukaryotic cells) (Moore et al., 2008). At the edges of the gyre and beyond, N is expected to be limiting, although the Gulf Stream Core is not expected to be limited in N and may often be limited by light rather than nutrients (Palter et al., 2011). Thus, we speculate seasonally enhanced mixing in the winter is bringing up more N compared to P, leading to a P limitation. N-fixation may play a role as well. More recent work has indicated that while N is the primary limitation, P does have a secondary co-limitation in the MAB during the summer, with atmospheric deposition possibly playing a role in enhancing growth (Sedwick et al., 2018).

Interestingly, the front is not visible in the nutrient data at the surface (Figure S4) and generally the optical data reveals a much smoother change than the physical data, possibly a result of the lag time in growth from any injections of nutrients. While unexpected, and not convincingly shown in our data, it is also possible that the front here leads to some enhancements because of complementary nutrient profiles. If the coastal water is nitrate limited and the Gulf Stream water is phosphate limited or vice versa, the enhancement could be entirely from a mixture of these nutrient compositions and communities with different nutrient limitations.

**Text S5. Potential Export from Cascading Events**

We see a persistent cold and fresh filament at depth in many of our transects (e.g. Figure S2) that does not appear to be related to the meander cycle and could be a cascade of shelf or even estuarine water forming yet another pathway for high POC water to be exported in this region (Shapiro, 2003). This cascading could be driven by atmospheric cooling (Ivanov et al., 2004) or a pathway for the nearby estuarine Chesapeake Bay or Pamlico Sound water. The first report of such a cascading event in this region is described in (Han et al., 2021) which hypothesizes that it stems from atmospheric cooling increasing the shelf water density. This higher density water then flows down the local sloping bathymetry across isopycnals until it becomes neutrally buoyant and becomes entrained into the Gulf Stream flow. Han et al (2021) suggest this occurs regularly at the Hatteras Front and given our ADCP measurements showing these features moving nearly with the Gulf Stream, we may be seeing them in this last stage - neutral buoyancy and entrainment into the Stream. This adds some context to previous work suggesting convergence of Mid-Atlantic Bight (MAB) and SAB waters at the Hatteras Front leads to export of productive shelf waters (Todd, 2020), though is different from the meander driven entrainment we focus on in this study.

**Text S6. Future Work**

The framework here of meander driven entrainment and detrainment controlling the frontal interface water mass origin and properties should provide a guide to future work in the region. Frontal eddies are a special case of the meander trough and should not be ignored as we strive towards a unified understanding of WBC biophysical dynamics. Seasonality should be further inspected, and this will likely be feasible with the Pioneer Array being relocated nearby in 2024. Cruises and airborne surveys can target a specific meander phase and allow this submesoscale context to guide deeper dives into finer resolution spatial, temporal, or community structure.

Lags in response to forcings challenge nearly all aspects of marine biophysical inquiry and this is exacerbated in WBCs with flow speeds >2.5m/s and related instabilities. As stated in Clayton et al 2023 (in https://www.us-ocb.org/wbc-series-fine-scale-biophysical-controls-on-nutrient-supply-phytoplankton-community-structure-and-carbon-export-in-western-boundary-current-regions/) measuring biological rates instead of standing stocks would help with process understanding. Clever approaches to measuring rates will need to be devised (e.g. Flow cytometry (FCM)) to estimate cell division rates *in situ* (Hunter-Cevera et al., 2016)) and implemented on autonomous platforms like BGC Argo or saildrones. The lack of bioacoustic work connected to these platforms and in relation to submesoscale biophysical interactions leaves much to be desired and will allow insight into high trophic levels. In this work we saw clear structuring of acoustic backscatter in relation to the front, but instrument issues prevented this from being a large component of the analysis.

High spatial resolution sequencing (Clayton et al., 2017; Gronniger et al., 2023) paired with nearly continuous measurements of optical-based PCC will permit better understanding and regular validation of optical data. The ability to rapidly sample nutrients would be helpful, possibly via optical nitrate sensors on a similar tow-yo system.

Lagrangian approaches are critical to measuring rates and observing community evolution and grazing over time. Yet it is extremely challenging in this environment to implement a Lagrangian approach given the rapid entrainment and detrainment of water masses and diapycnal mixing. Following a frontal eddy or cold filament specifically could help illuminate processes specific to these features. The high resolution surface current data (Muglia et al., 2022) will be critical for future work and we strongly suggest the Pioneer Array incorporate HF radar into its monitoring plans.

Winds can rapidly increase turbulent mixing (D'Asaro et al., 2011) and we did not connect our study to local winds due to the two nearby NOAA buoys being out for repairs during the bulk of this work. Given the evidence for the subsurface nutrient strata so near the surface, diapycnal mixing due wind events is likely another source of patchy nutrient injections in this region.

When Gulf Stream water is covered by 5m of fresher water the frontal processes may be entirely absent from surface measurements and obscured in above-water radiometry - also during the summer the main area of productivity is at depth > 40m. The complex vertical structure here indicates the importance of LiDAR for understanding fronts where just the top 50m can be extremely informative. These extensive *in situ* results can help contextualize recent work showing intermittent chl-a enhancement at fronts (Haëck et al., 2023; Mangolte et al., 2022).

Longer time series such as Oleander could be used to verify this across time and satellite based proxies for meander phase could be derived to quantify the total impact of these processes and if there is any change in meander dynamics.

We see compelling evidence from a large dataset that both subduction and obduction are occurring at the front, with major local biogeochemical implications, though the full extent is not quantified. While we expect these results may generalize to other WBCs it is unclear how well they will translate to the much larger and more extensive field of ephemeral fronts spanning the global ocean. In other WBCs the impacts will likely depend on the water column stratification and nutrient profiles of the isopycnals the front is allowing to outcrop.

**Figure S5**. All transects used from our two years of sampling are detailed in the attached PDF. This includes SST imagery for large scale context, all profiling data, and ADCP data.
