## Supplemental Figure 5 (all transects visualized) for "Evidence for kilometer-scale biophysical features at the Gulf Stream front"

### All transects used in this analysis begin here:

```
5
data/all_adcp/processed/101706_20201010T141504UTC_100s
Transect does not cross into coastal water
```

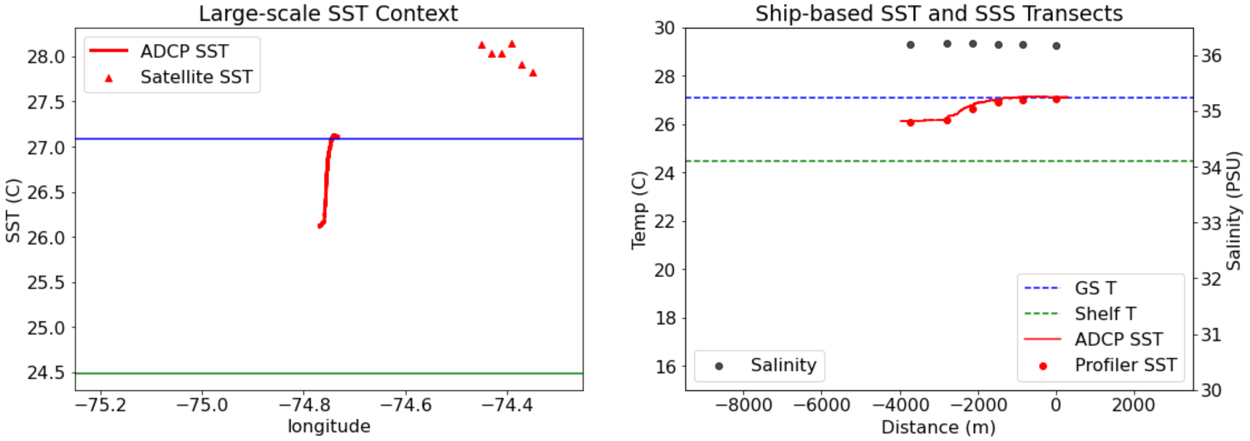

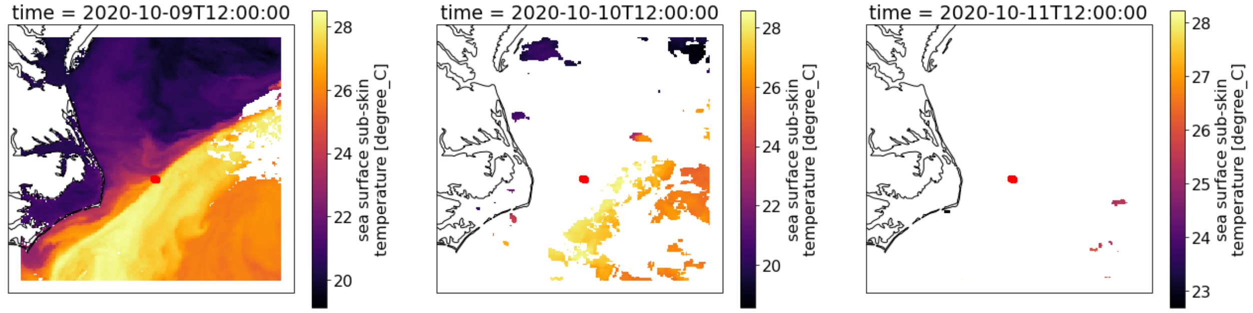

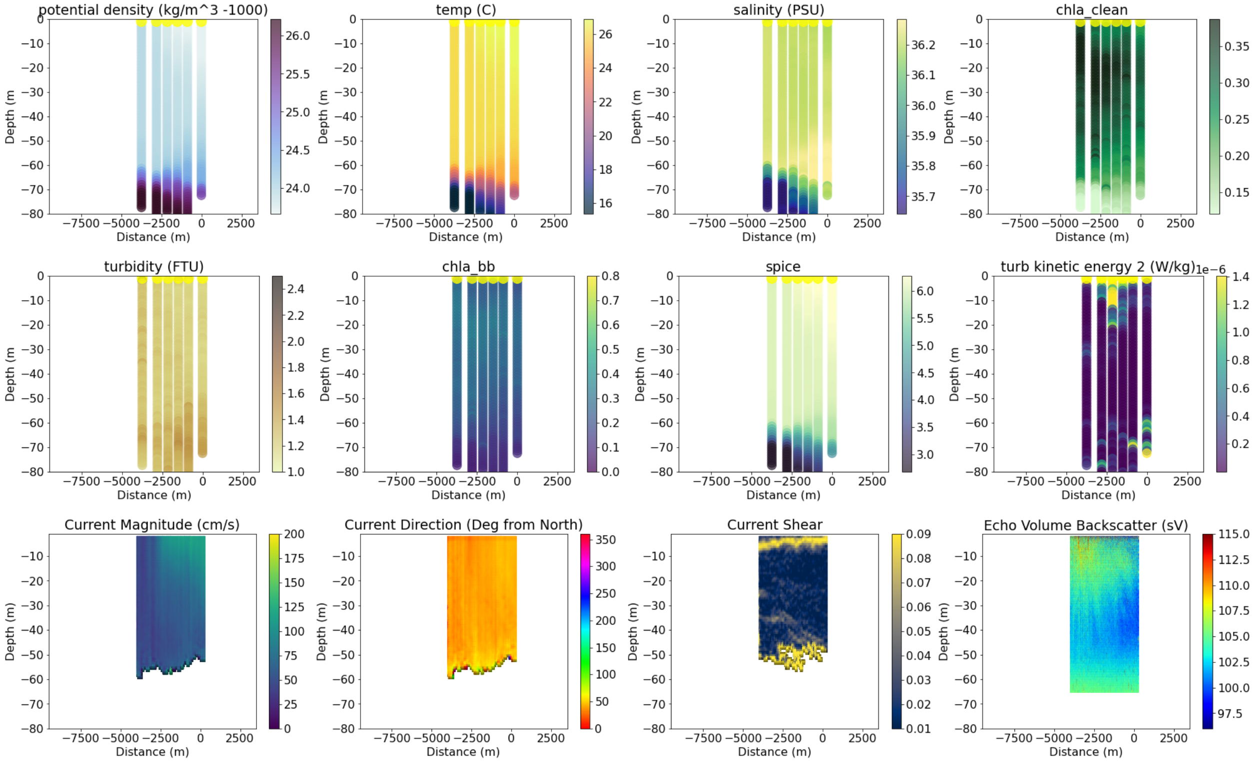

```
----------------------------------------
6
data/all_adcp/processed/101706_20201011T135347UTC_100s
Transect does not cross into coastal water
```

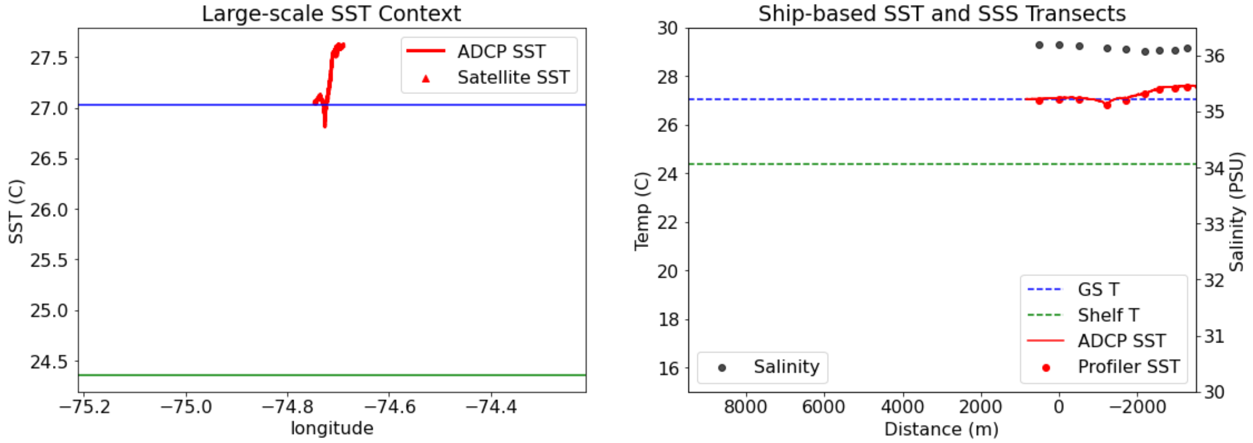

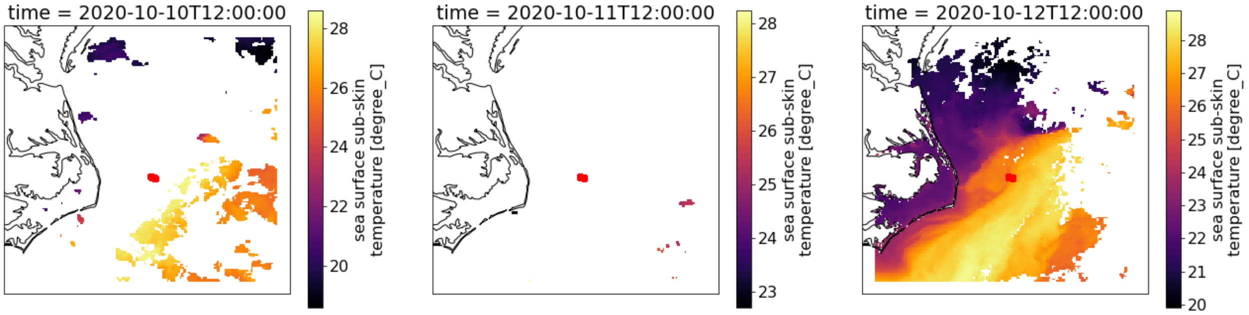


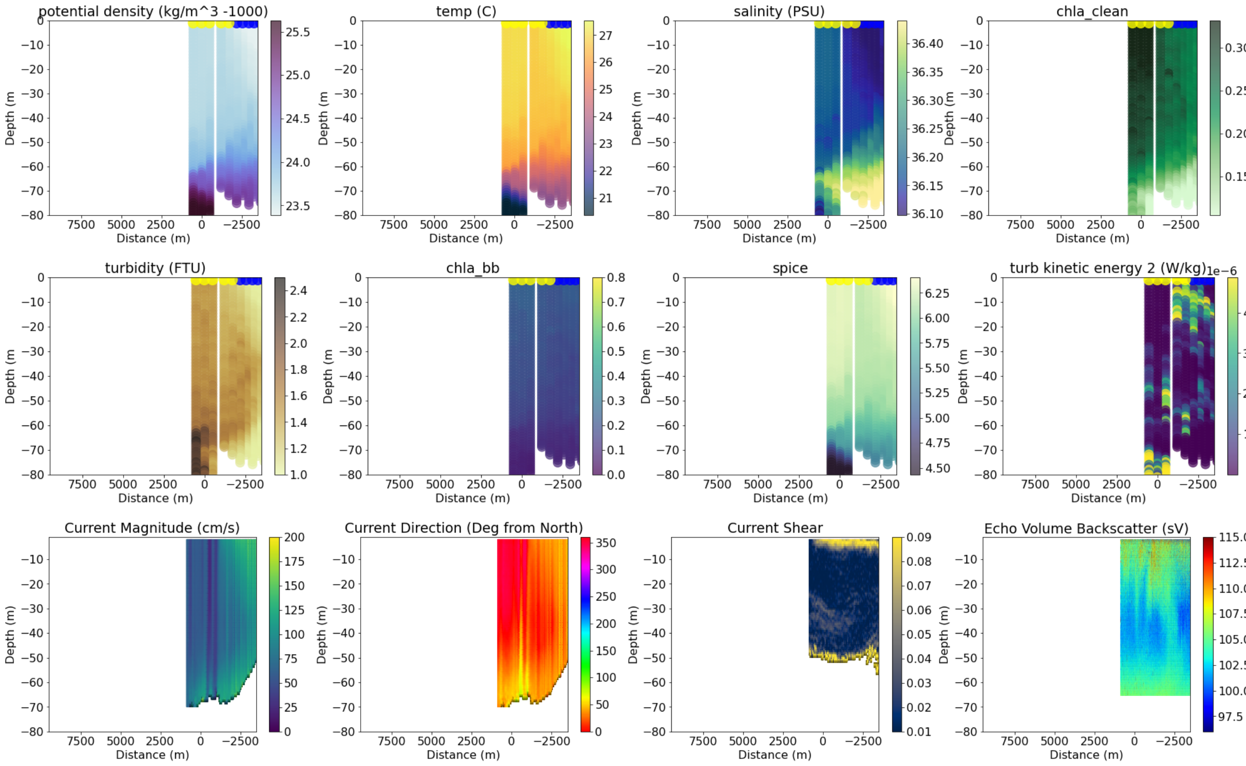


-----------------------------------
7
data/all_adcp/processed/101706_20201015T142907UTC_100s
Distance b/w GS and Coastal water is 5728.623147439484

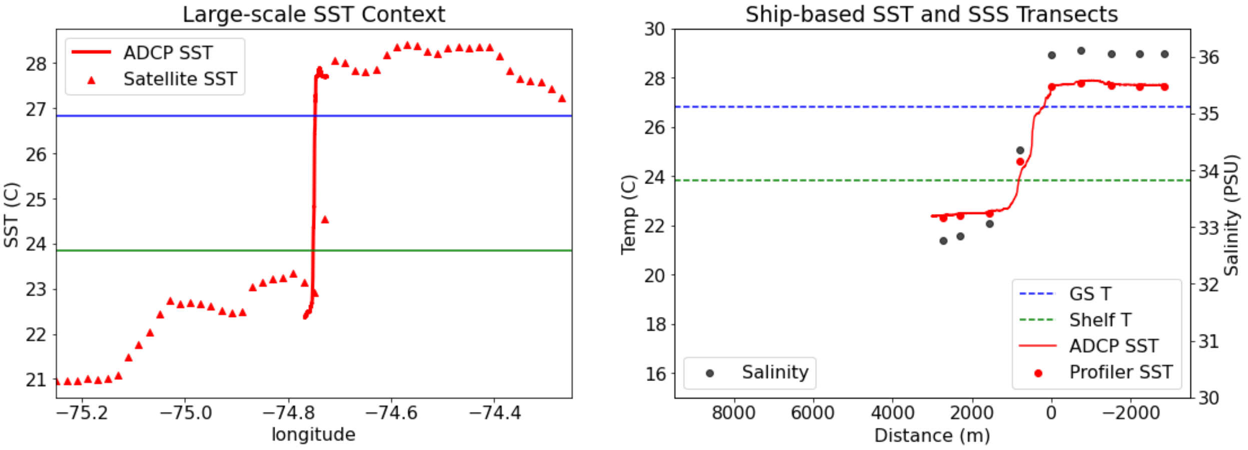

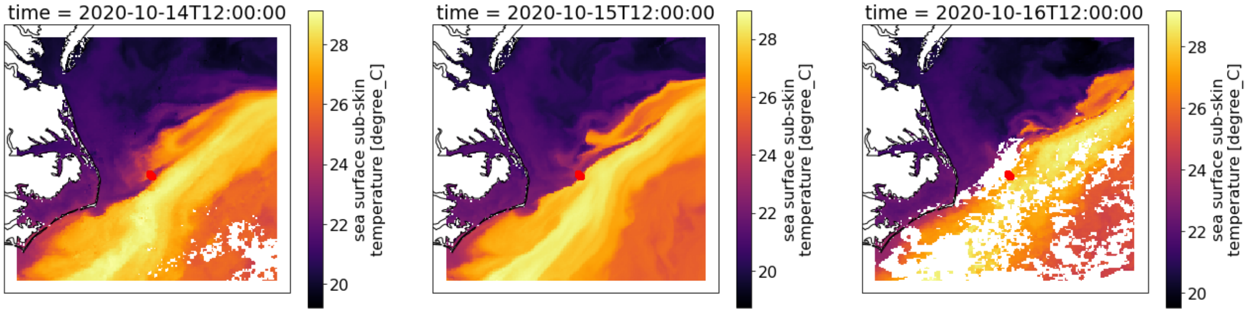


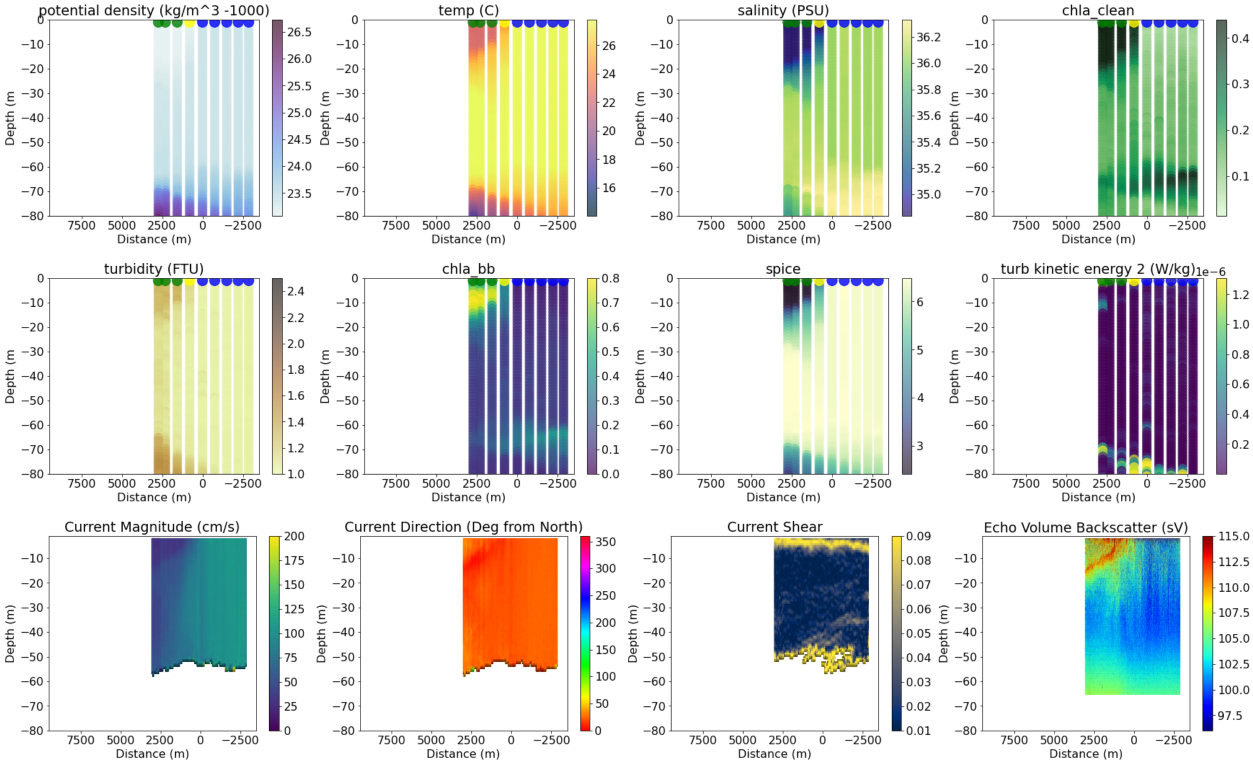


```
----------------------------------------
8
data/all_adcp/processed/101706_20201015T164414UTC_100s
Distance b/w GS and Coastal water is 2864.363522551168
```

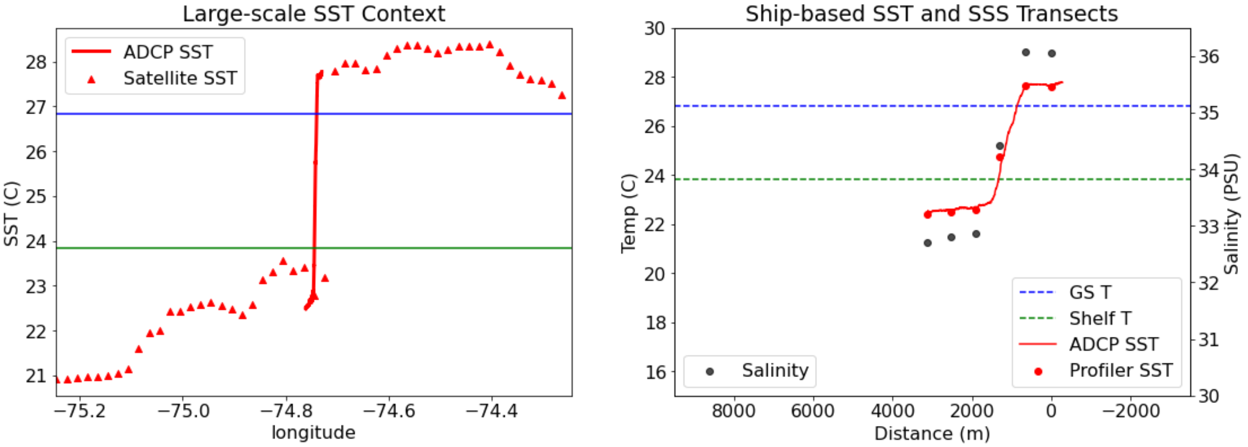

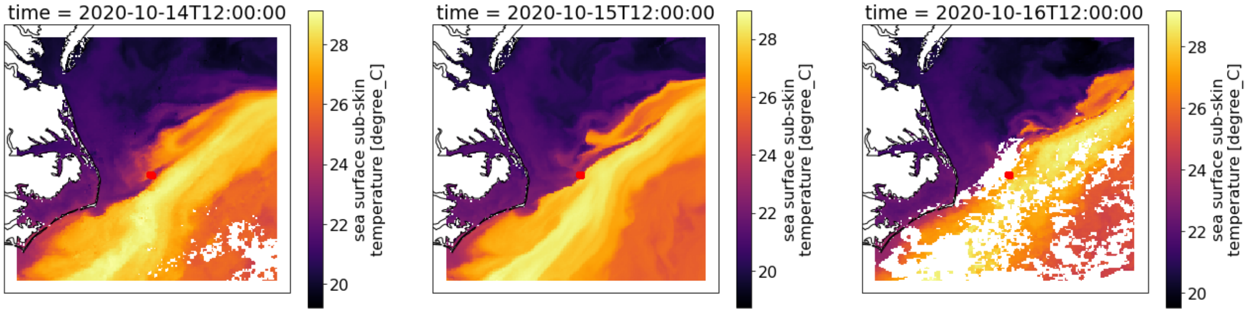


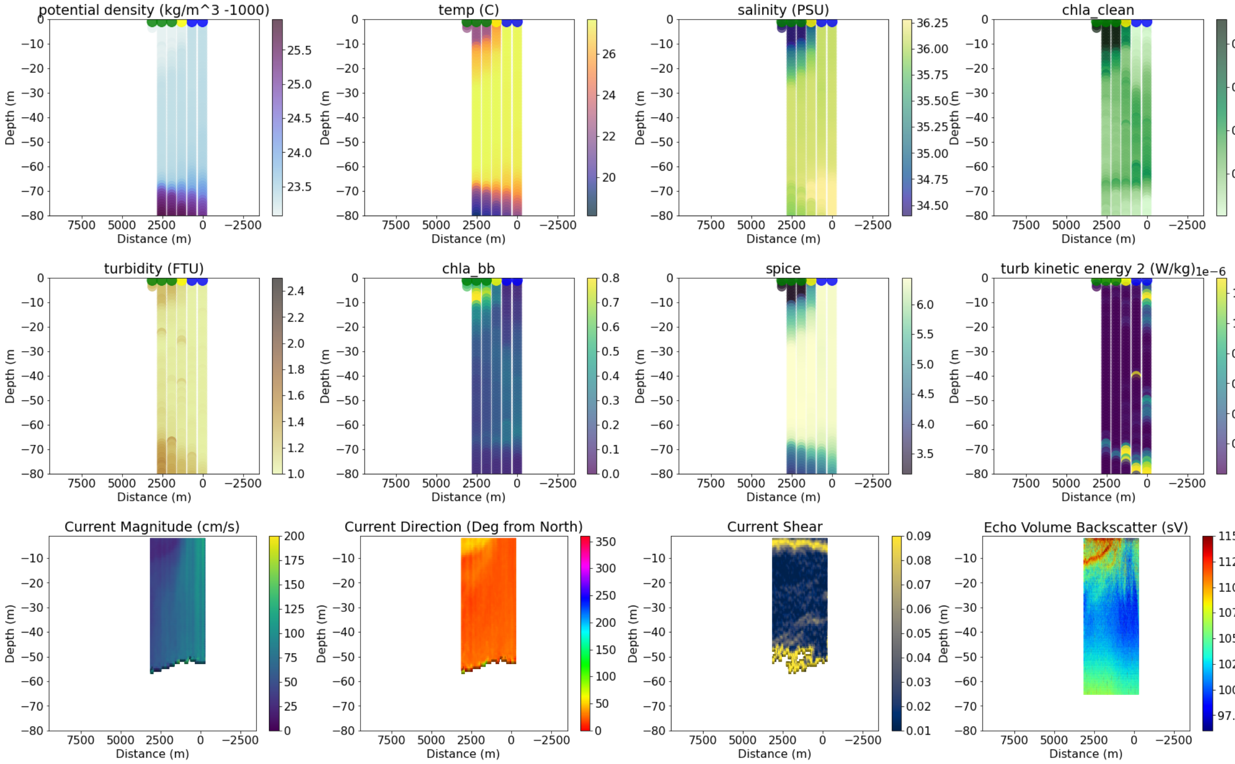


```
----------------------------------------
9
data/all_adcp/processed/101706_20201016T130156UTC_100s
Distance b/w GS and Coastal water is 22900.737788582464
```

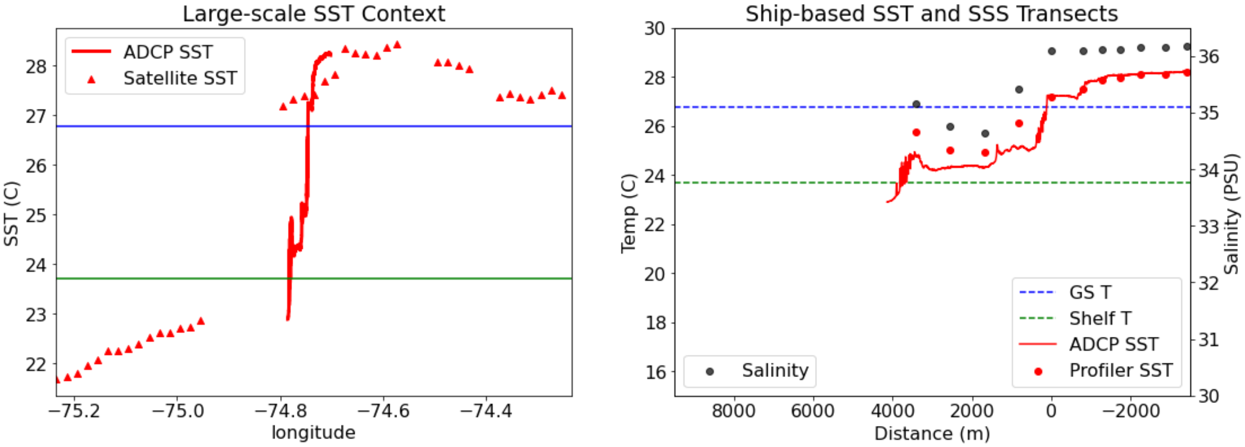

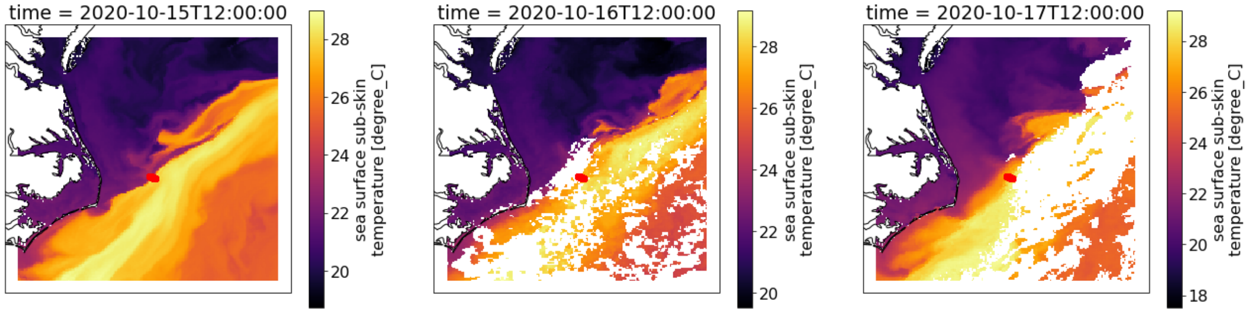


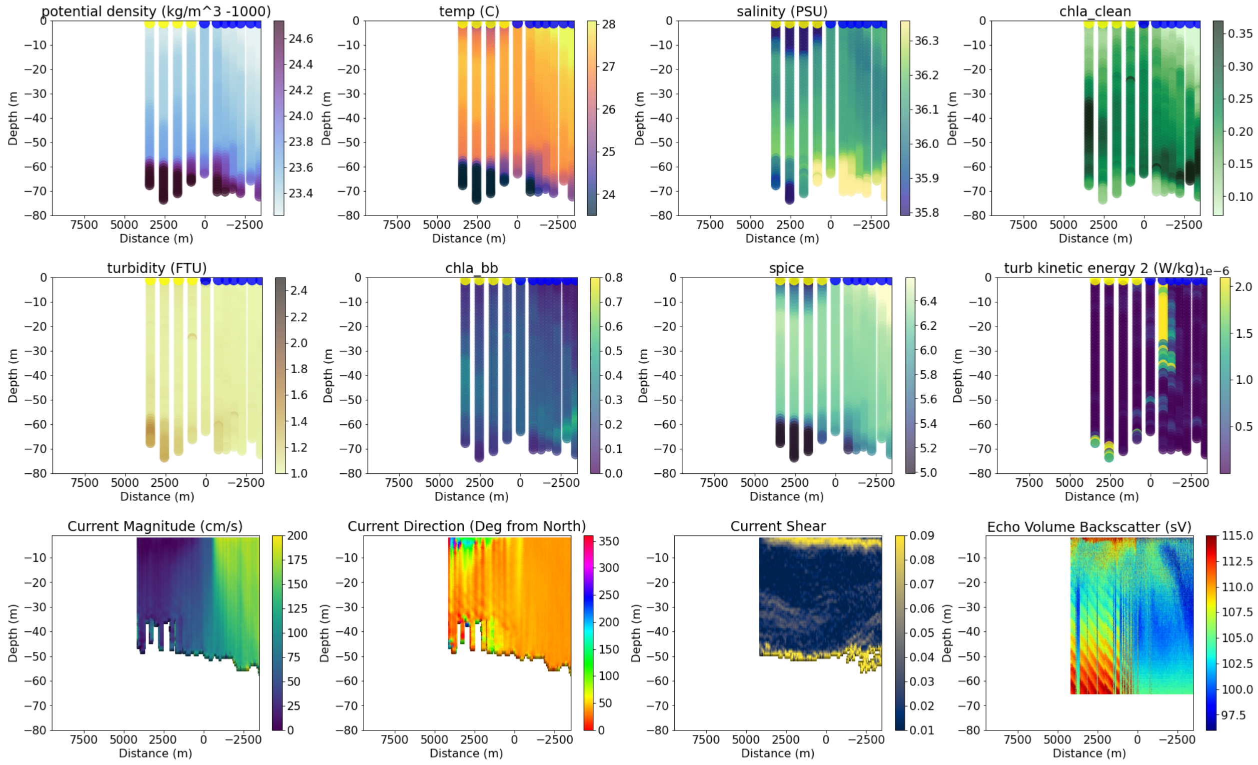


```
-------------------------------------
10
data/all_adcp/processed/101706_20201016T161245UTC_100s
Distance b/w GS and Coastal water is 25762.30420162812
```

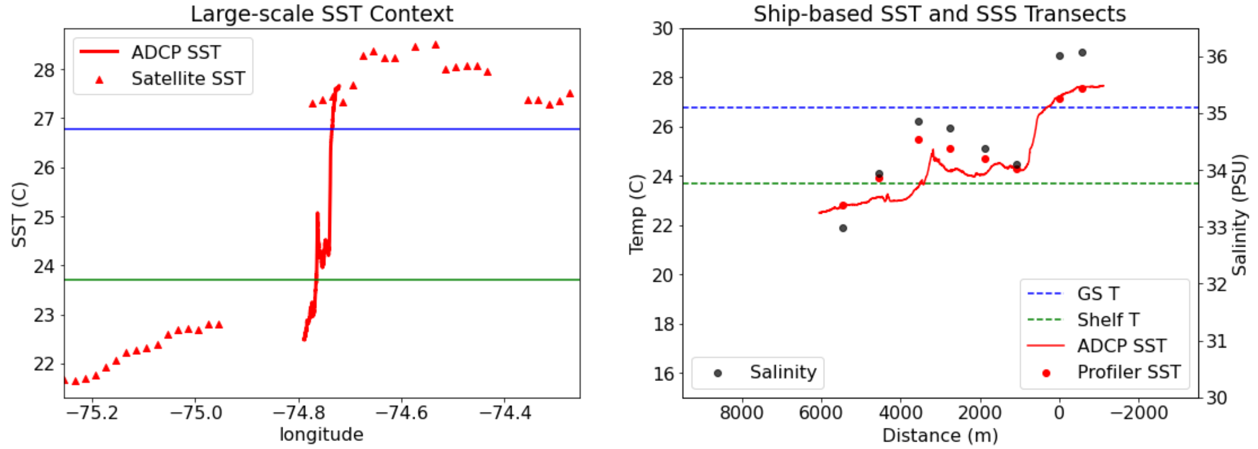

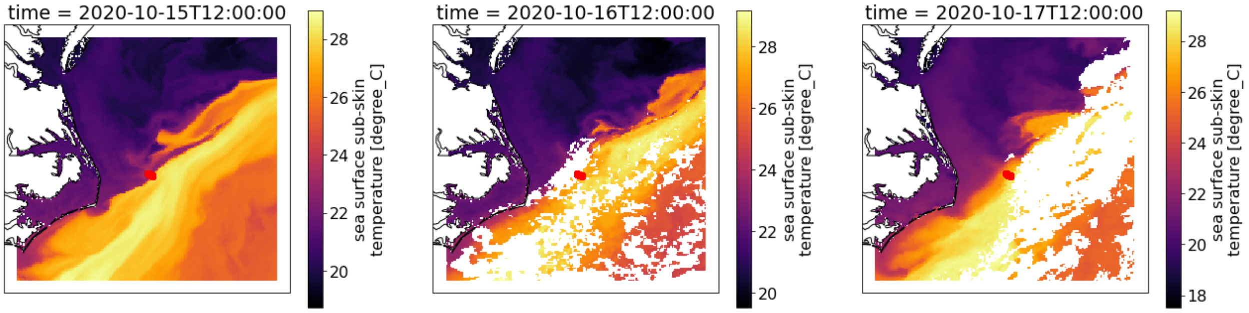


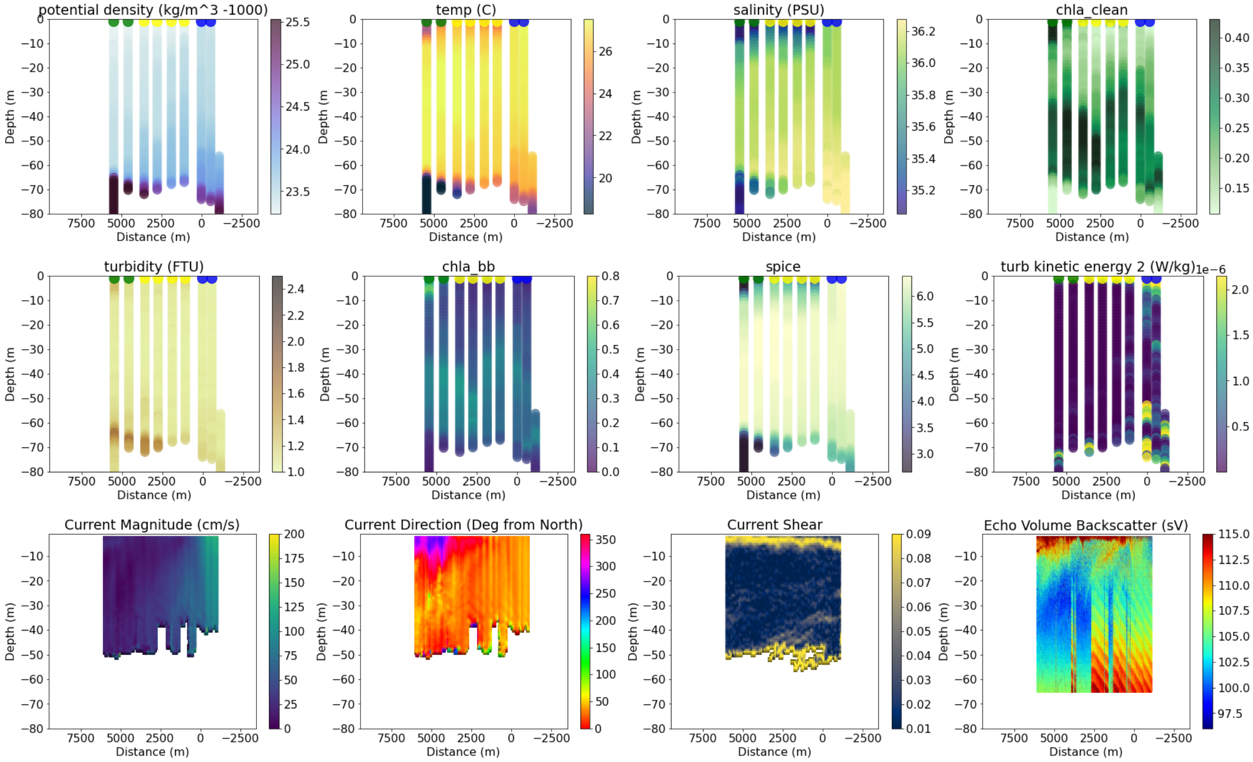


```
-------------------------------------
12
data/all_adcp/processed/101706_20201020T133400UTC_100s
Distance b/w GS and Coastal water is 8598.709764931029
```

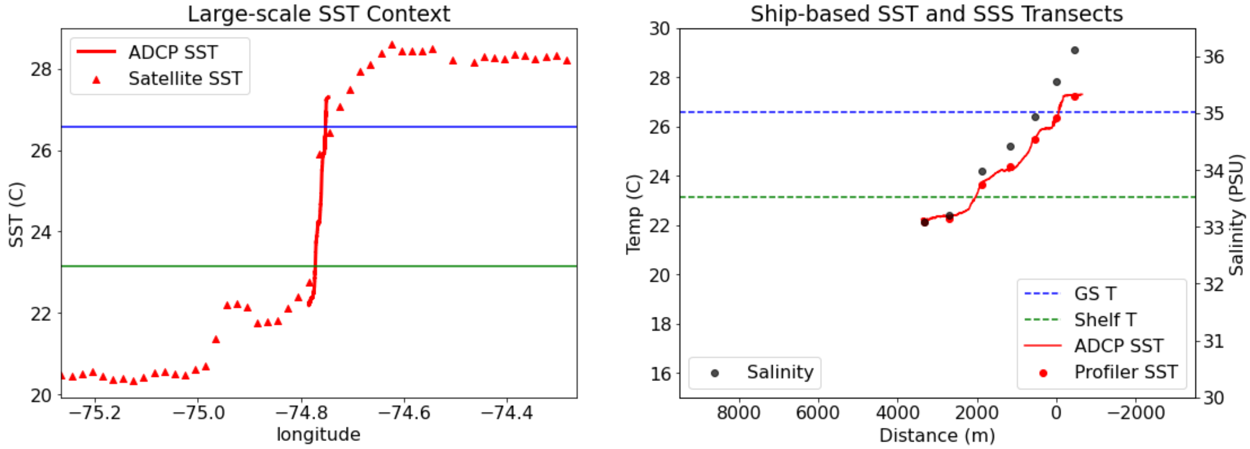

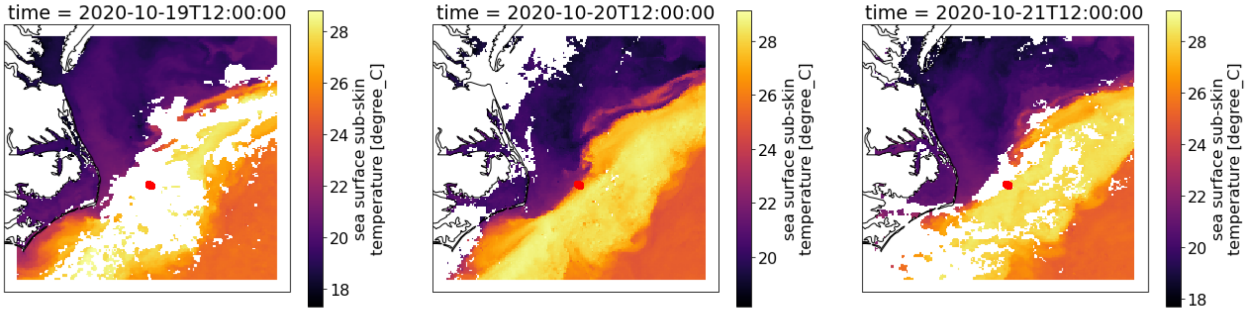


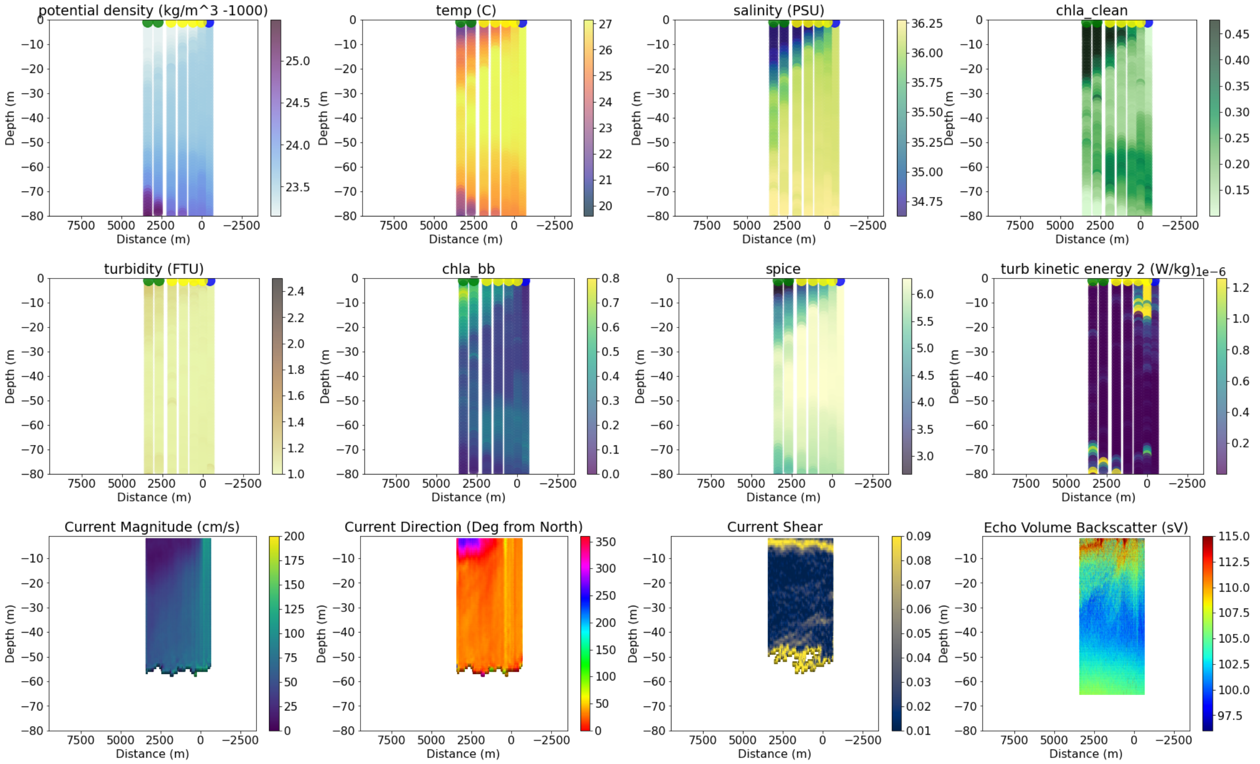


```
--------------------------------------
13
data/all_adcp/processed/101706_20201020T155242UTC_100s
Distance b/w GS and Coastal water is 5731.633277711782
```

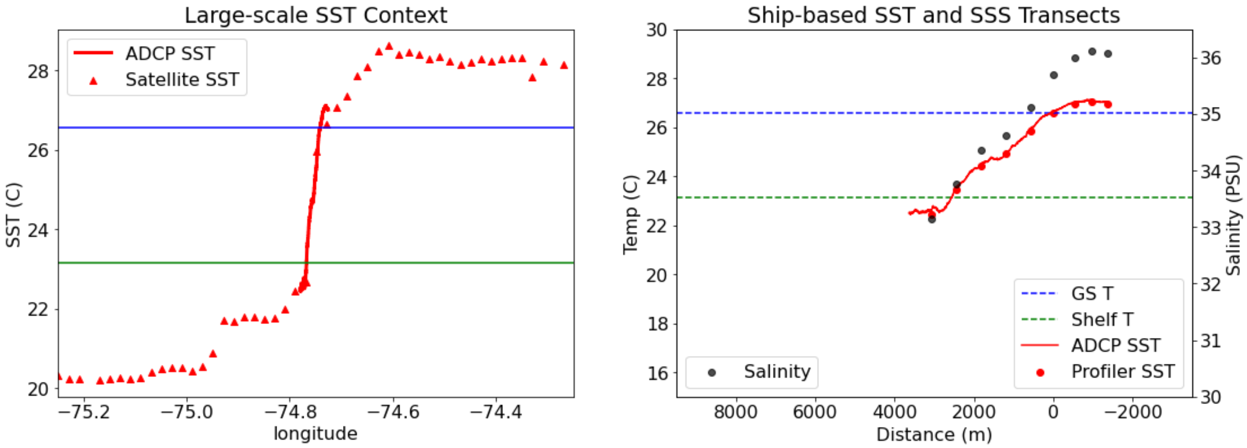

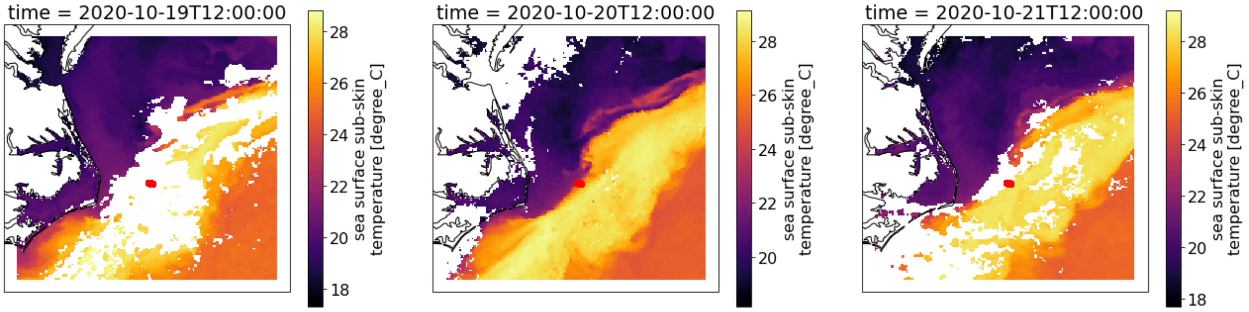


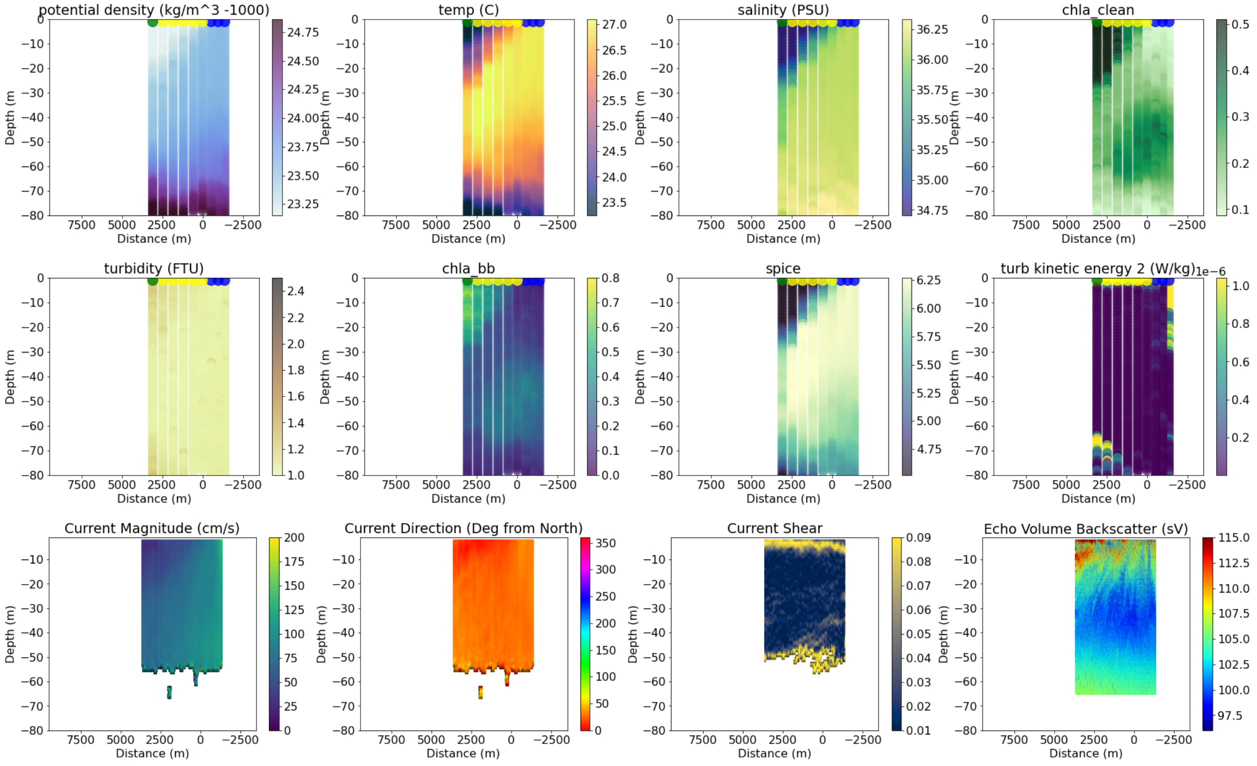


```
-------------------------------------
16
data/all_adcp/processed/101706_20210420T123333UTC_100s
Distance b/w GS and Coastal water is 28670.065325534604
```

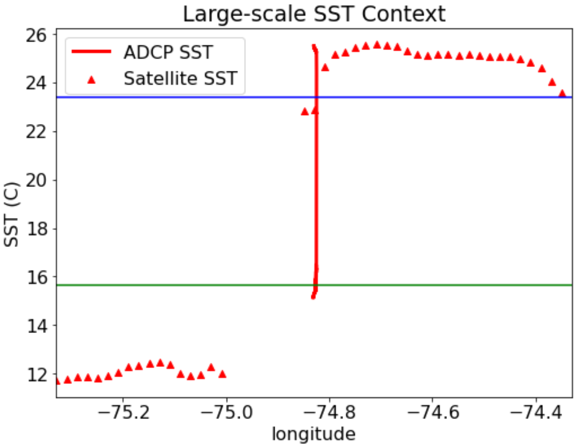


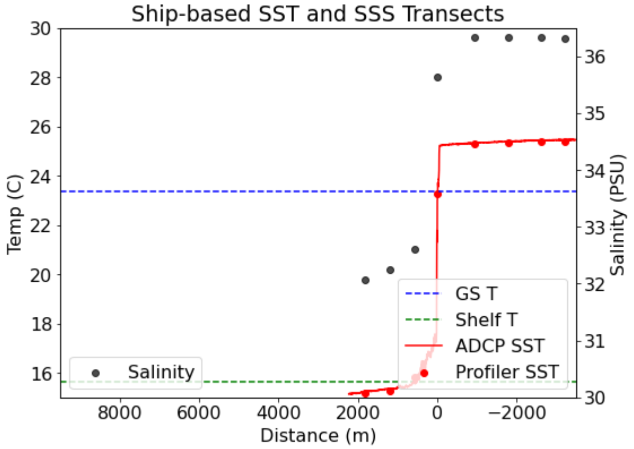

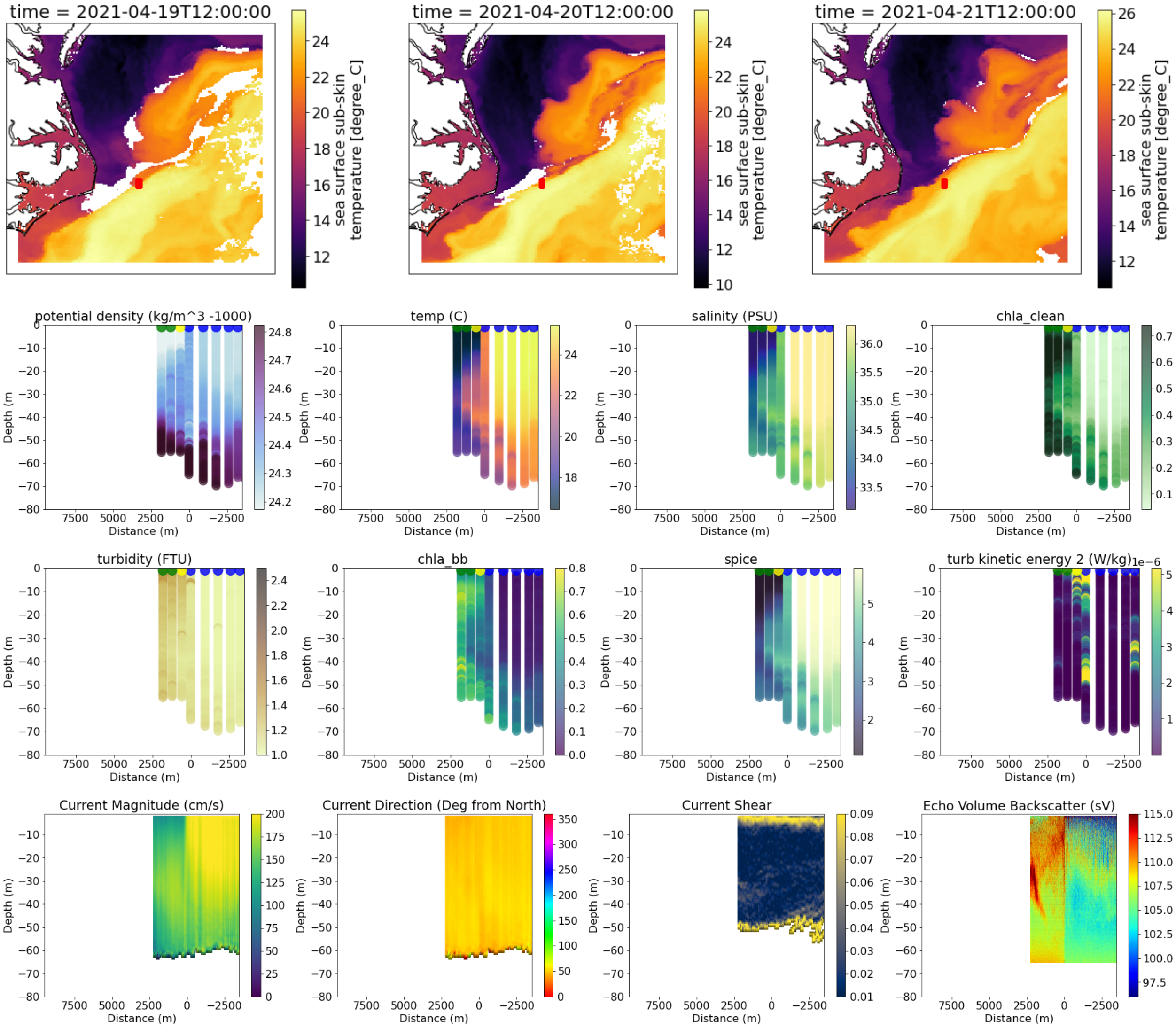


```
-------------------------------------
17
data/all_adcp/processed/101706_20210420T151905UTC_100s
Distance b/w GS and Coastal water is 25795.522432565424
```

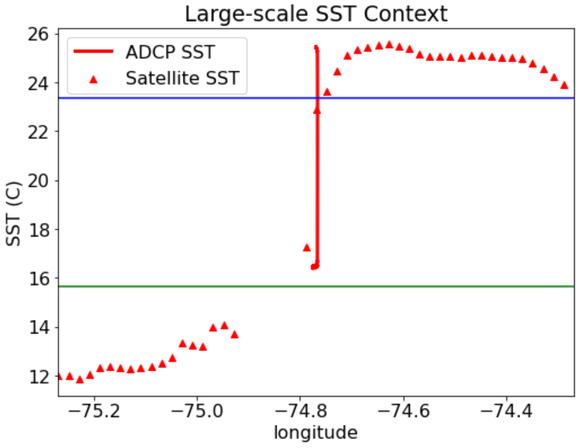


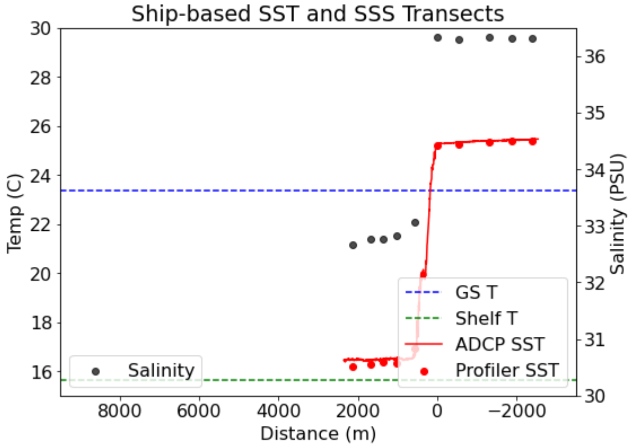

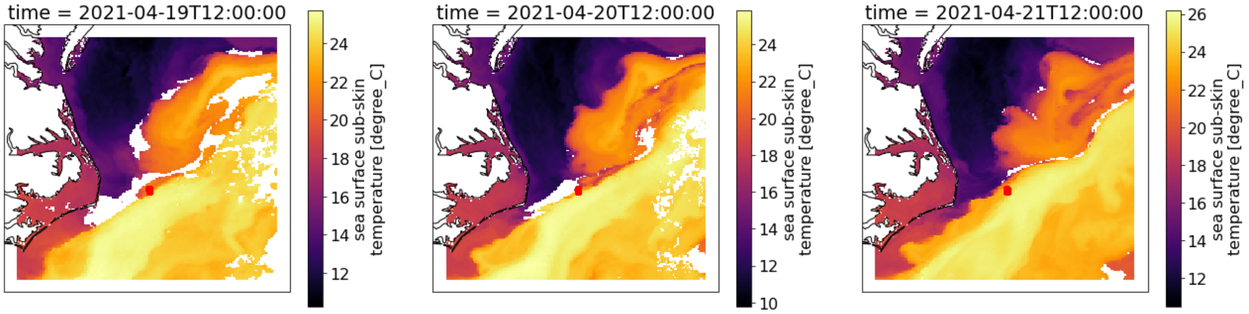


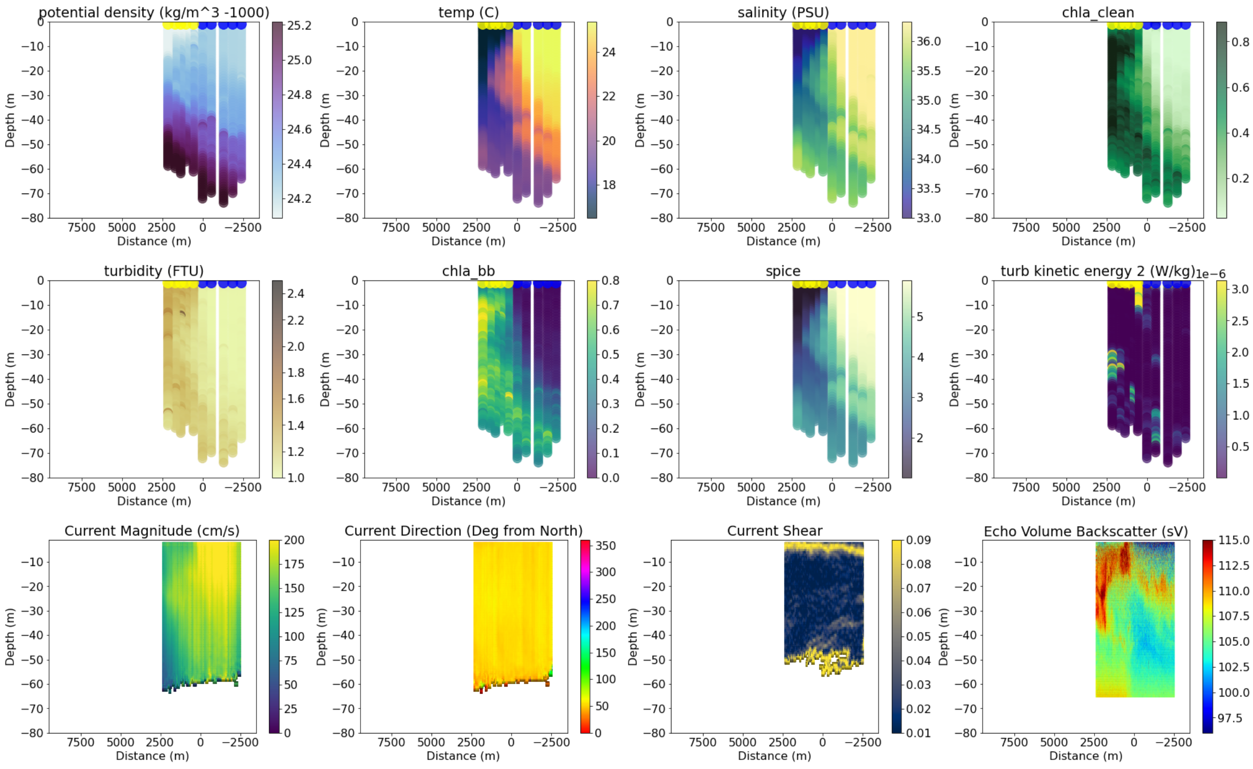


```
-----------------------------------------
19
data/all_adcp/processed/101706_20210421T150145UTC_100s
Distance b/w GS and Coastal water is 14330.036313650391
```

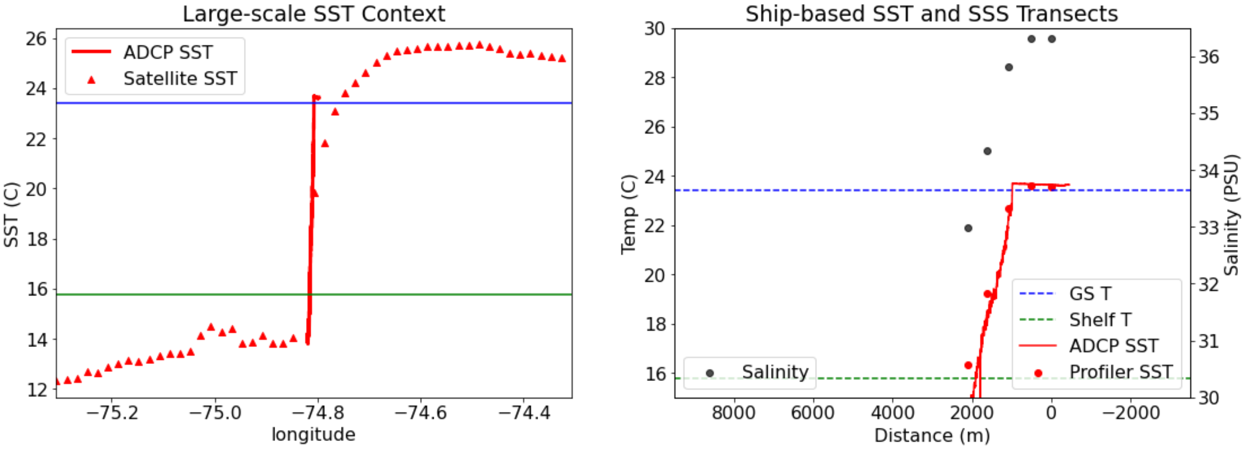

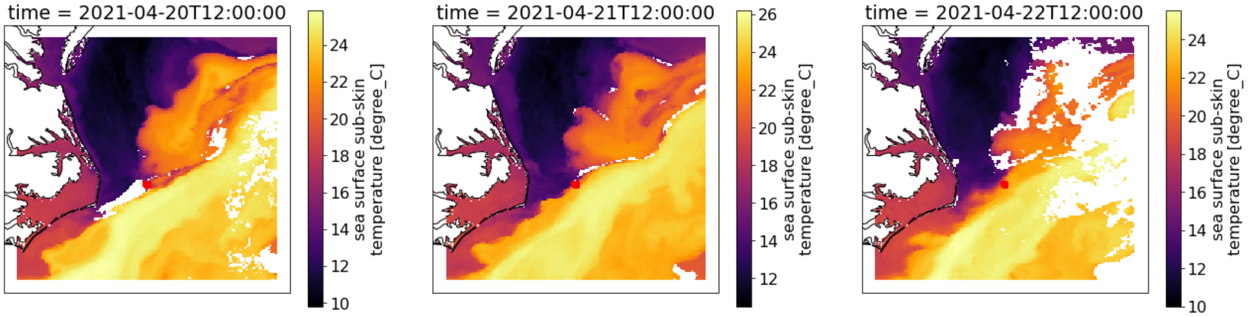


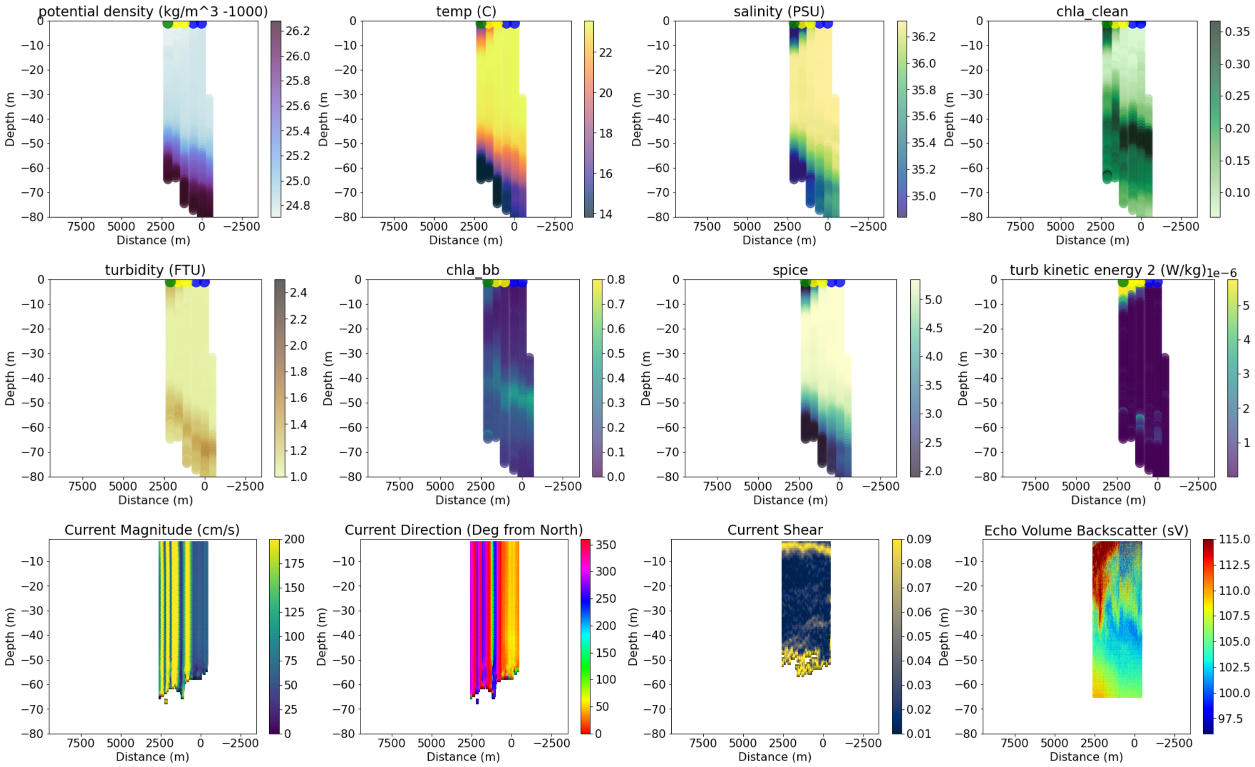


----------------------------------------
20
data/all_adcp/processed/101706_20210427T122655UTC_100s
Distance b/w GS and Coastal water is 62998.82820093336

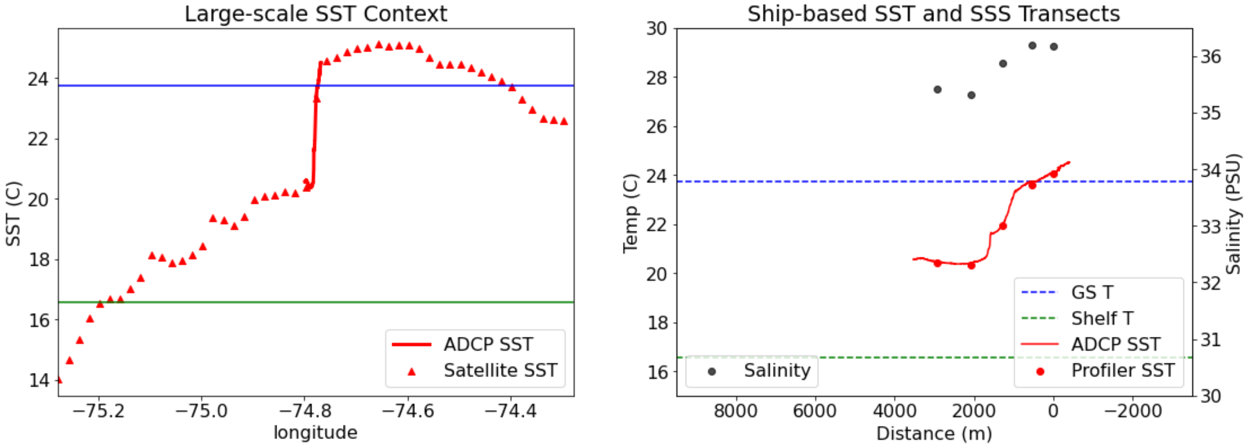

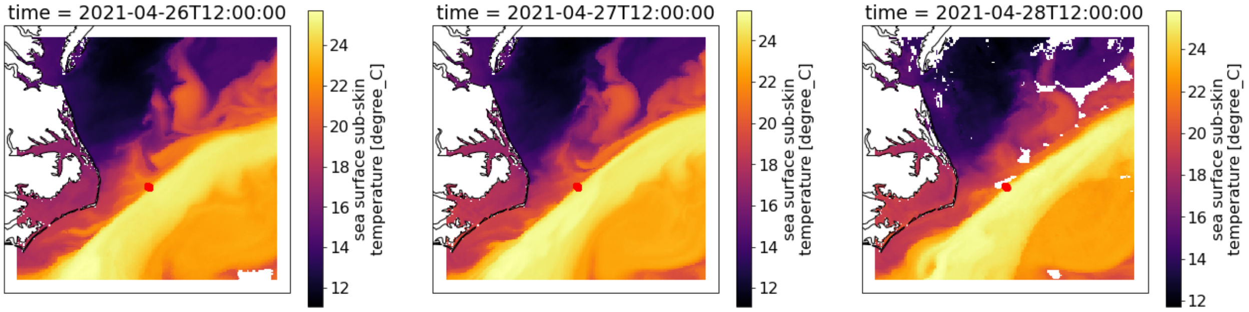

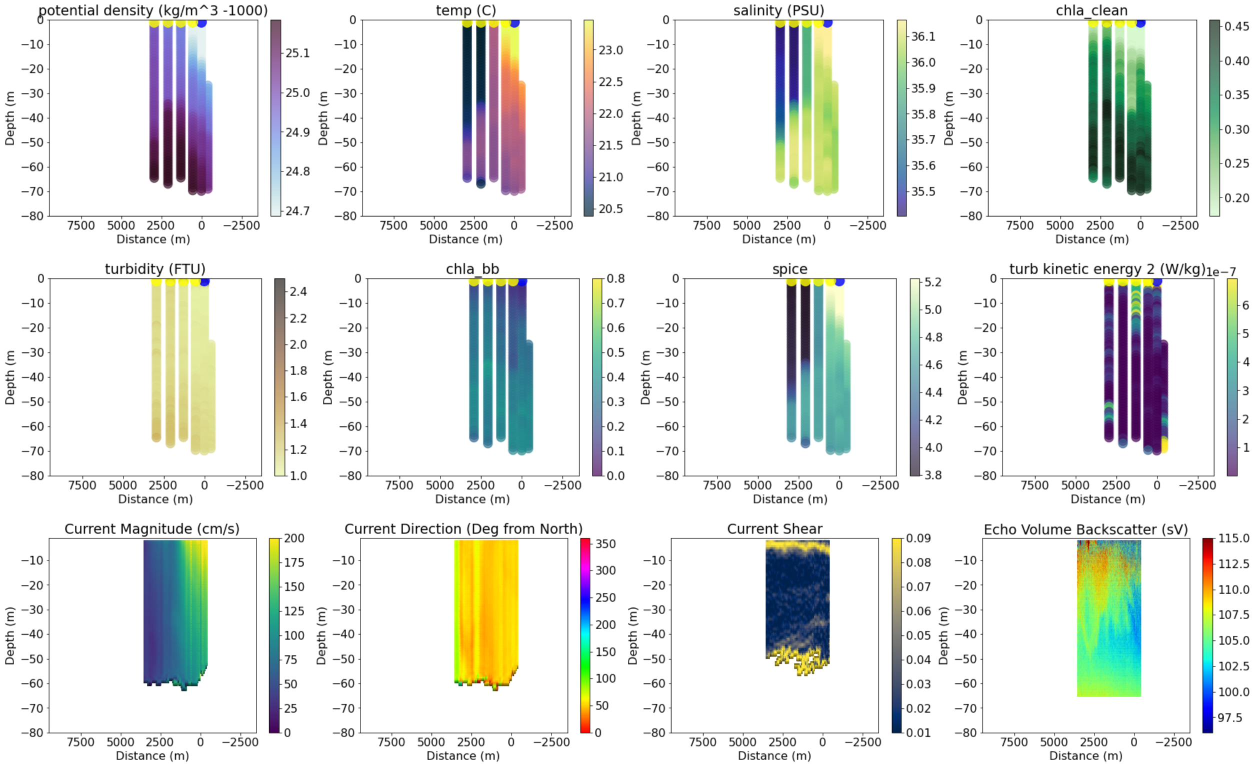

```
------------------------------------
22
data/all_adcp/processed/101706_20210615T121502UTC_100s
Transect does not cross into coastal water
```

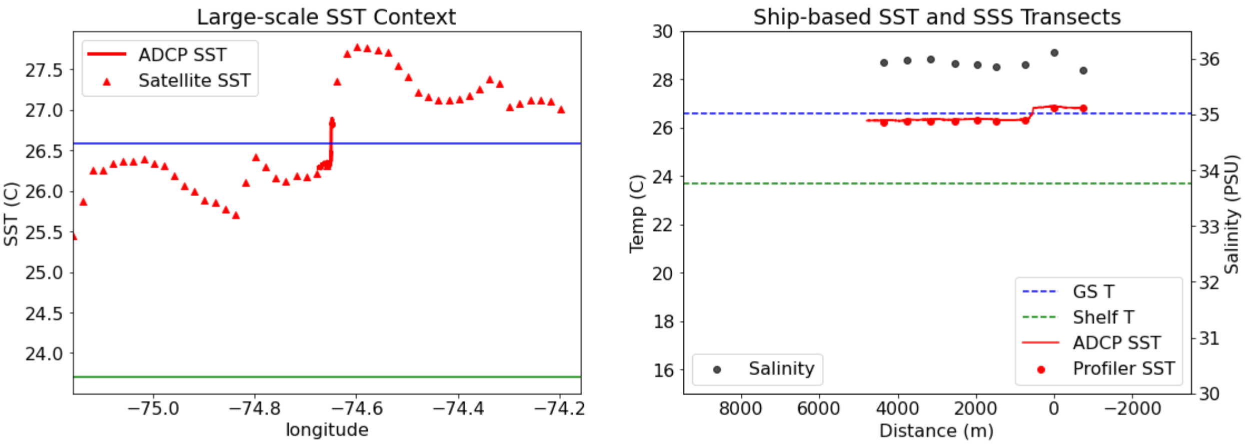

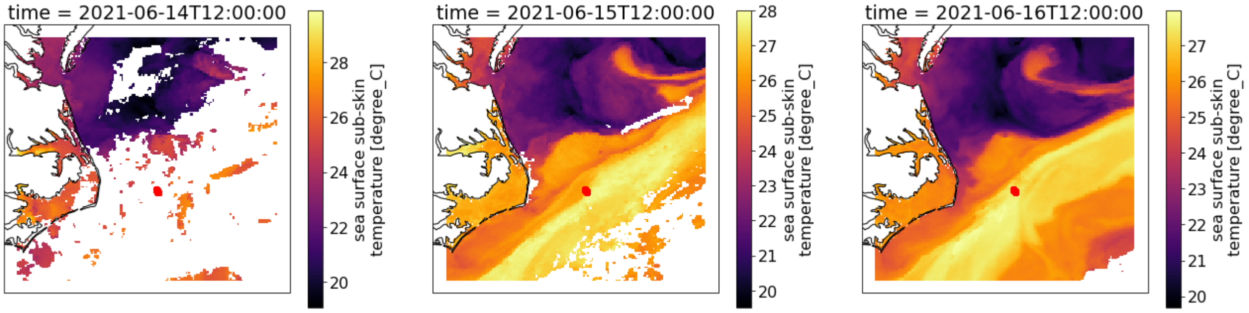


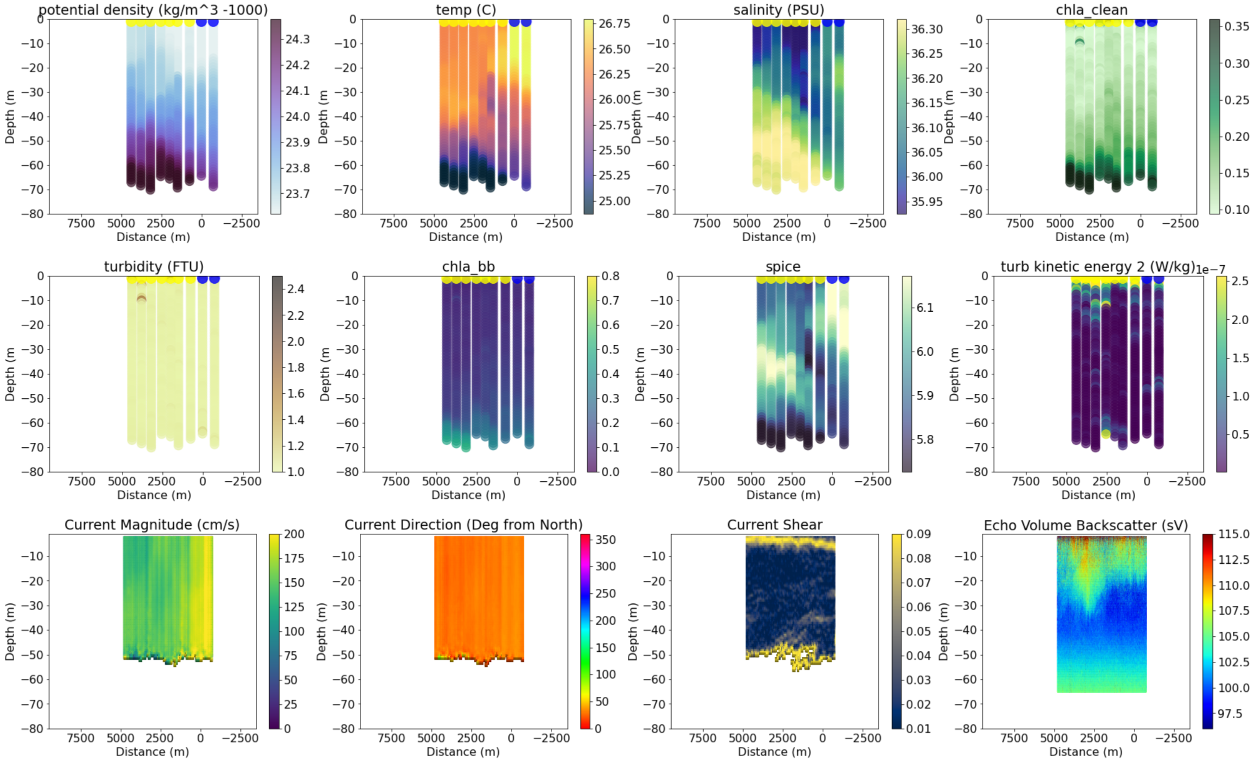


```
------------------------------------
23
data/all_adcp/processed/101706_20210615T144858UTC_100s
Transect does not cross into coastal water
```

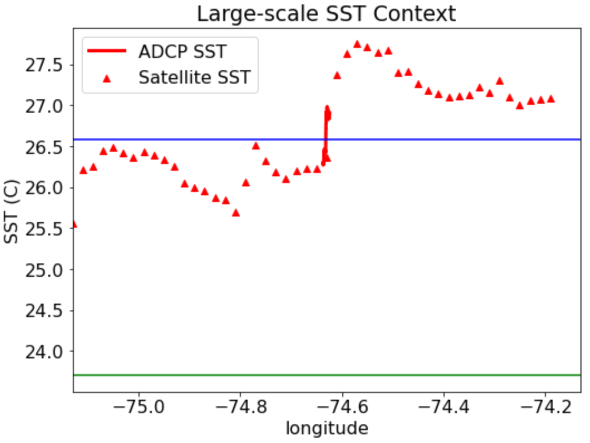


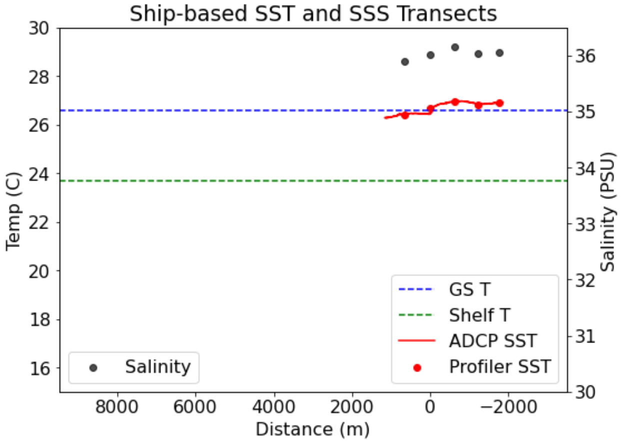

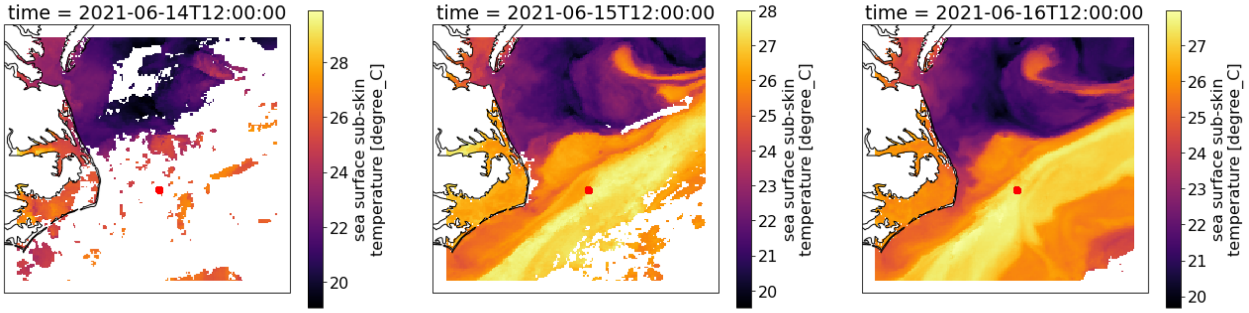

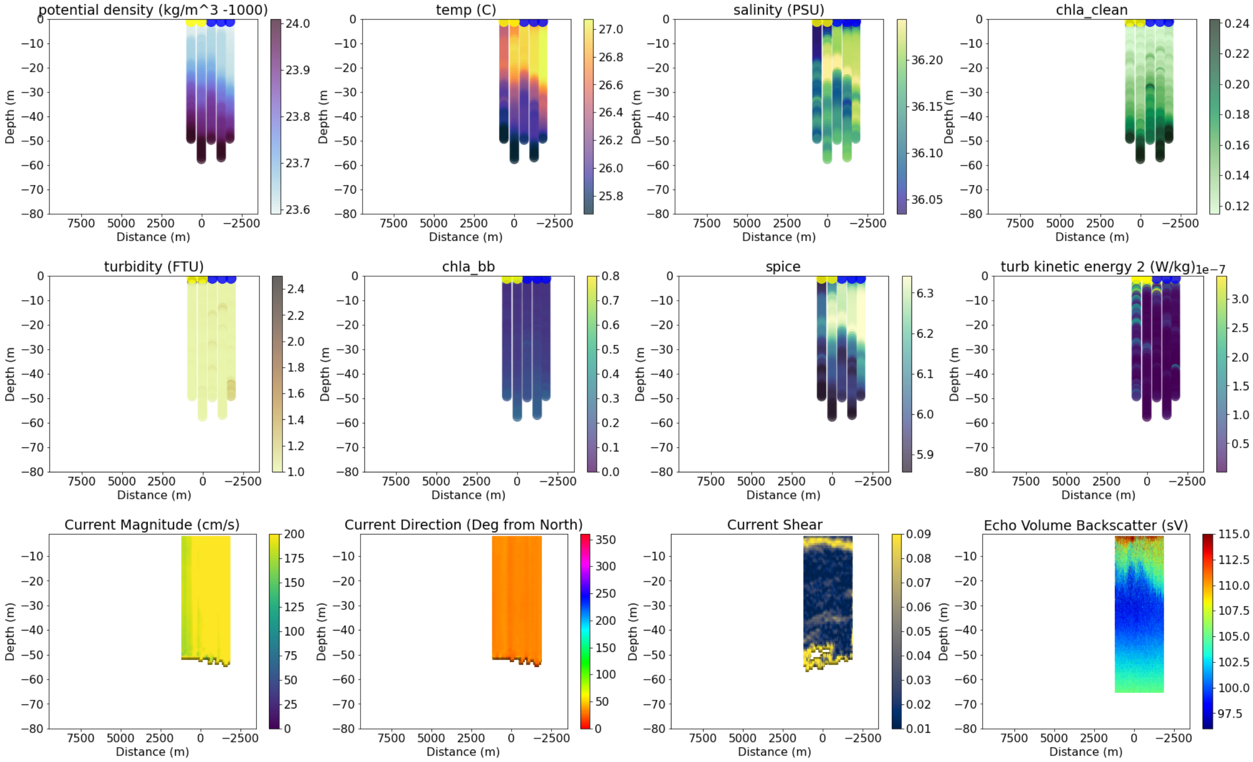

```
------------------------------------
24
data/all_adcp/processed/101706_20210616T124212UTC_badGPS_100s
Distance b/w GS and Coastal water is 62973.65484193107
```

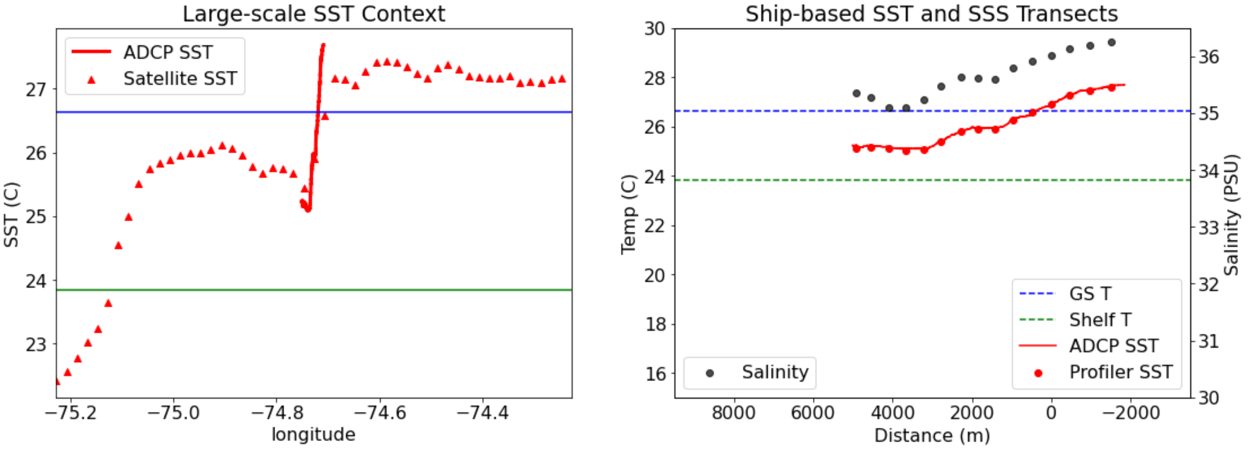

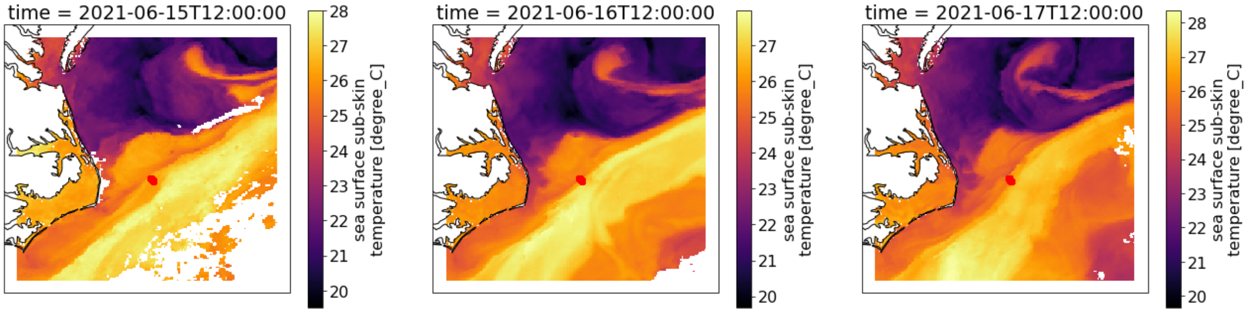


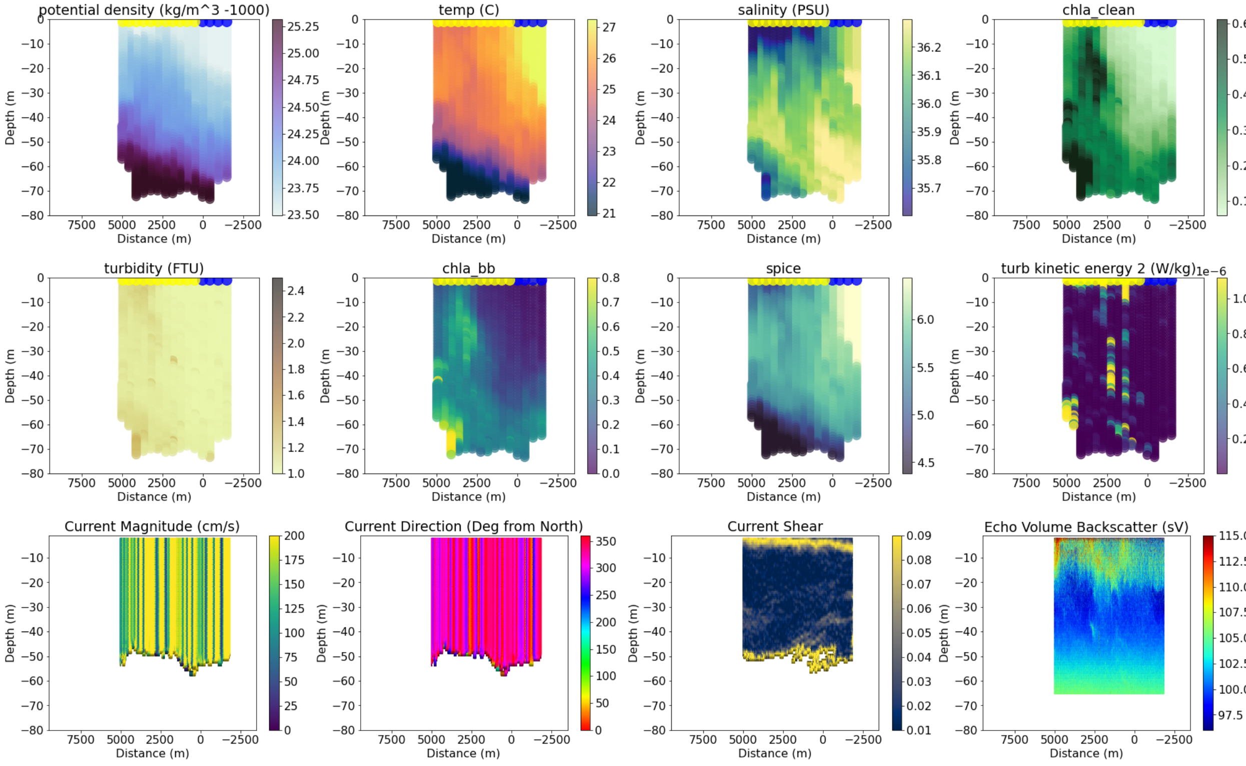


```
----------------------------------------
26
data/all_adcp/processed/101706_20210617T124353UTC_100s
Distance b/w GS and Coastal water is 71564.74461272624
```

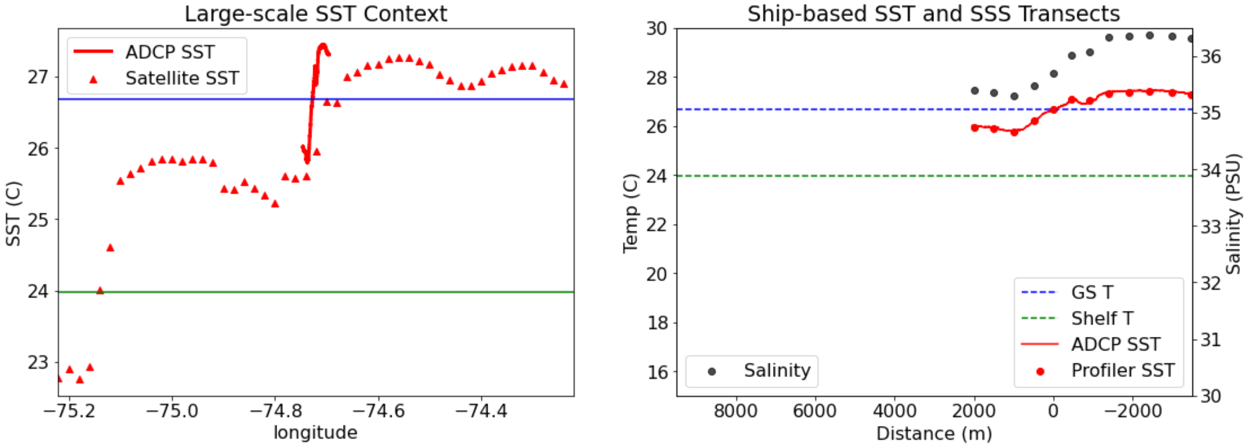

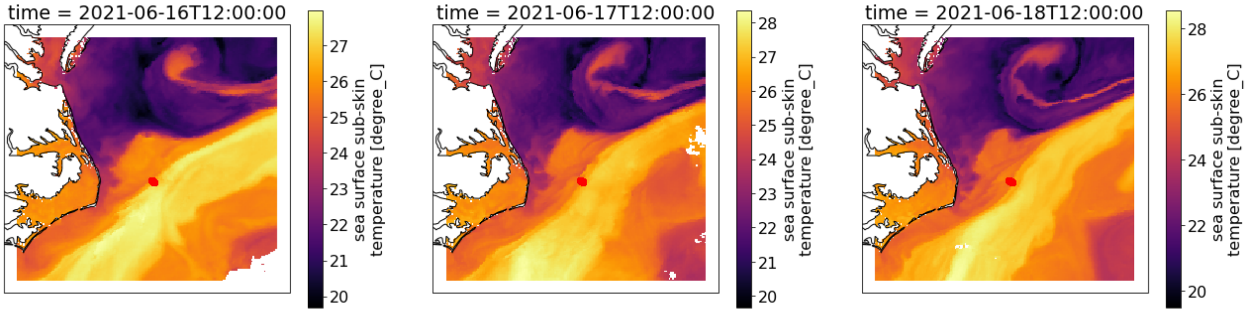


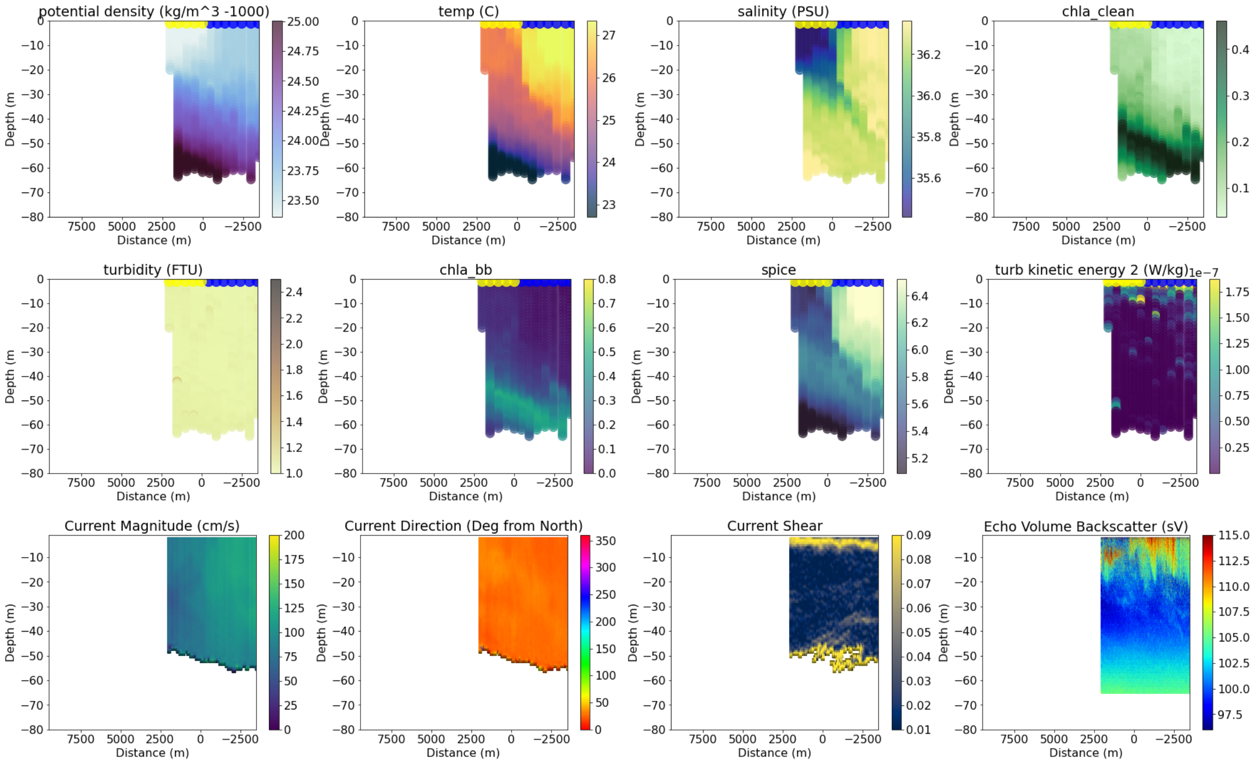


```
-------------------------------------
27
data/all_adcp/processed/101706_20210617T141632UTC_100s
Distance b/w GS and Coastal water is 62965.80565924342
```

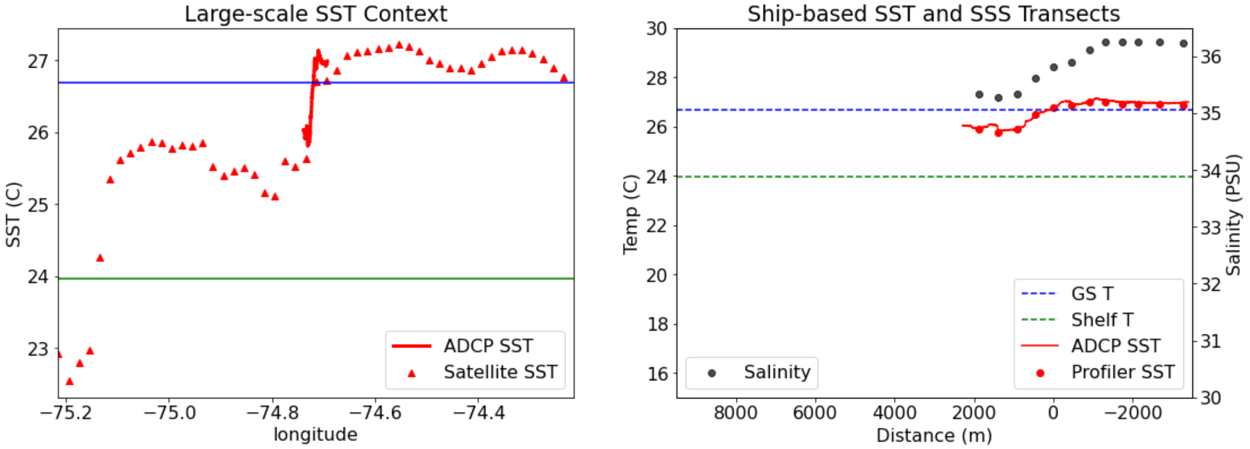

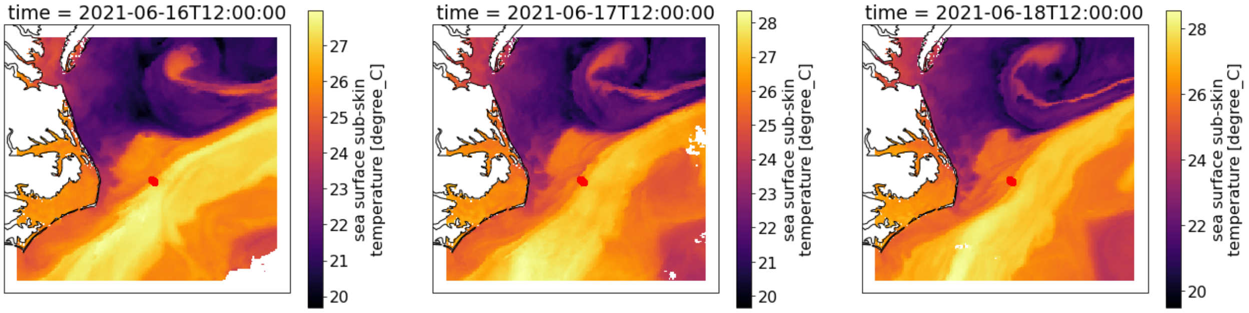


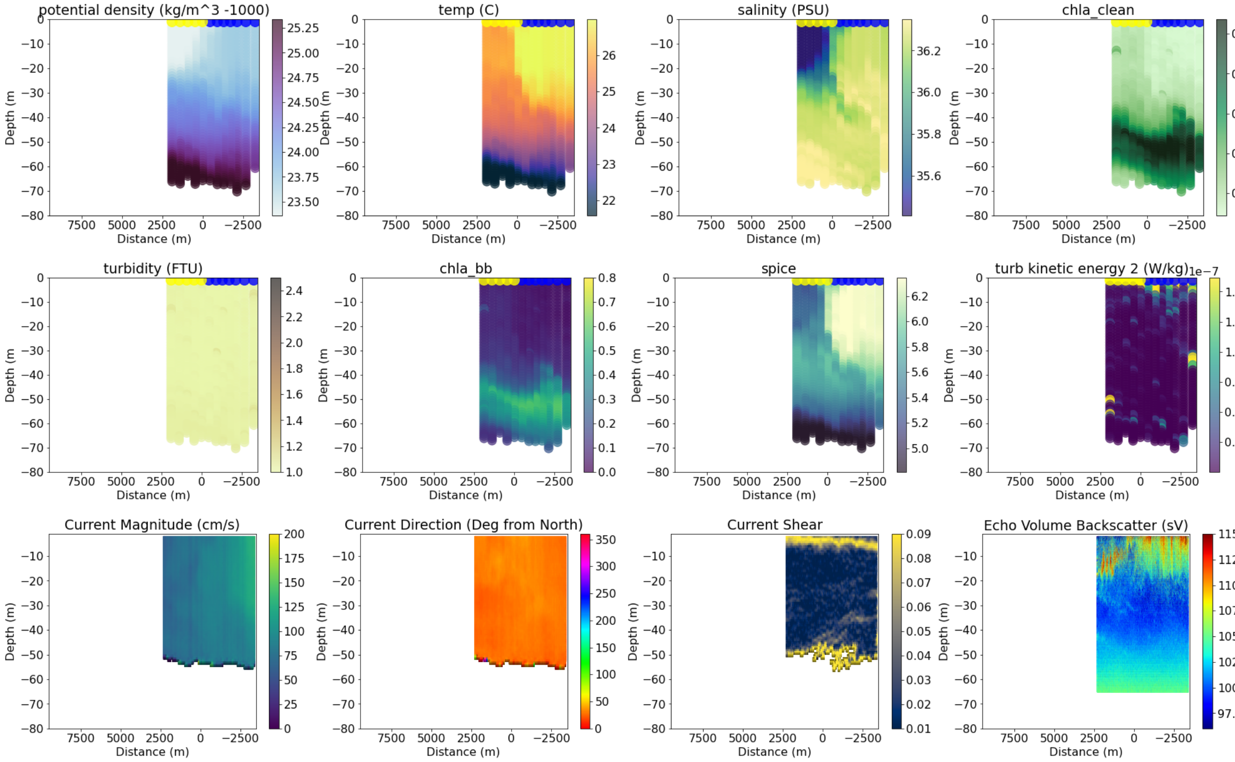


```
------------------------------------
29
data/all_adcp/processed/101706_20210826T120738UTC_100s
Distance b/w GS and Coastal water is 20047.909826163574
```

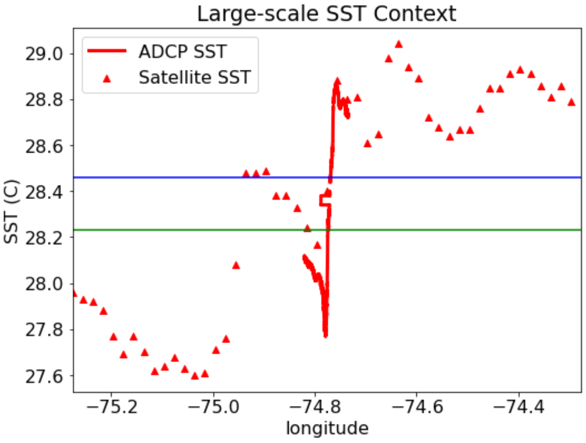


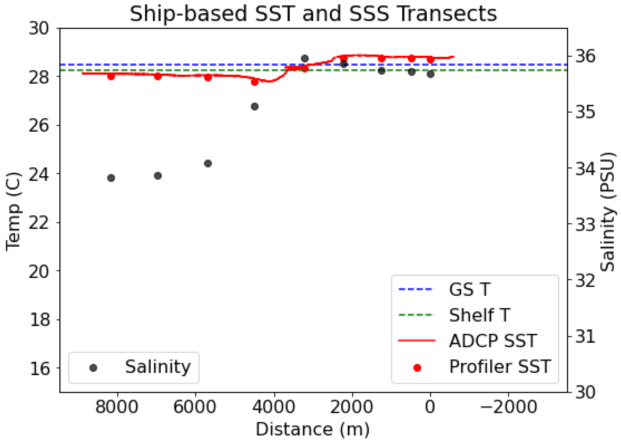

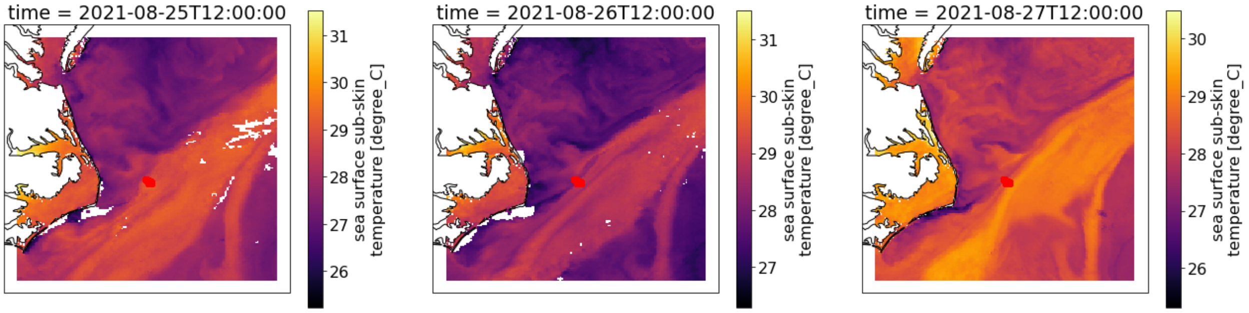


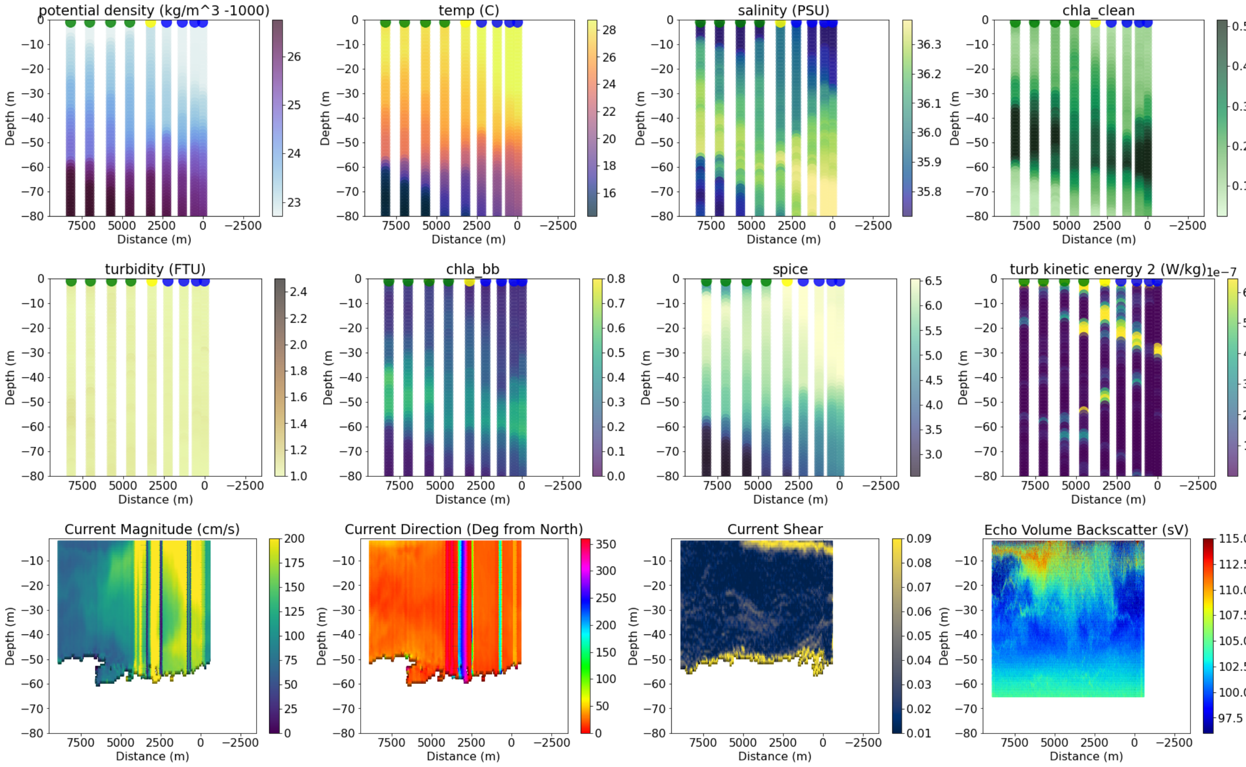


```
-----------------------------------
31
data/all_adcp/processed/101706_20210827T112801UTC_100s
Distance b/w GS and Coastal water is 5729.217364111245
```

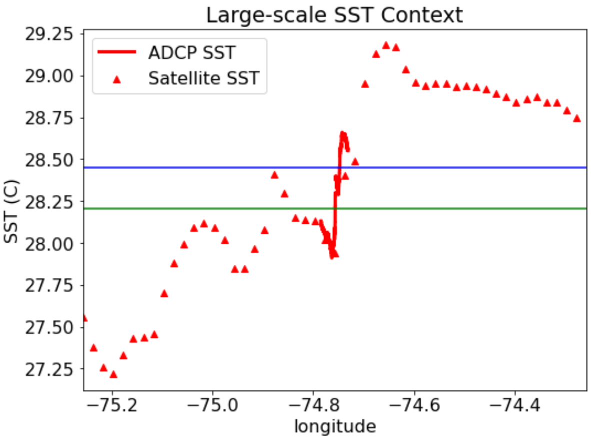


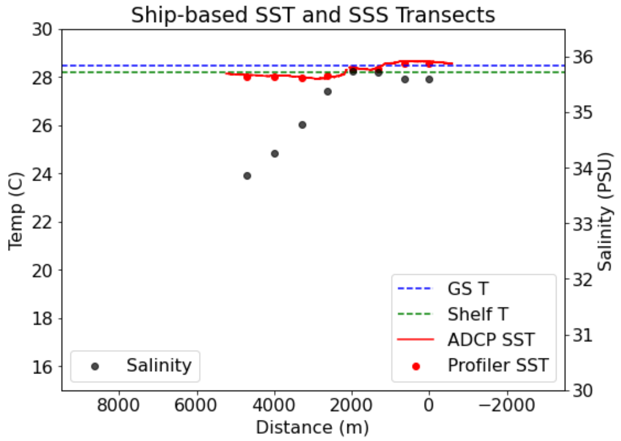

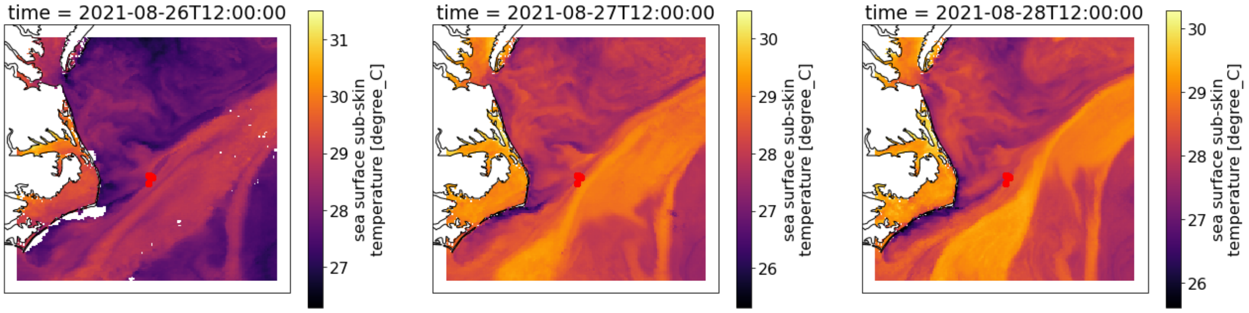


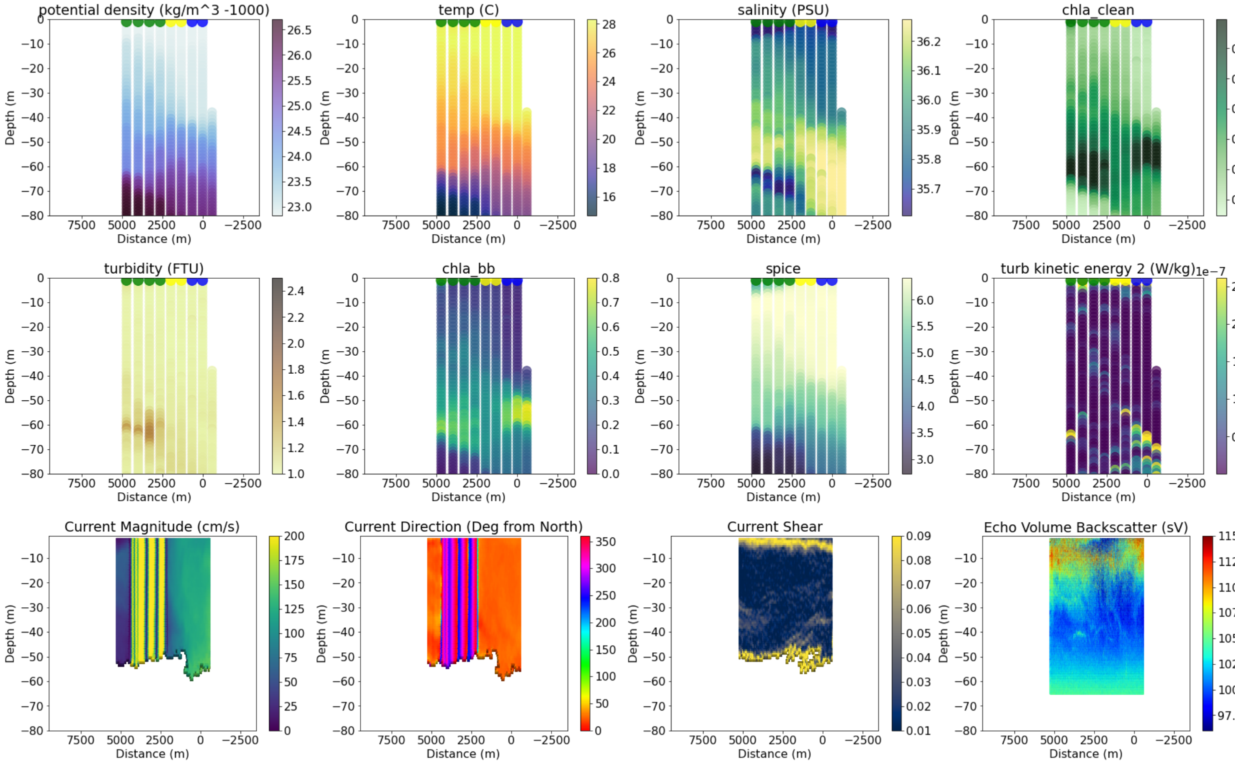


```
------------------------------------
34
data/all_adcp/processed/101706_20210828T104556UTC_100s
Distance b/w GS and Coastal water is 65975.7612004256
```

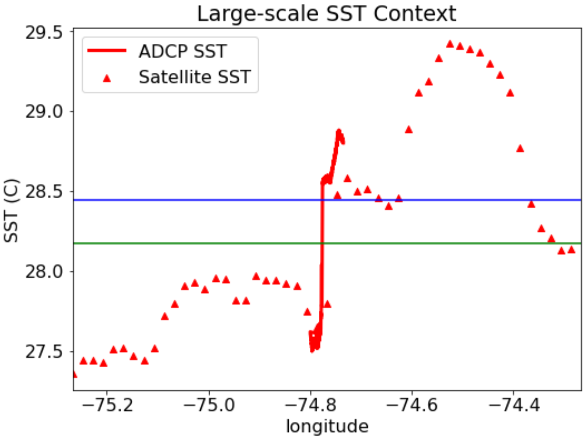


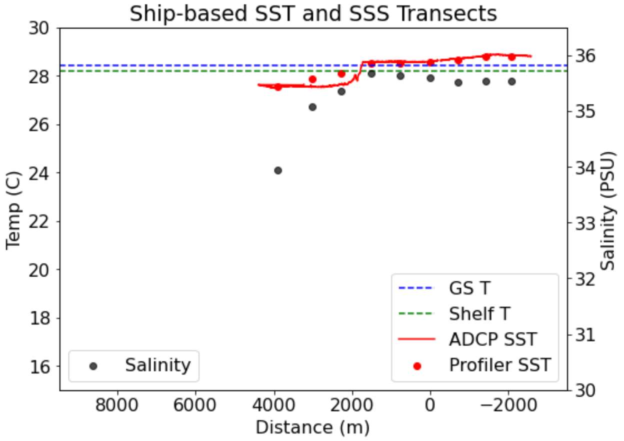

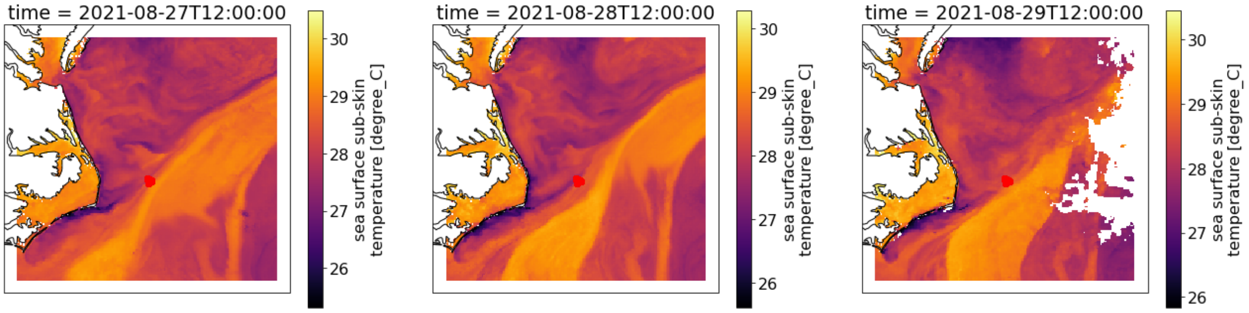


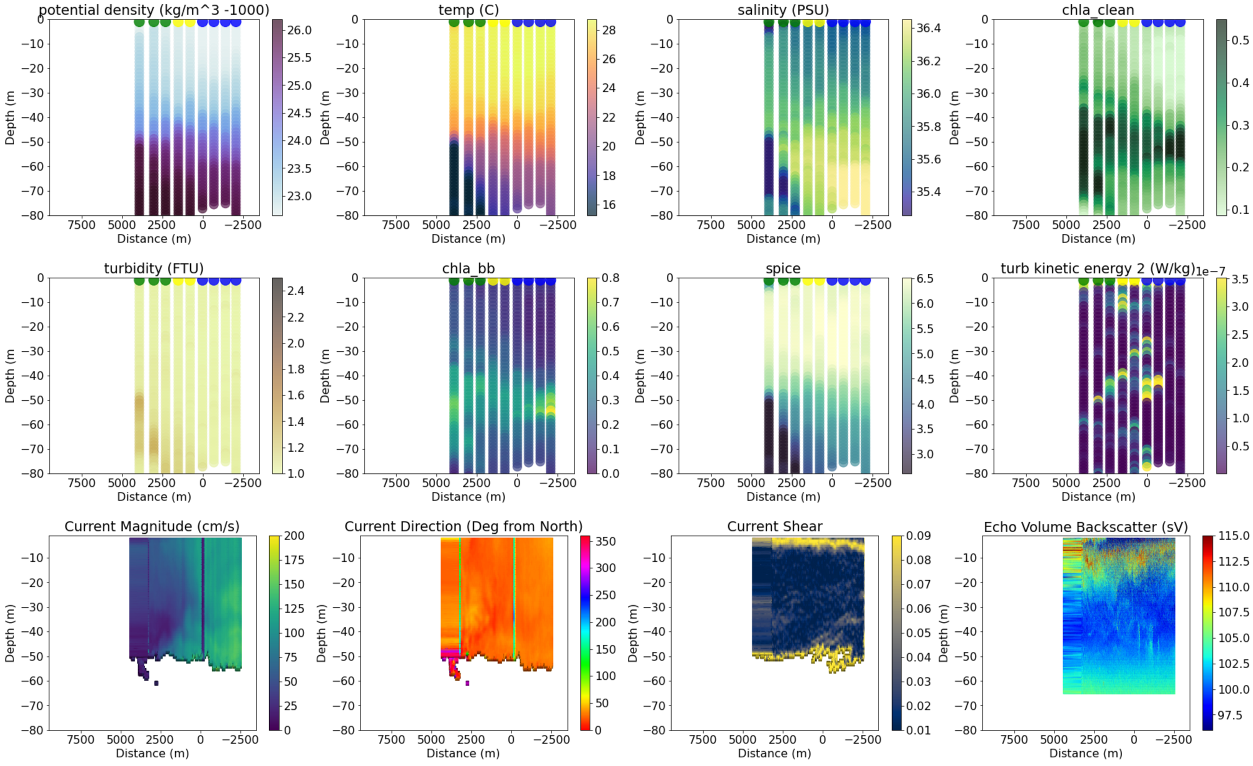


```
--------------------------------------
35
data/all_adcp/processed/101706_20210828T151028UTC_100s
Distance b/w GS and Coastal water is 65945.68891895749
```

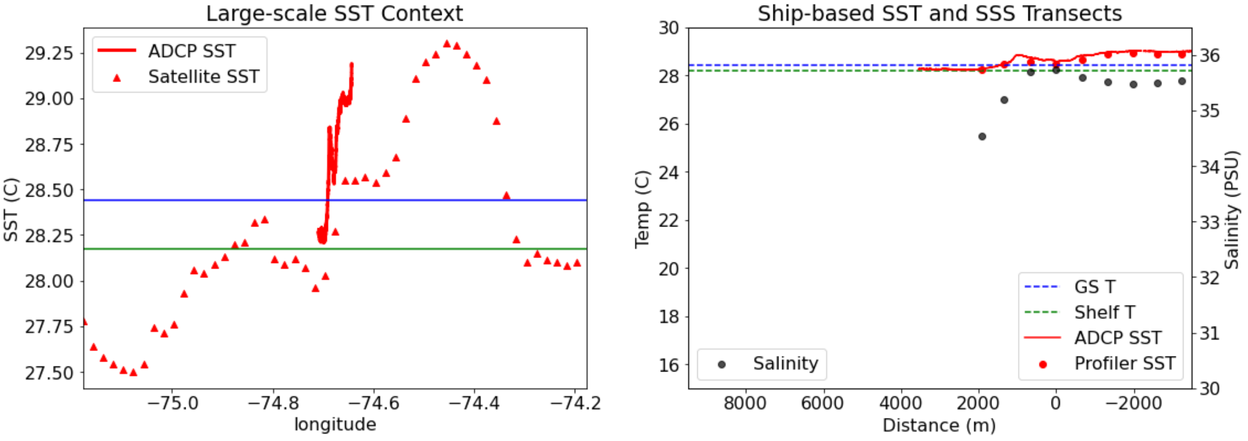

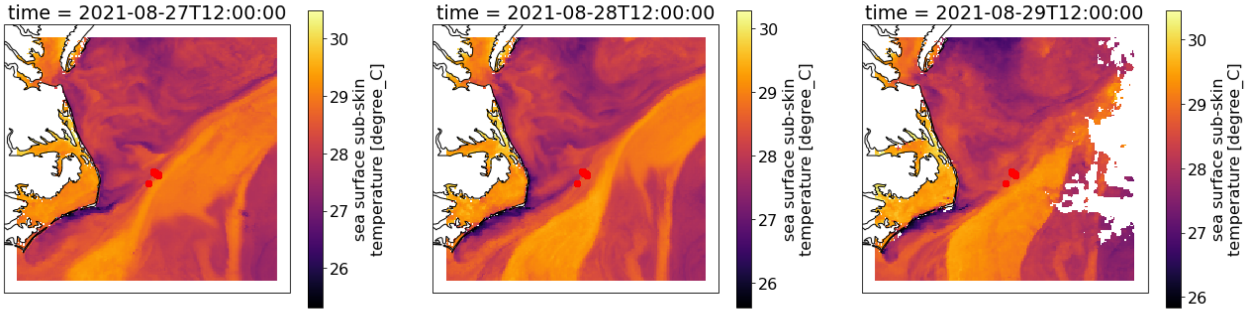

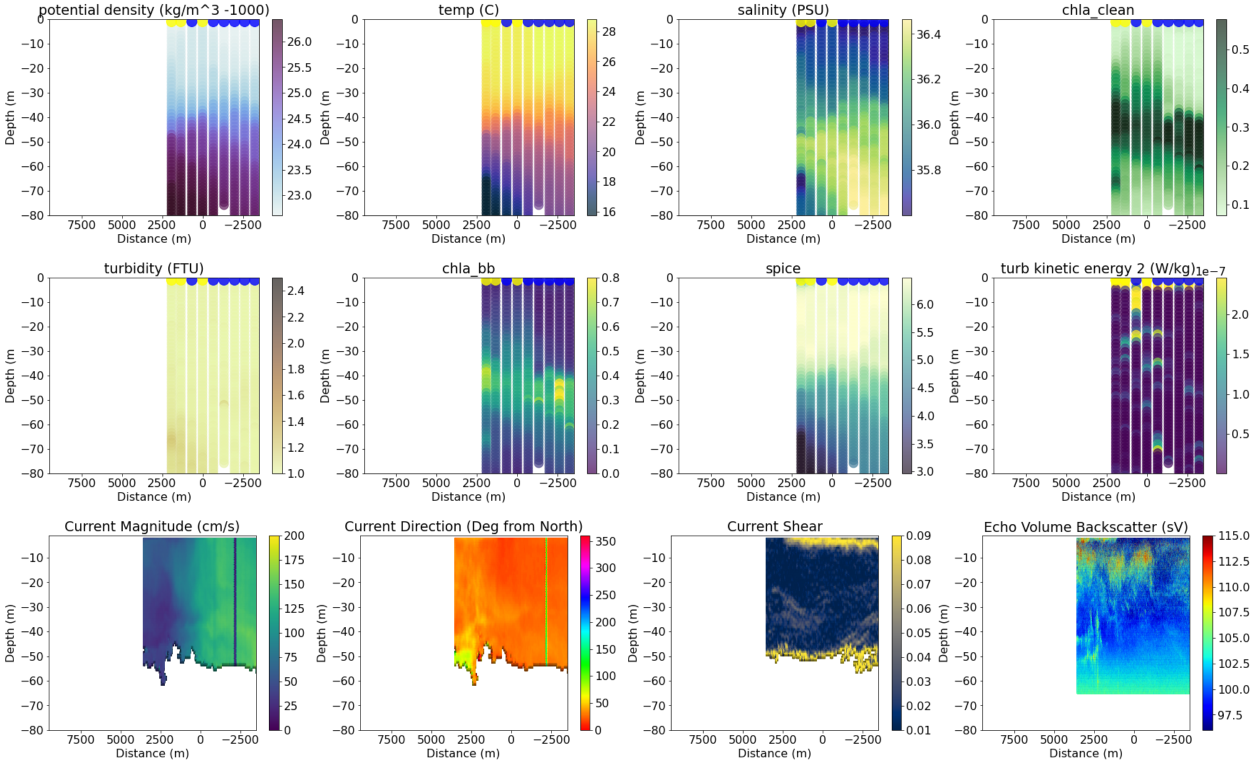

```
------------------------------------
38
data/all_adcp/processed/101706_20210829T162104UTC_100s
Distance b/w GS and Coastal water is 8579.519847171594
```

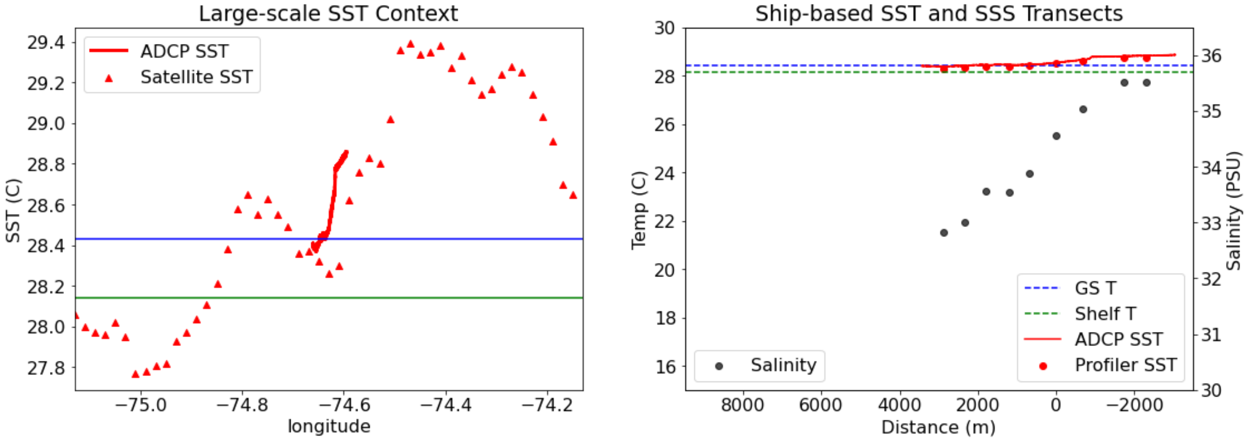

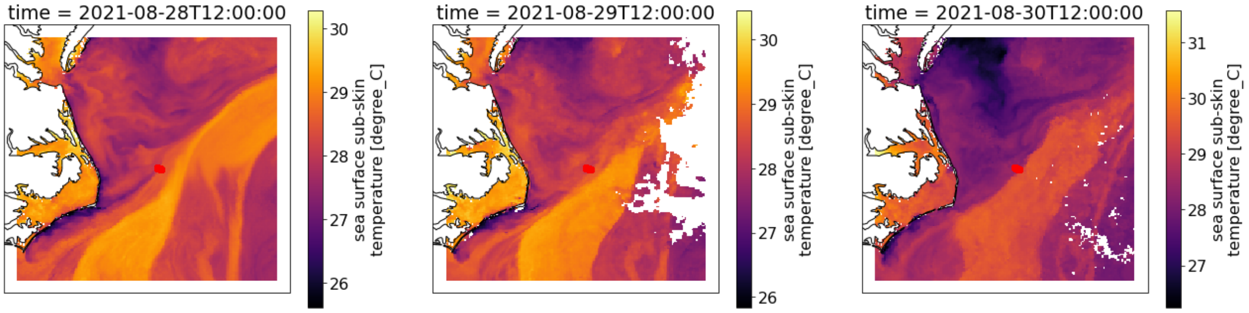


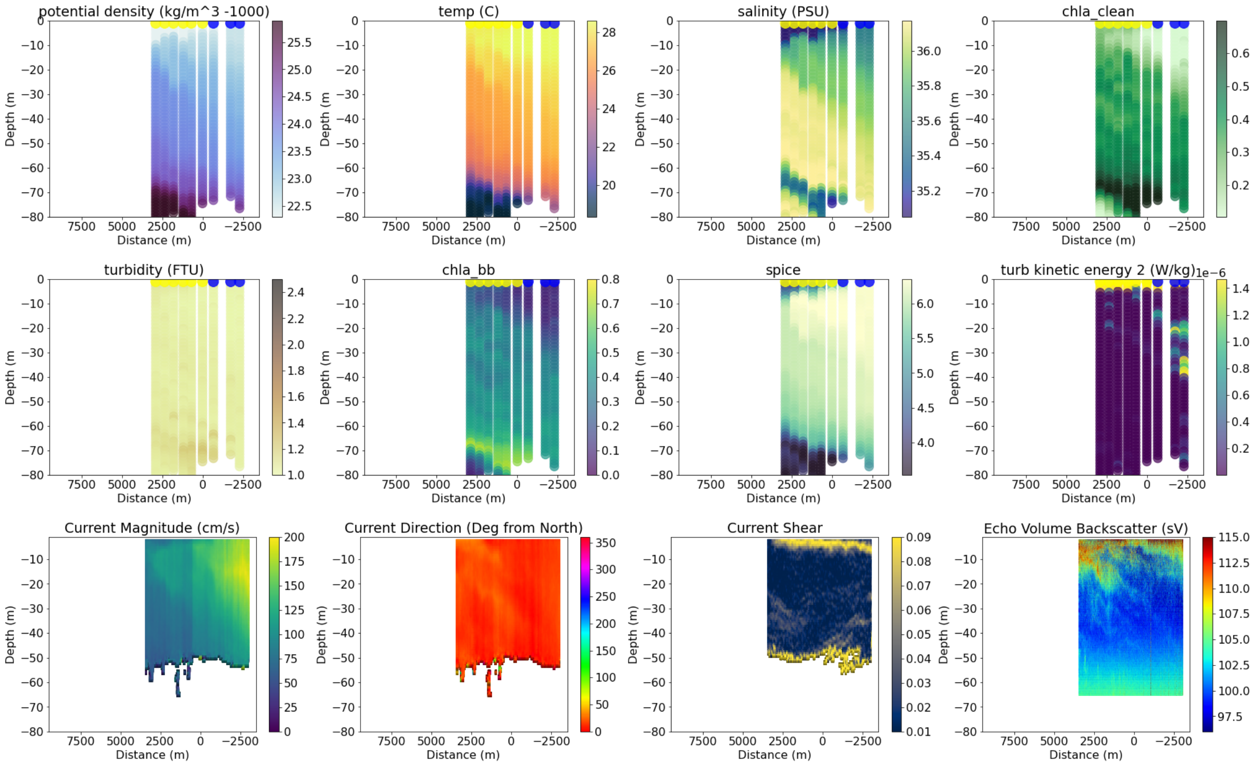


```
-----------------------------------
40
data/all_adcp/processed/101706_20210906T121429UTC_100s
Distance b/w GS and Coastal water is 2861.3597246488307
```

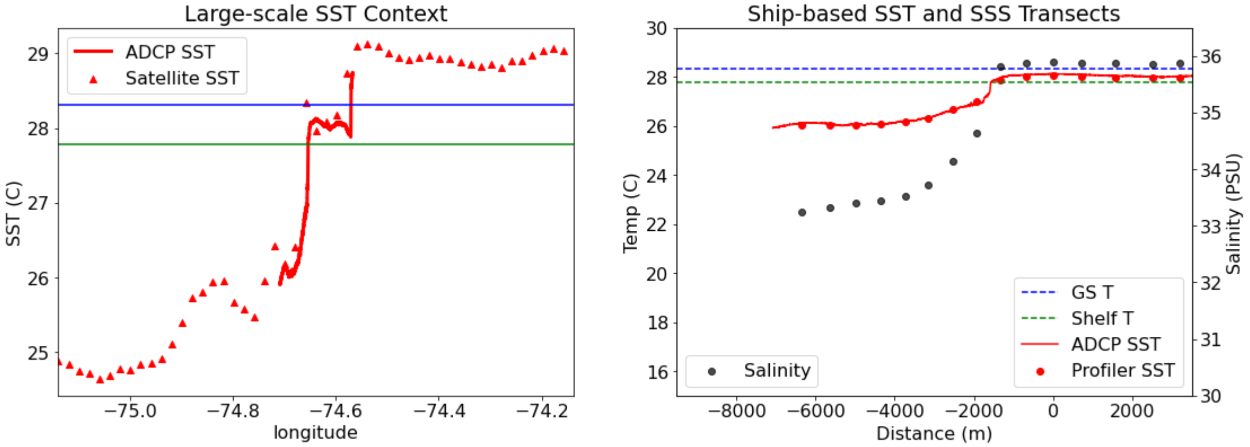

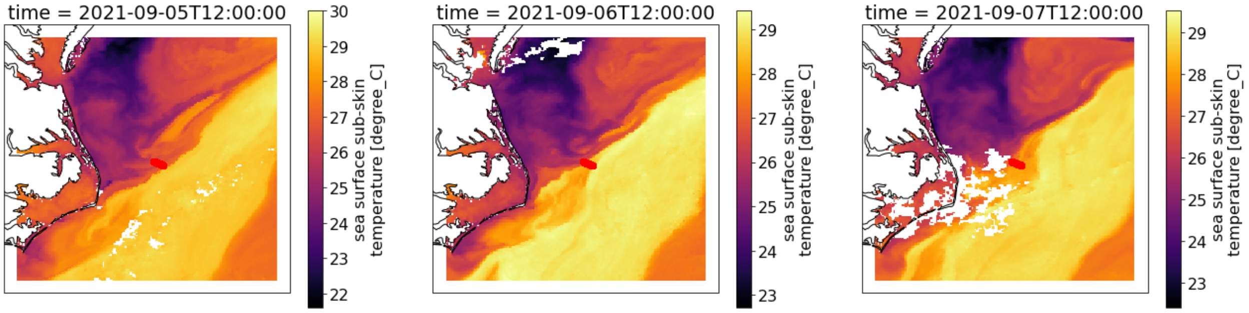


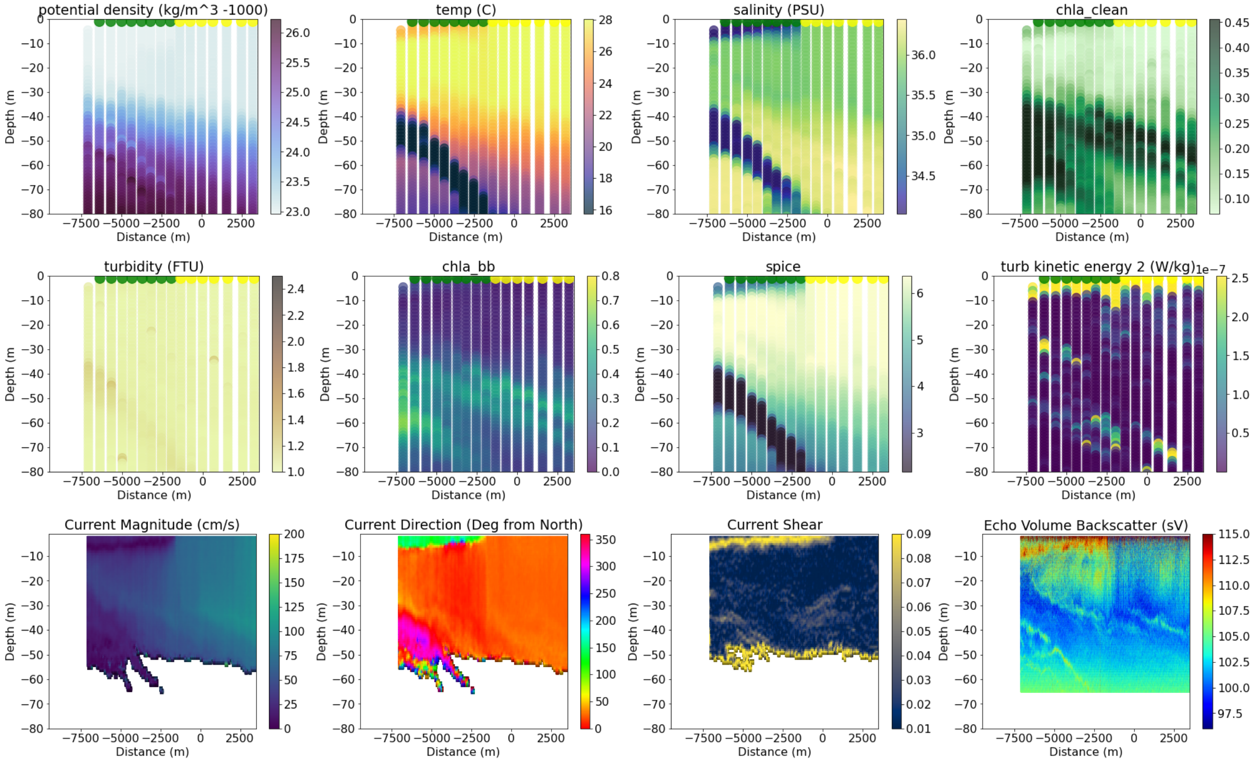


```
-------------------------------------
41
data/all_adcp/processed/101706_20210906T164454UTC_100s
Distance b/w GS and Coastal water is 2861.7423197417843
```

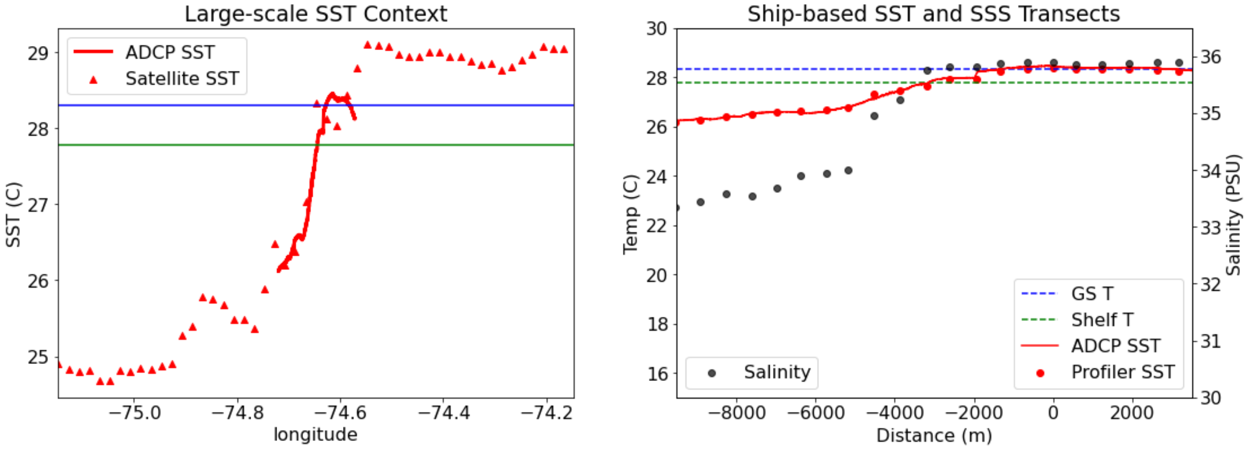

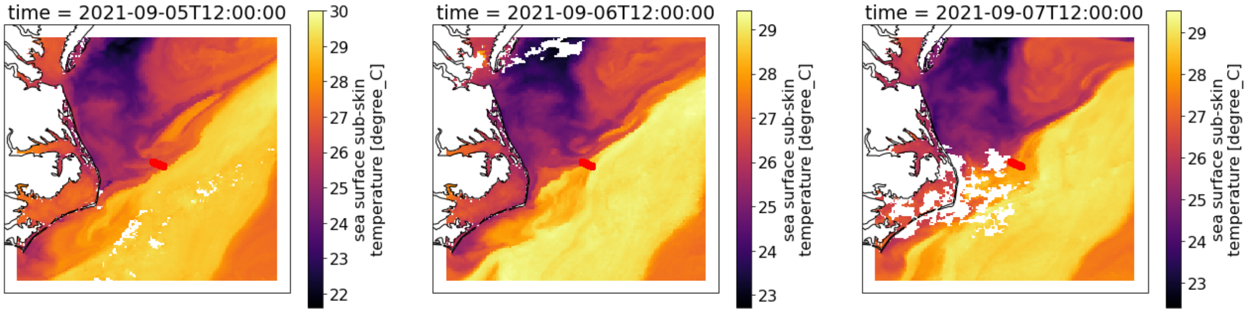


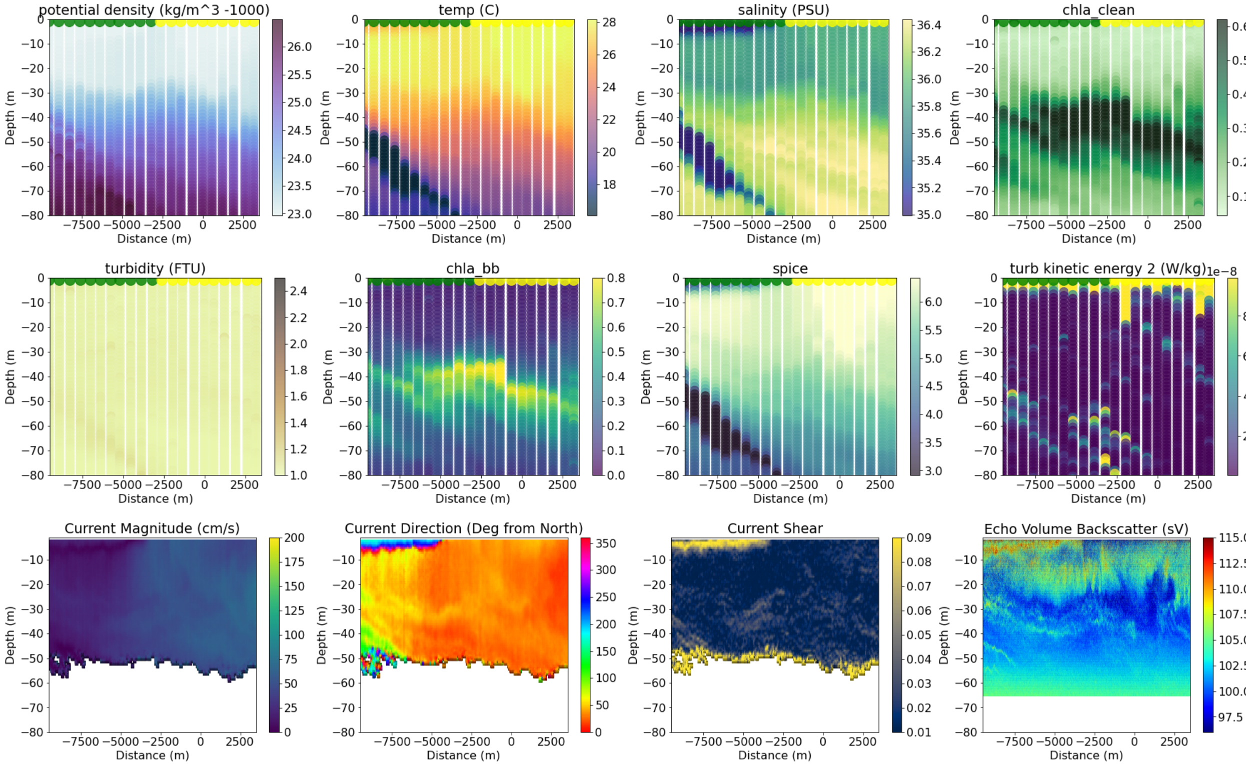


```
----------------------------------------
42
data/all_adcp/processed/101706_20210907T123502UTC_100s
Distance b/w GS and Coastal water is 37243.7983945212
```

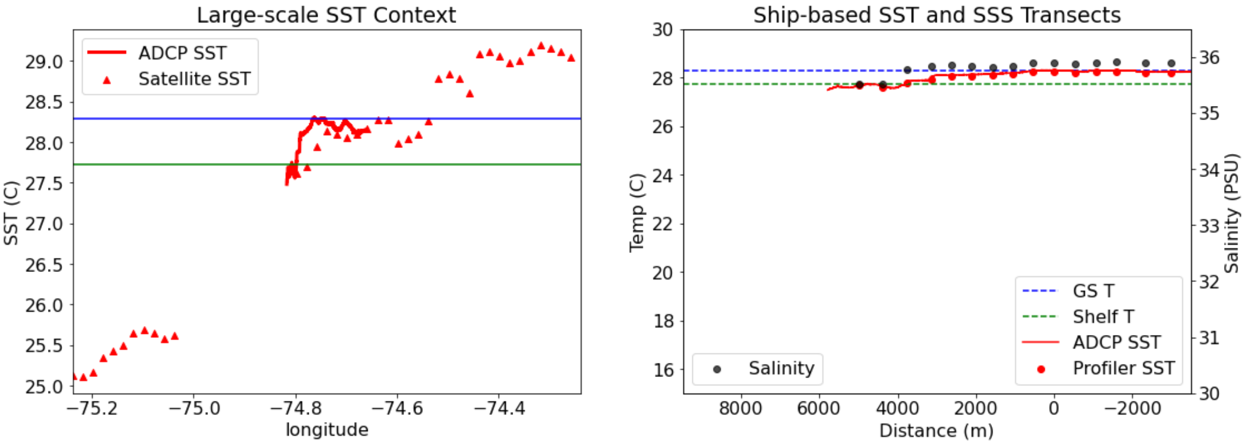

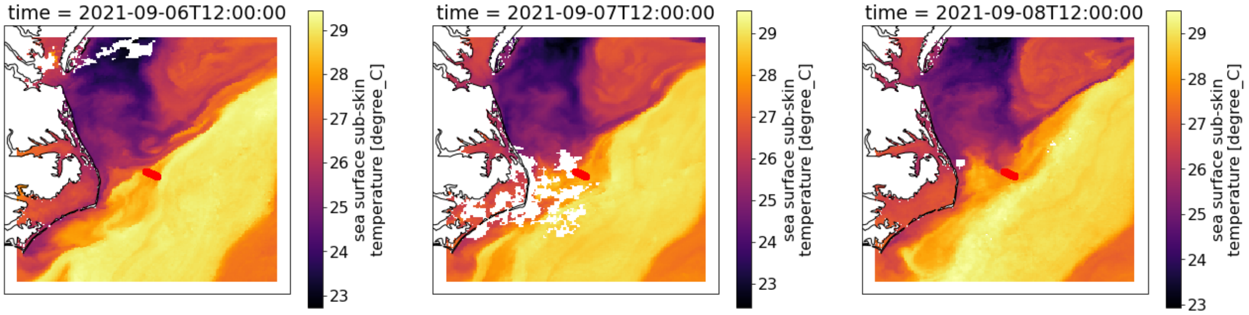


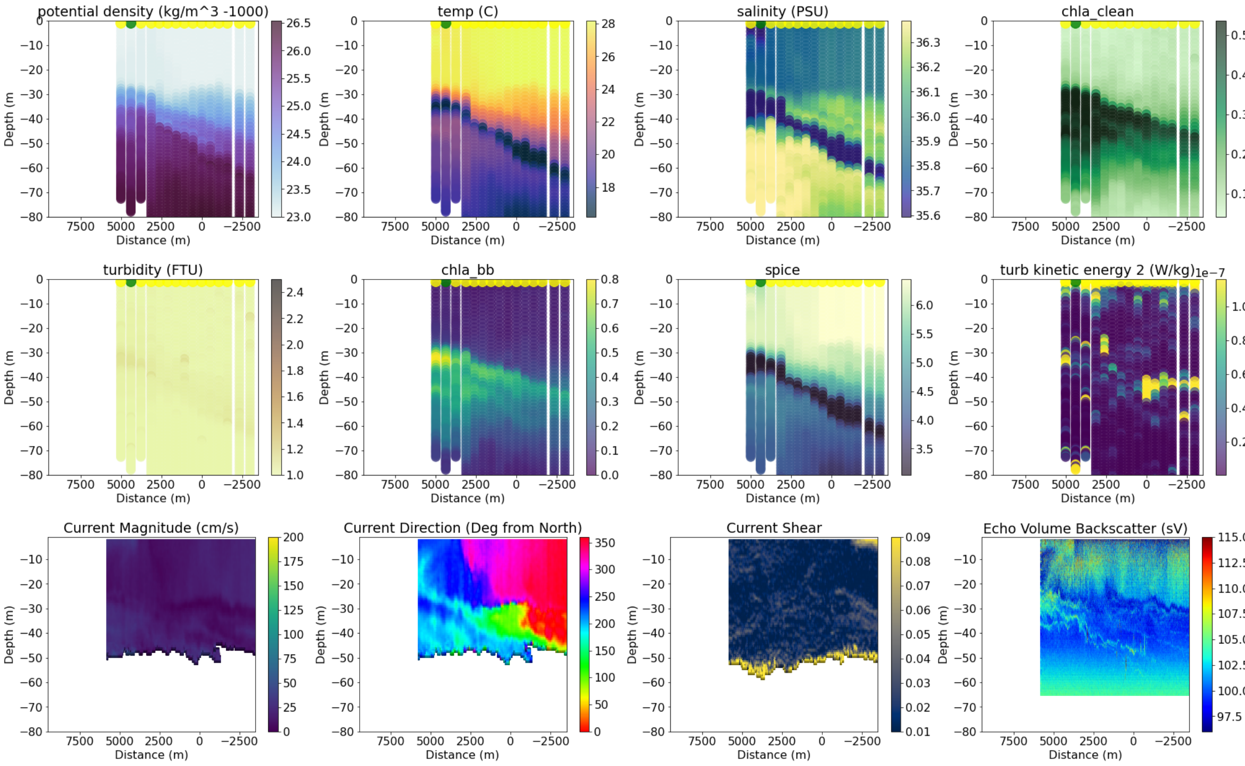


```
-----------------------------------
43
data/all_adcp/processed/101706_20211005T124141UTC_100s
Distance b/w GS and Coastal water is 25773.966086211552
```

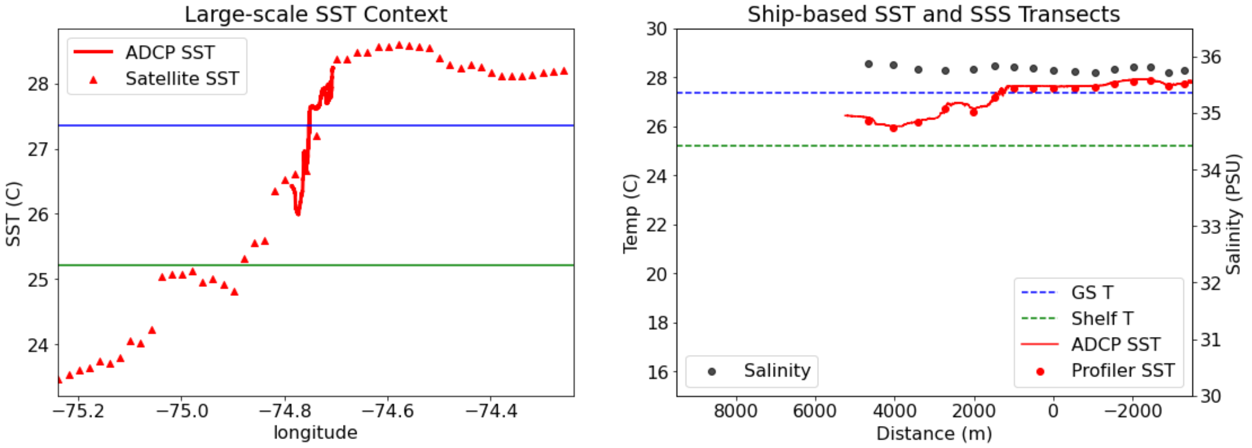

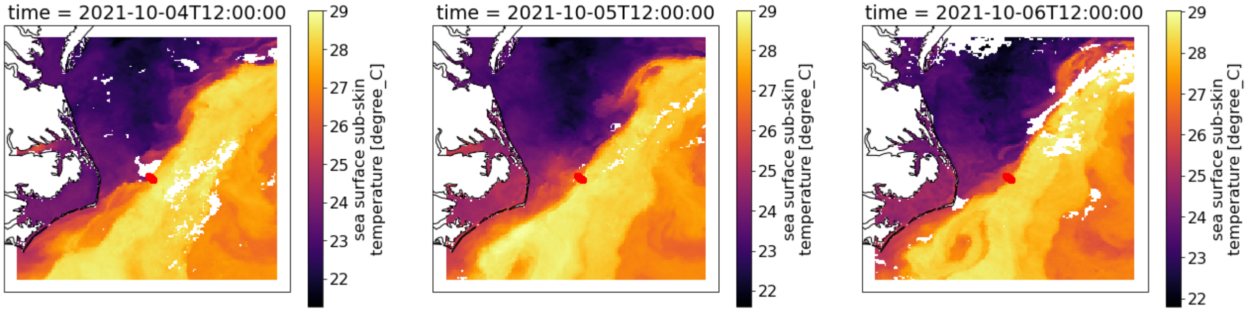


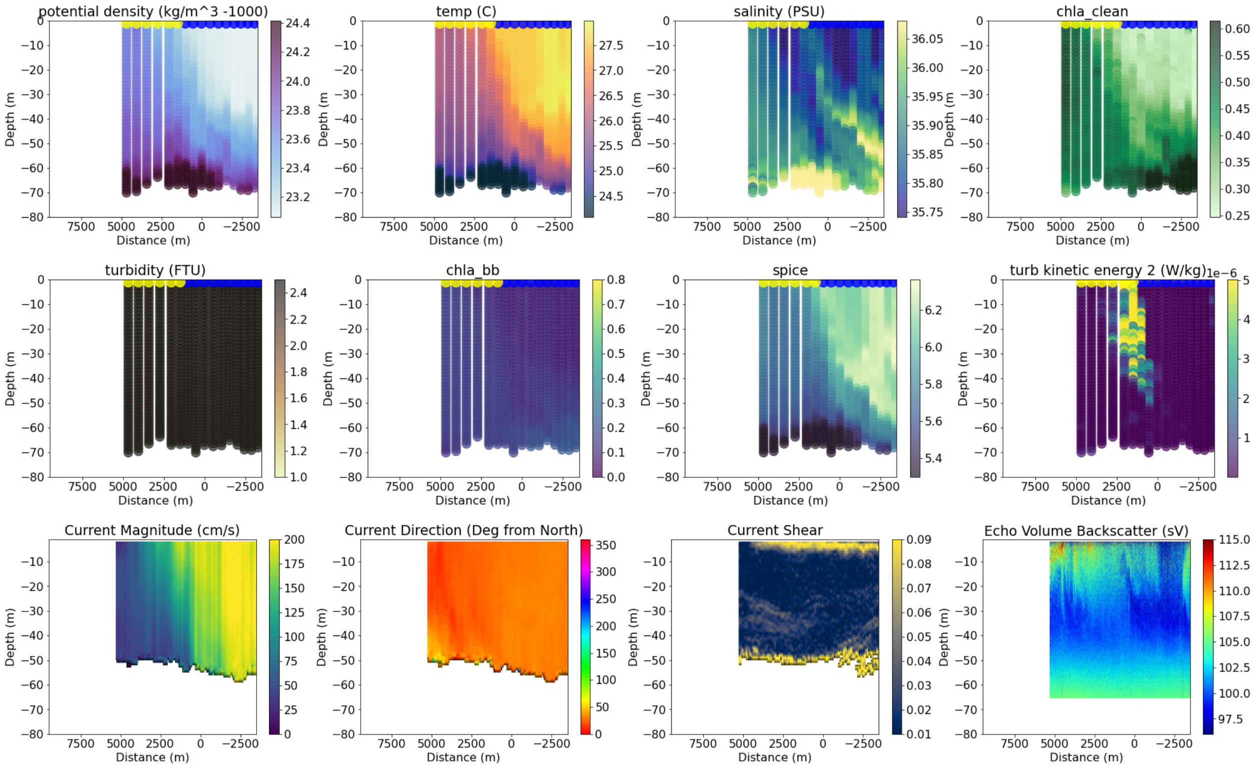


```
-------------------------------------
44
data/all_adcp/processed/101706_20211005T141731UTC_100s
Distance b/w GS and Coastal water is 25772.31156997663
```

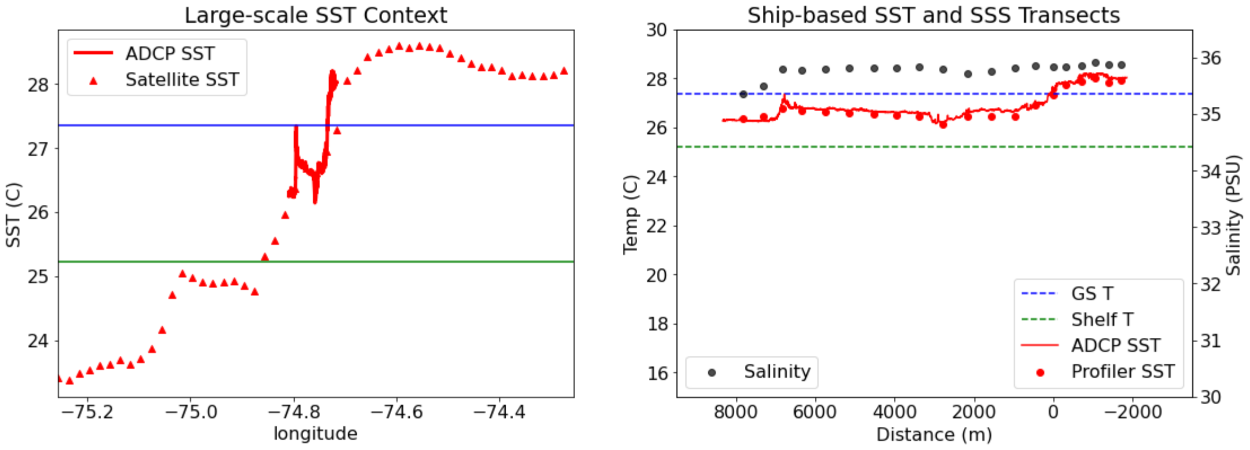

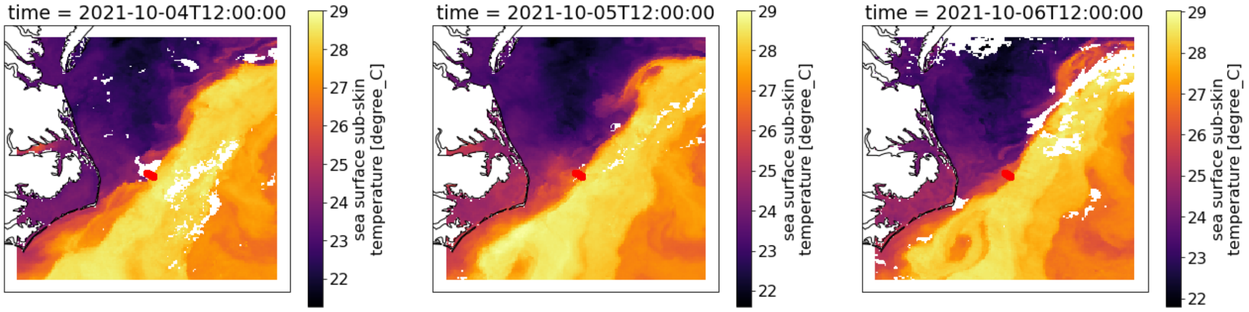


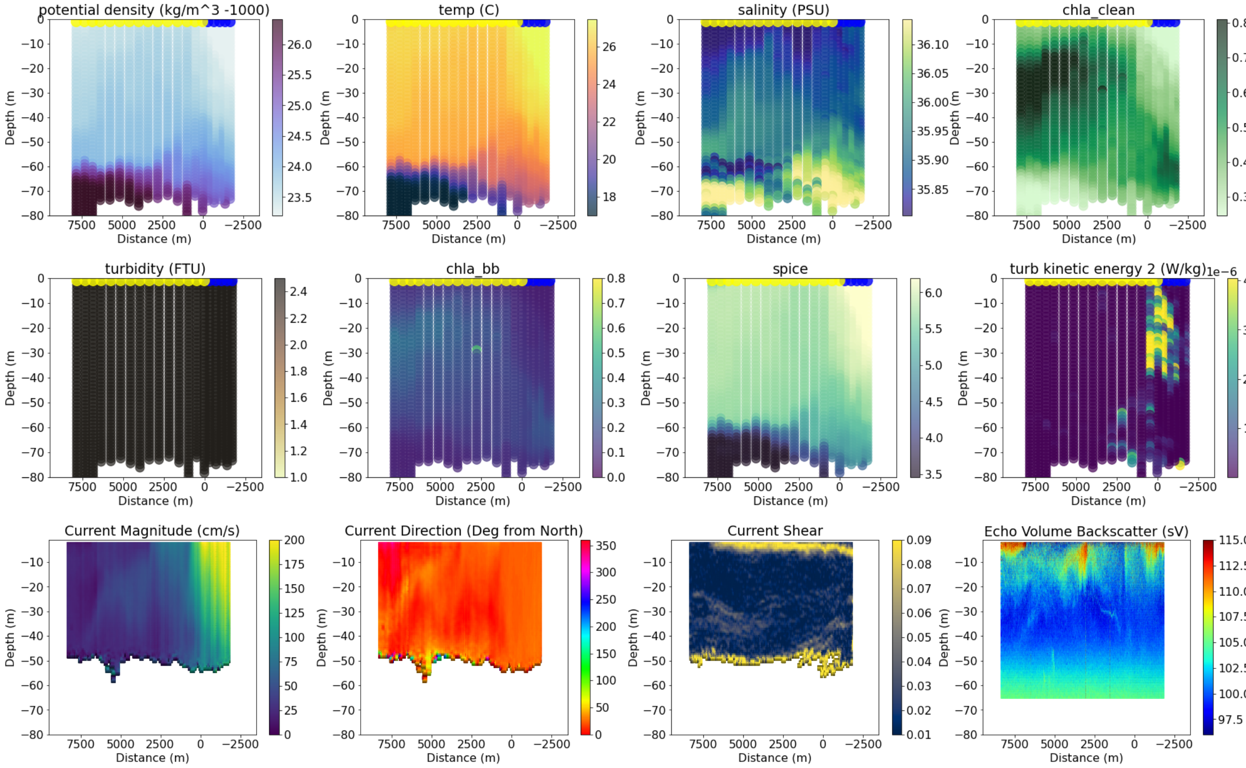


```
----------------------------------------
45
data/all_adcp/processed/101706_20211006T130945UTC_100s
Distance b/w GS and Coastal water is 22912.19596784995
```

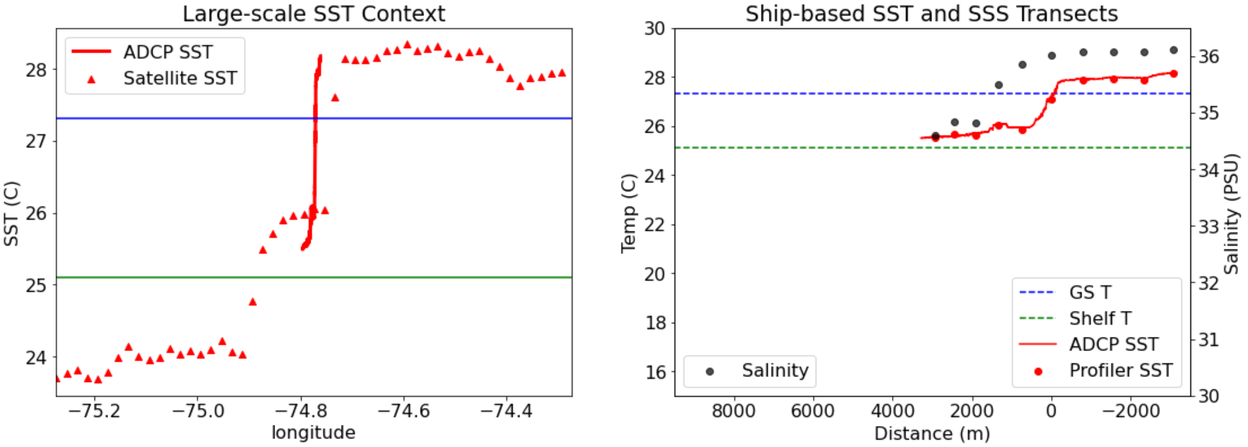

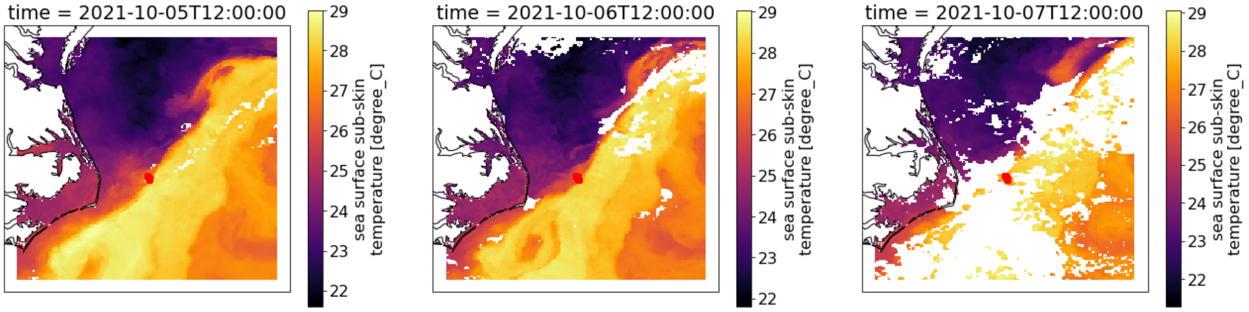

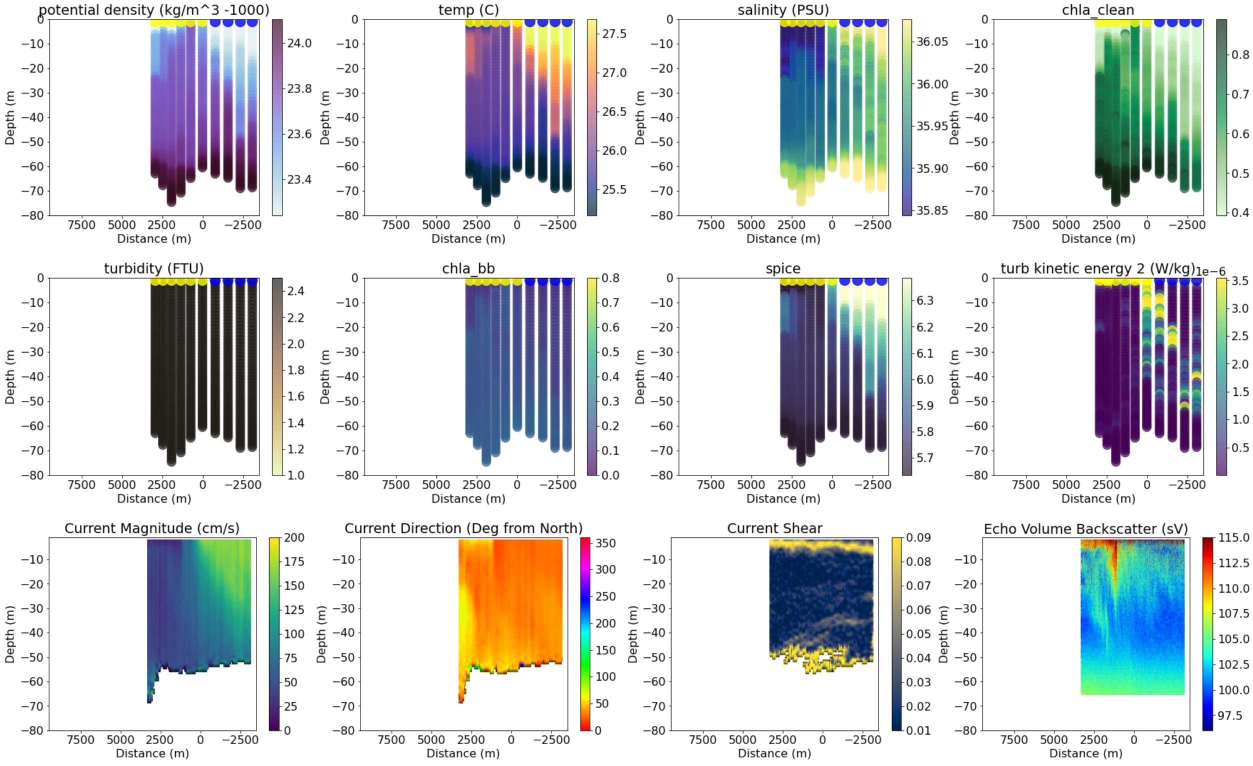

```
----------------------------------------
46
data/all_adcp/processed/101706_20211006T150751UTC_100s
Distance b/w GS and Coastal water is 22916.116300134763
```

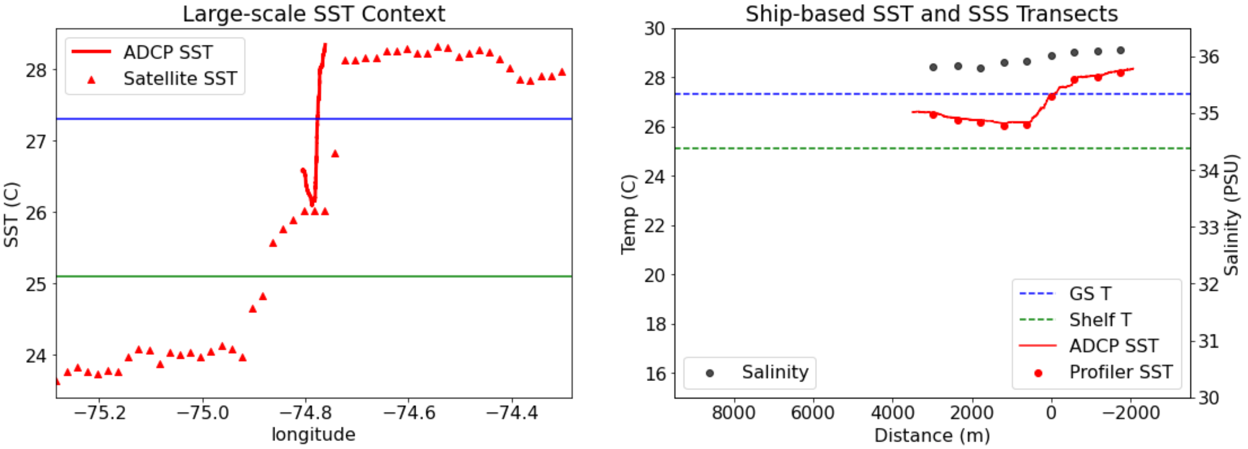

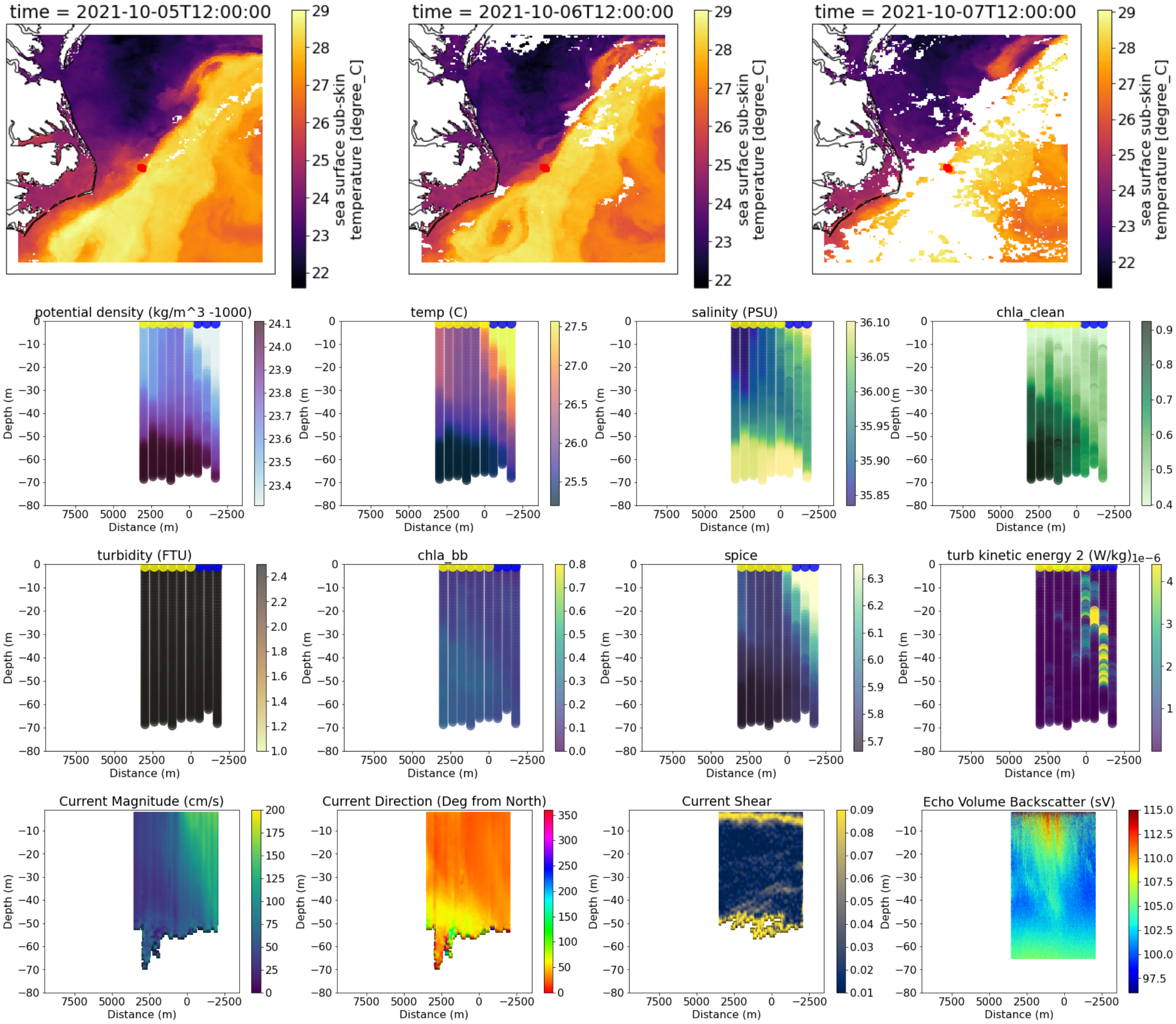


```
------------------------------------
48
data/all_adcp/processed/101706_20220131T132515UTC_100s
Distance b/w GS and Coastal water is 60098.31828756475
```

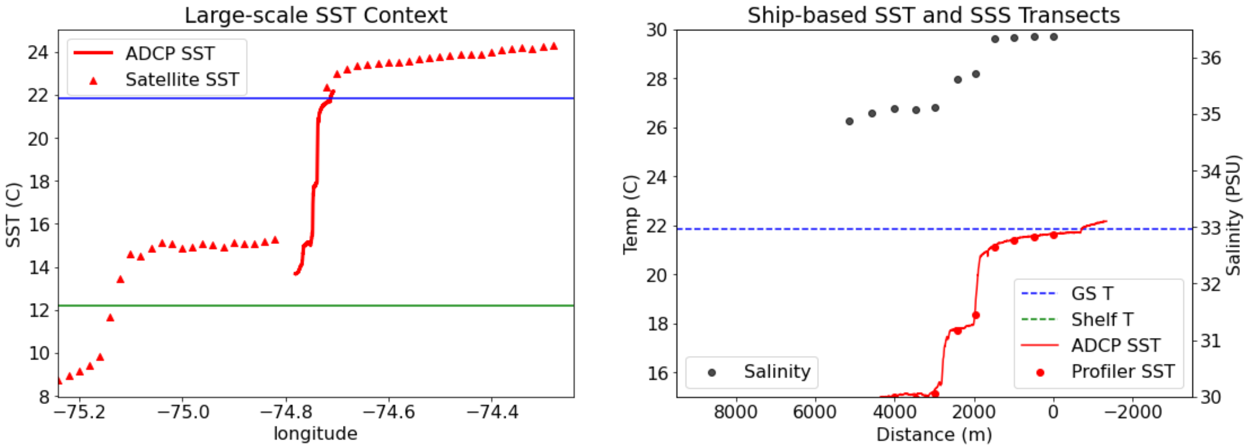

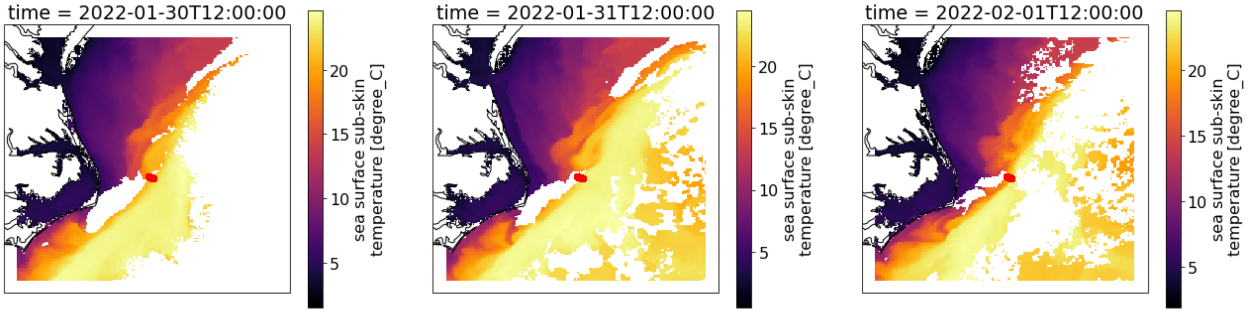


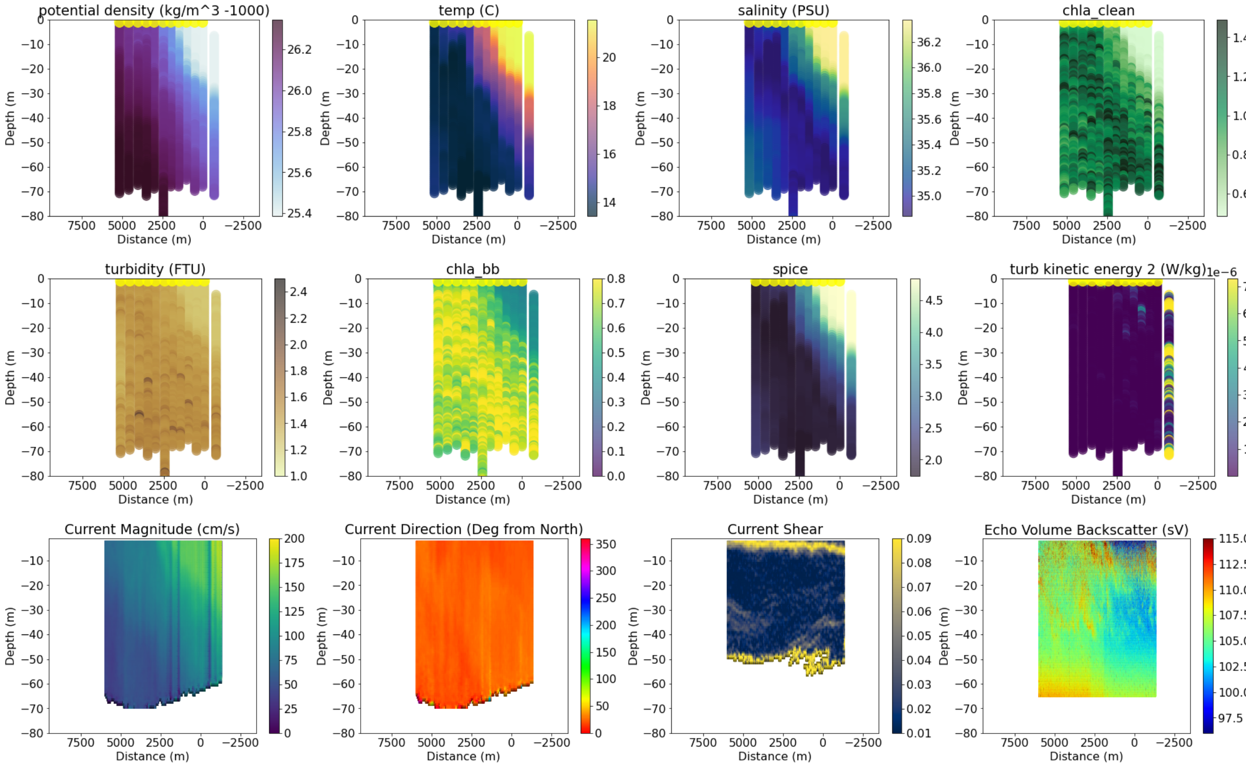


```
-----------------------------------
50
data/all_adcp/processed/101706_20220131T210615UTC_100s
Distance b/w GS and Coastal water is 57178.17716670775
```

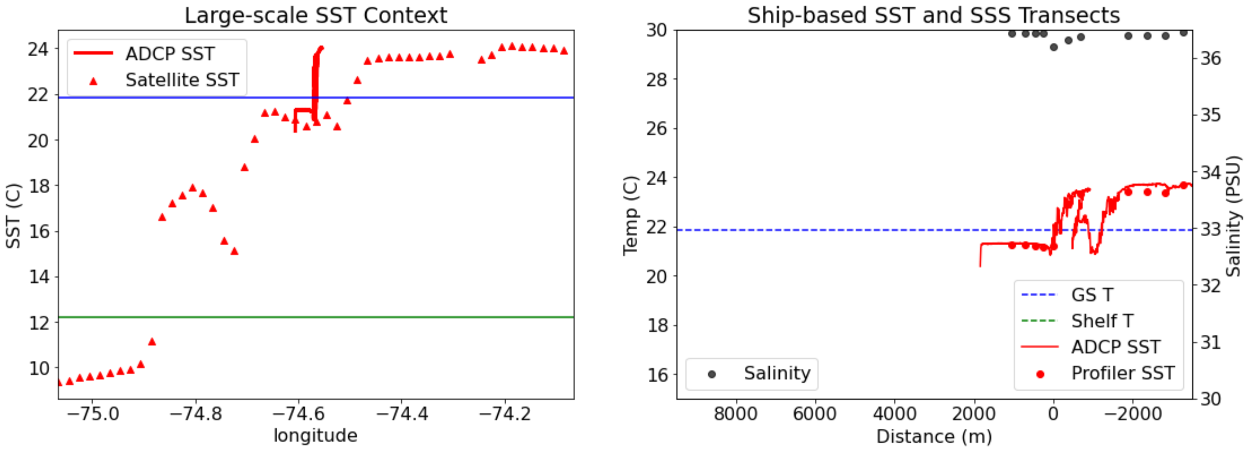

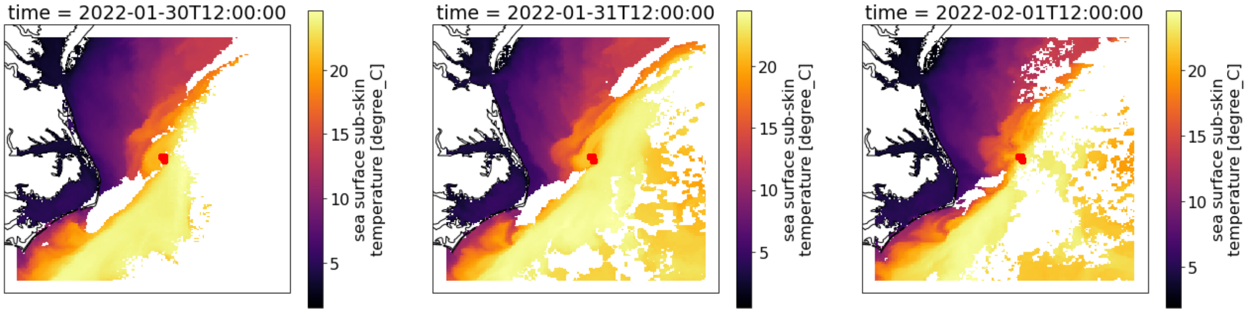


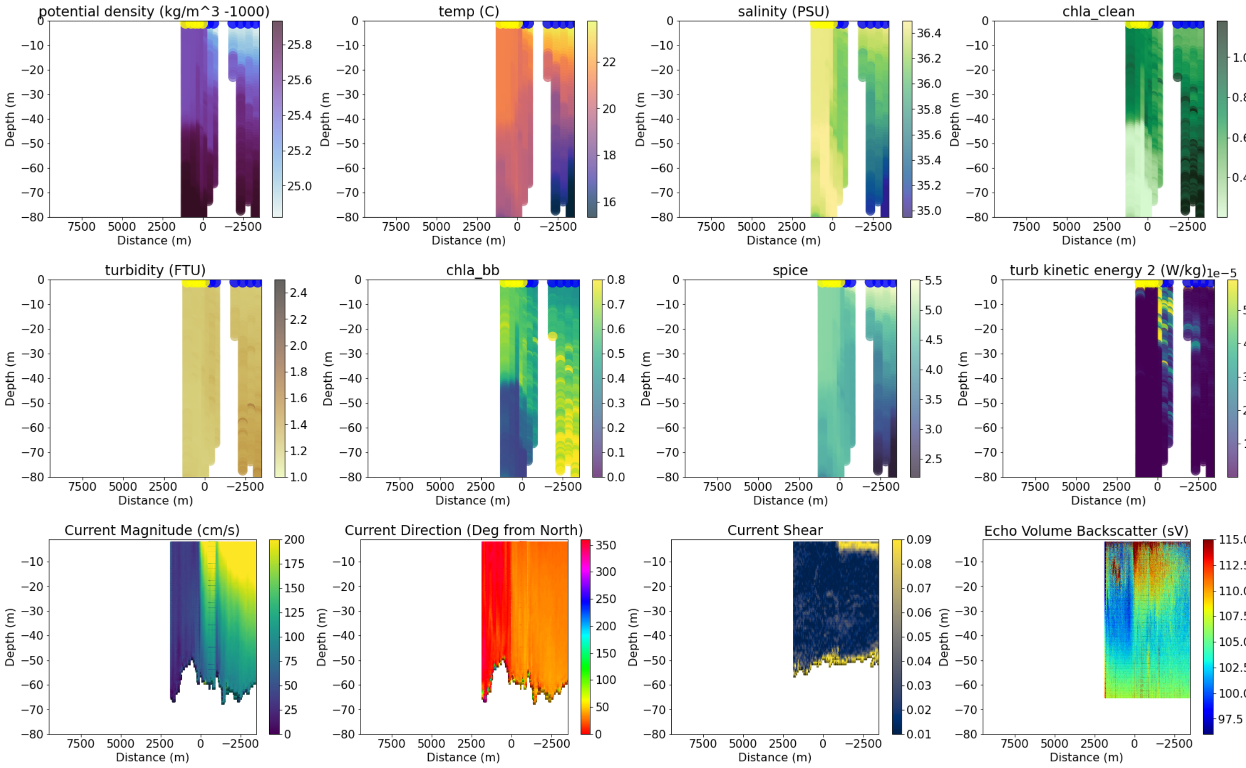


```
-----------------------------------
51
data/all_adcp/processed/101706_20220202T145622UTC_100s
Distance b/w GS and Coastal water is 42908.2093070654
```

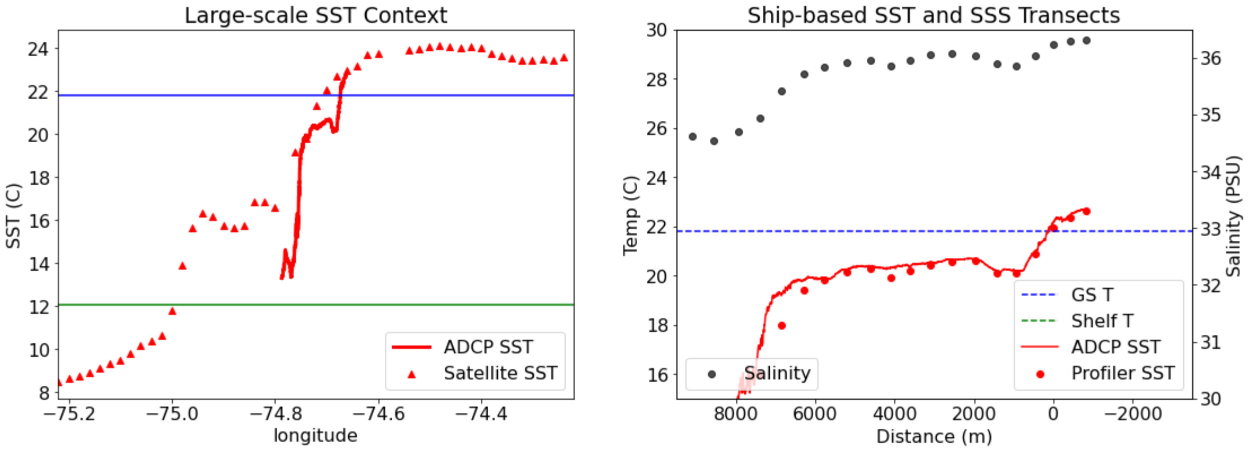

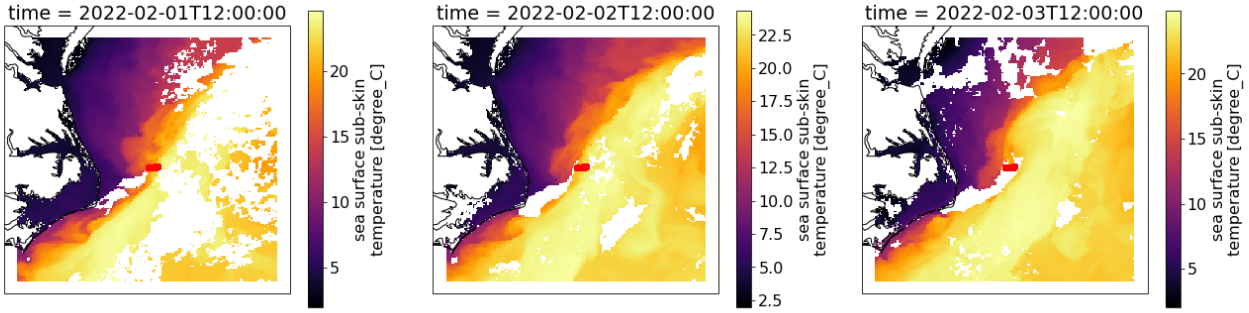


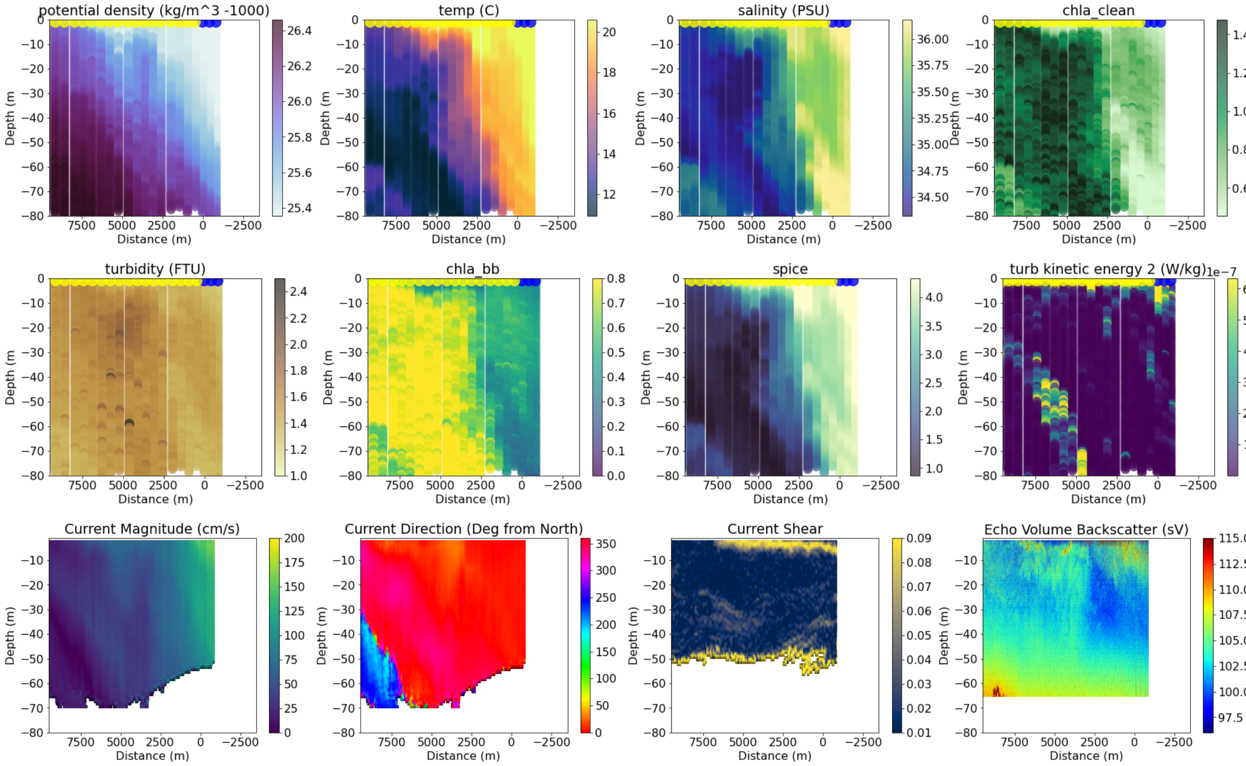


```
------------------------------------
52
data/all_adcp/processed/101706_20220202T192604UTC_100s
Distance b/w GS and Coastal water is 37179.4255283457
```

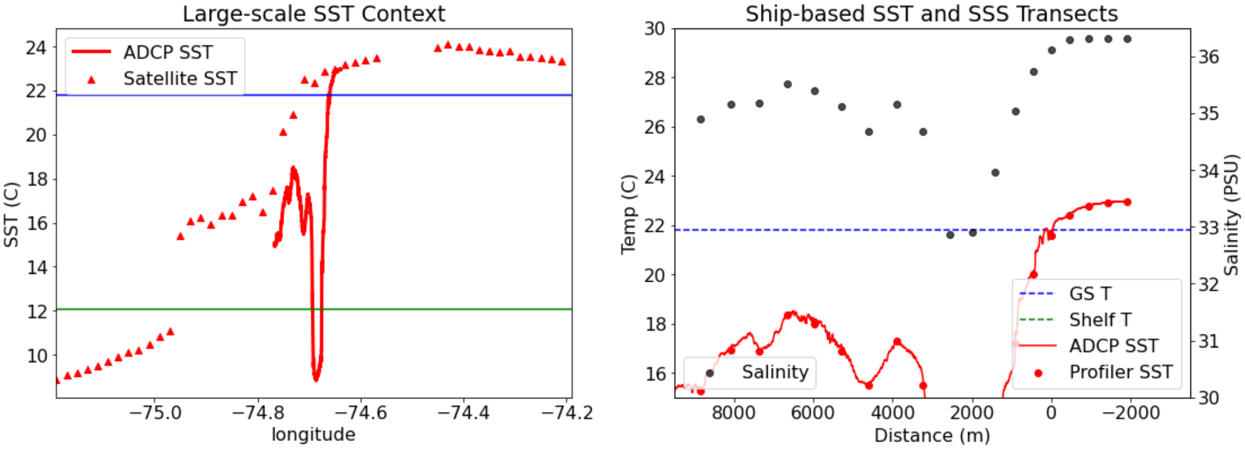

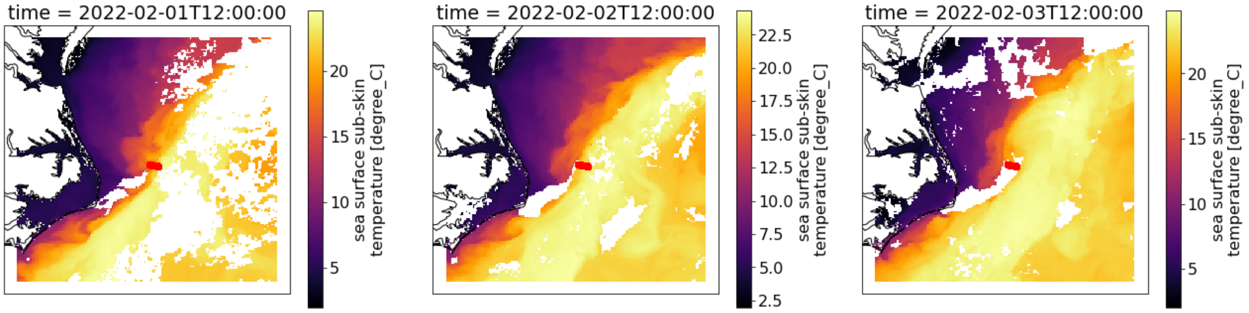


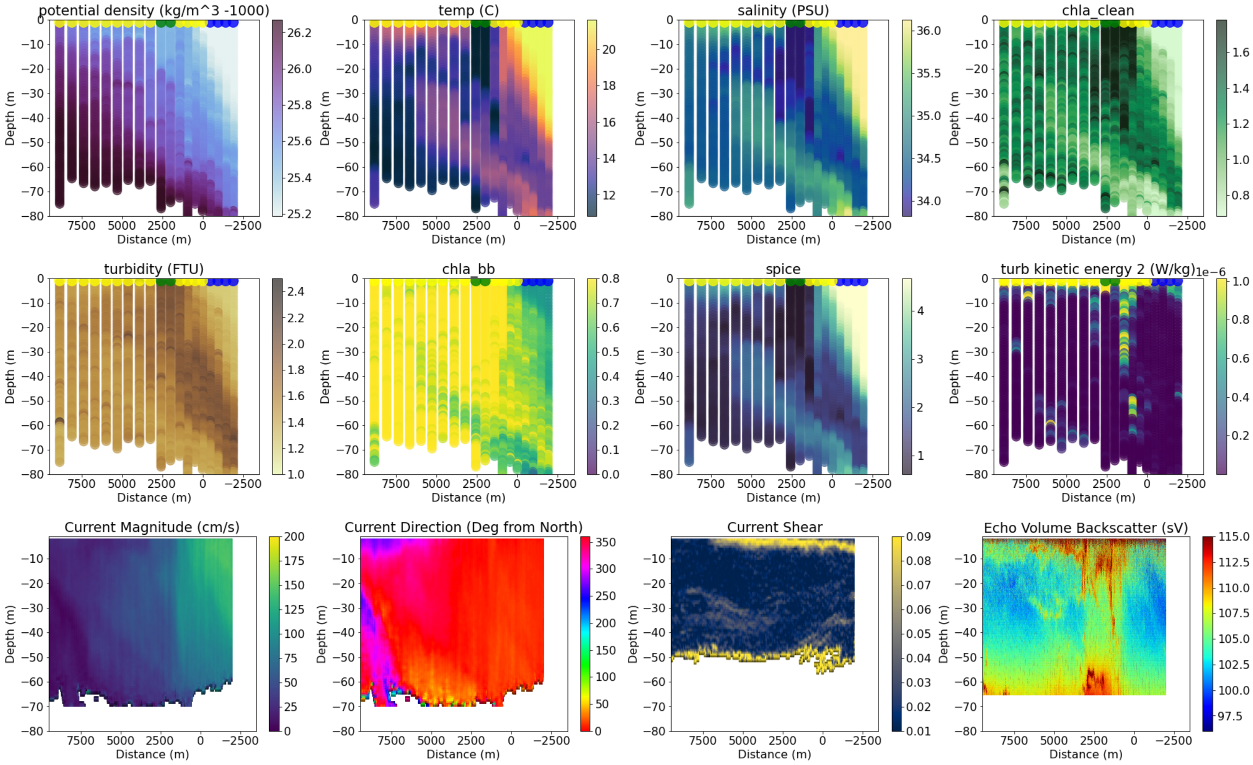


```
--------------------------------------
53
data/all_adcp/processed/101706_20220309T151806UTC_100s
Distance b/w GS and Coastal water is 100148.57675680815
```

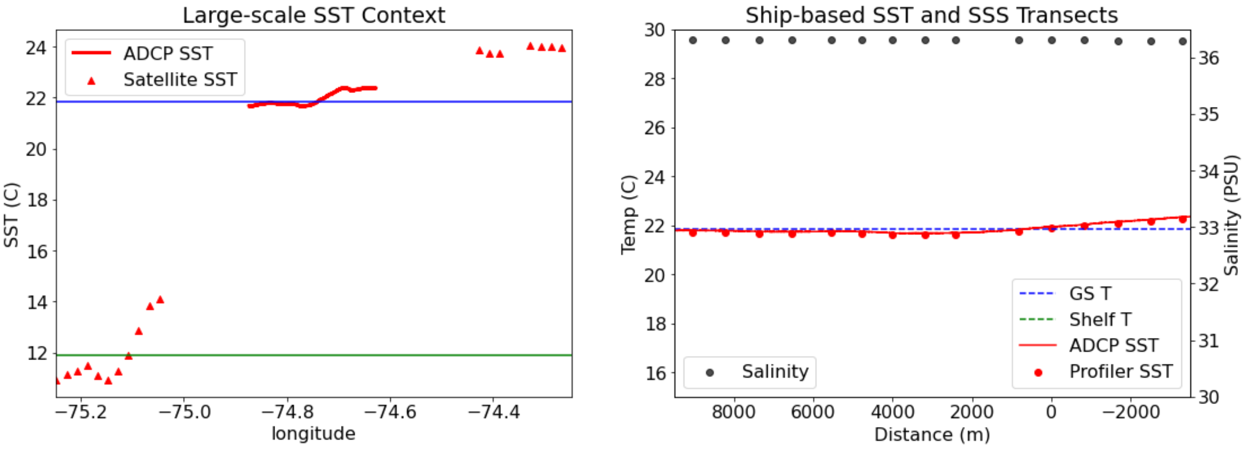

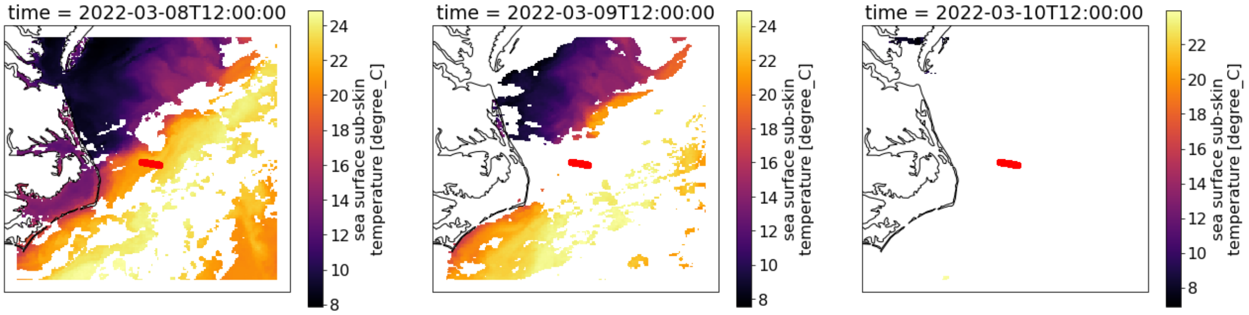


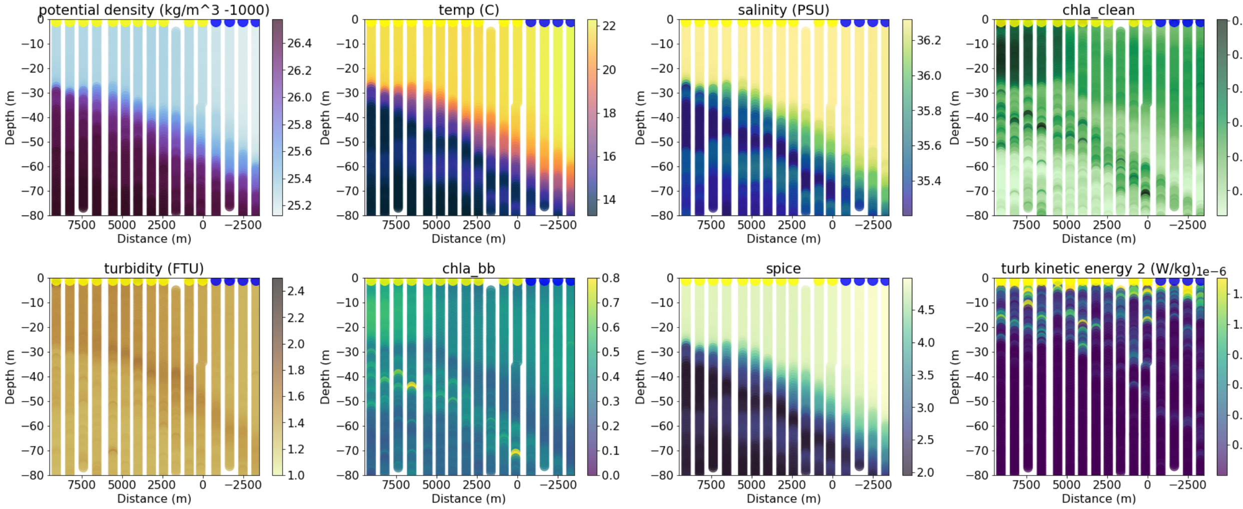


```
Incompatible X, Y inputs to pcolormesh; see help(pcolormesh)
54
data/all_adcp/processed/101706_20220309T203937UTC_100s
Transect does not cross into coastal water
```

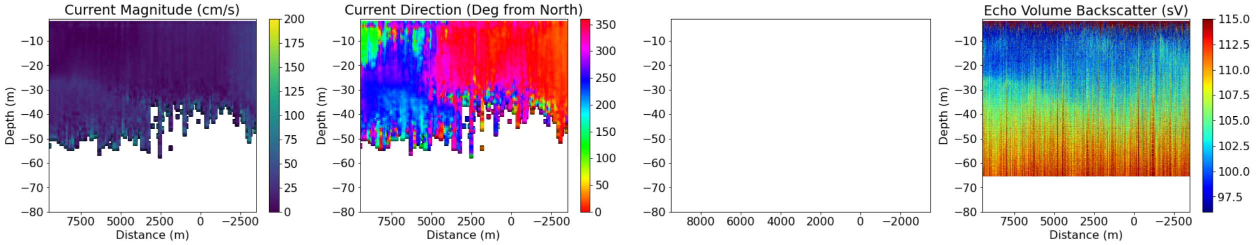


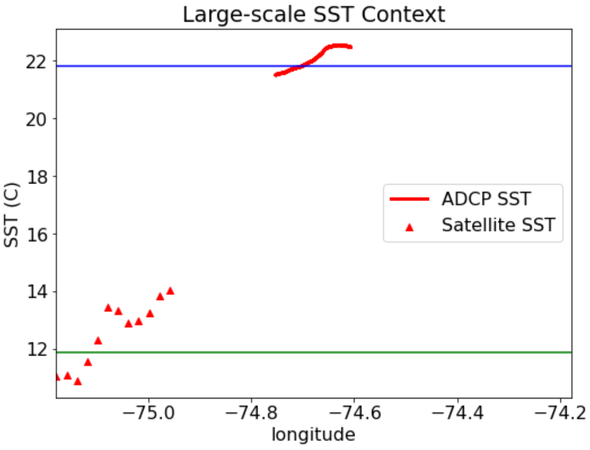


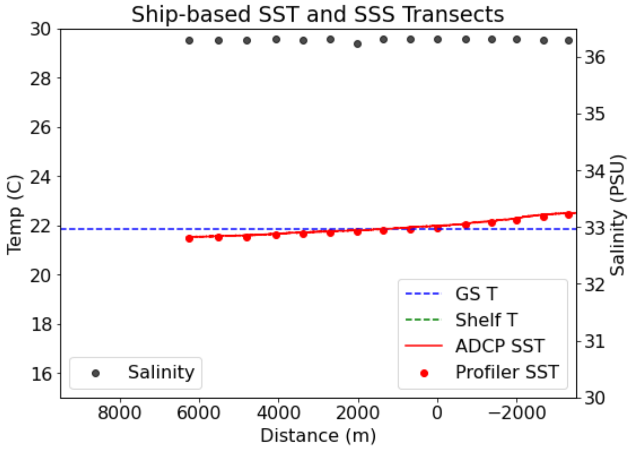

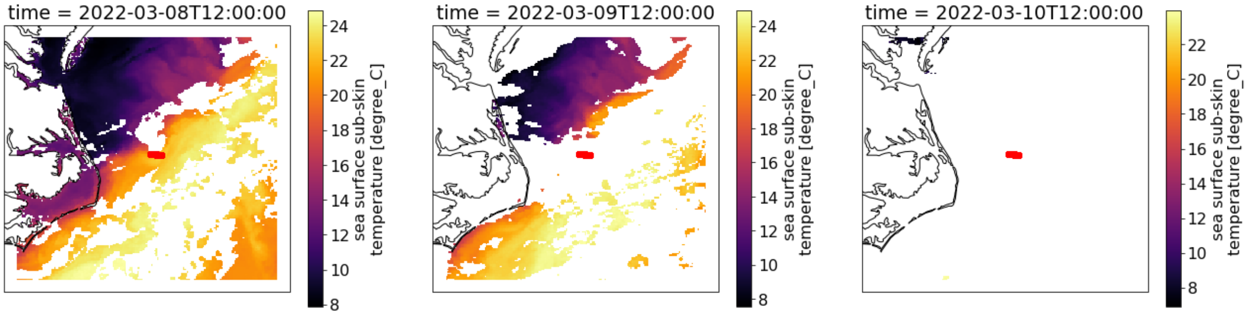


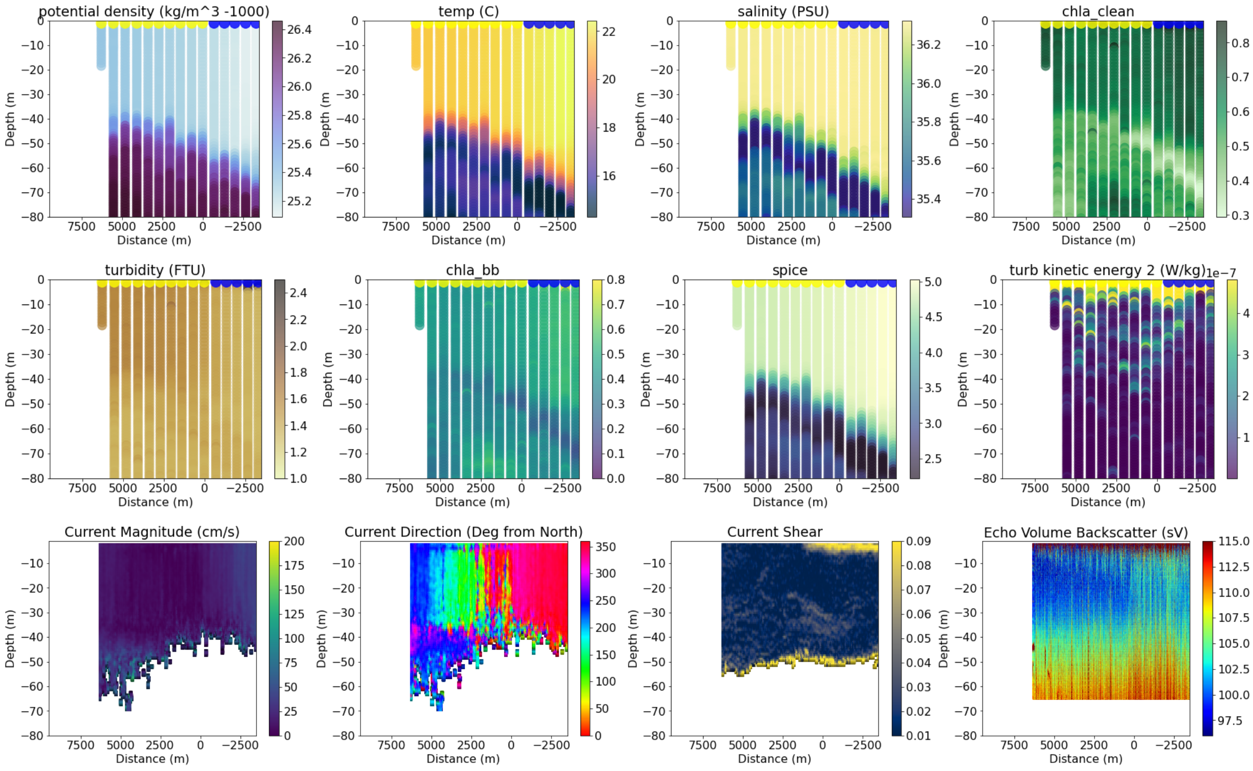


--------------------------------------
55
data/all_adcp/processed/101706_20220310T130351UTC_100s
Transect does not cross into coastal water

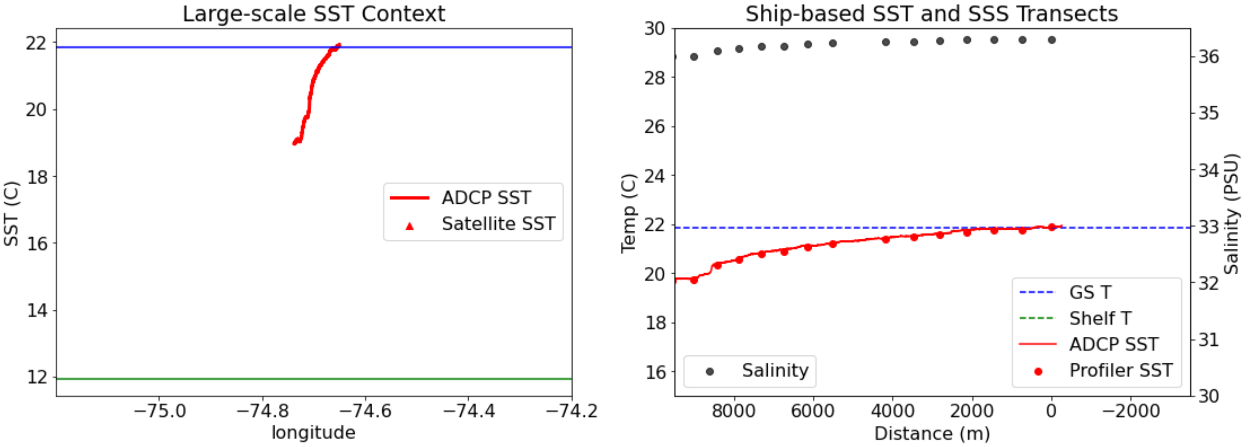

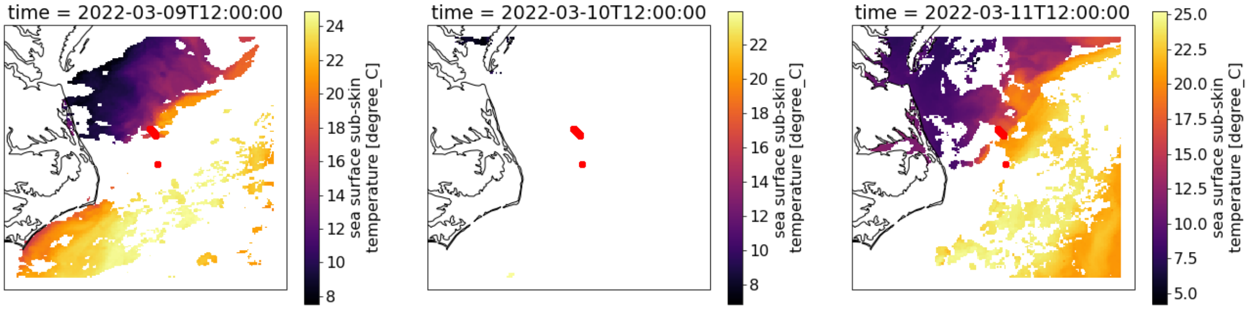


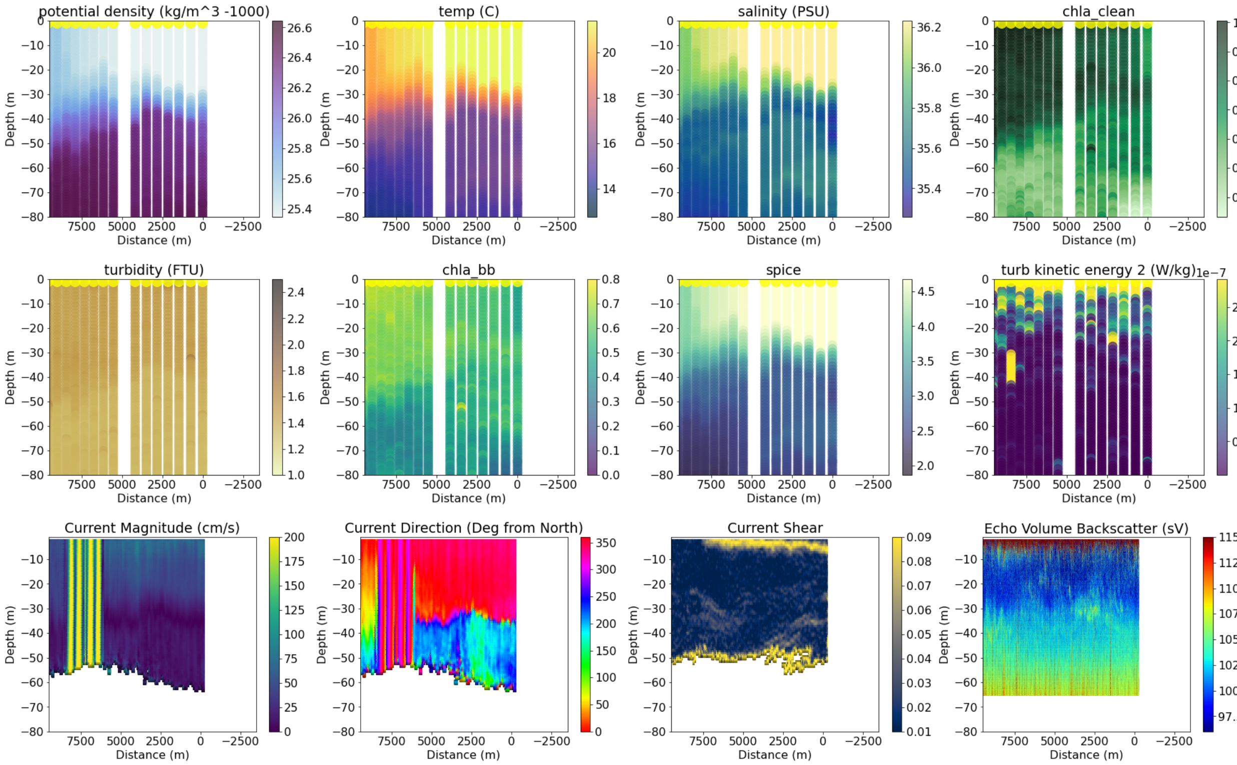


```
-----------------------------------
57
data/all_adcp/processed/101706_20220311T150047UTC_100s
Distance b/w GS and Coastal water is 126006.79679906329
```

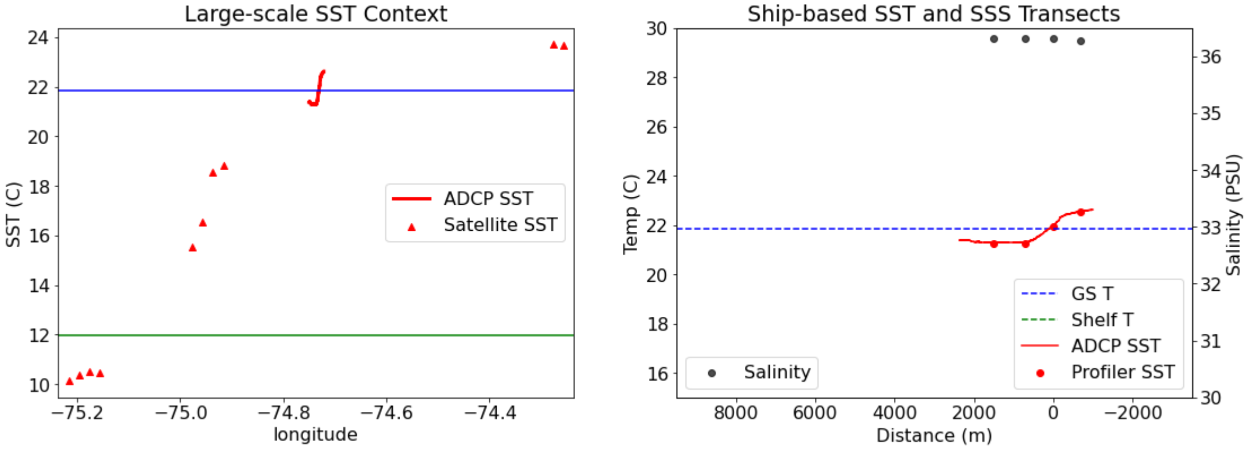

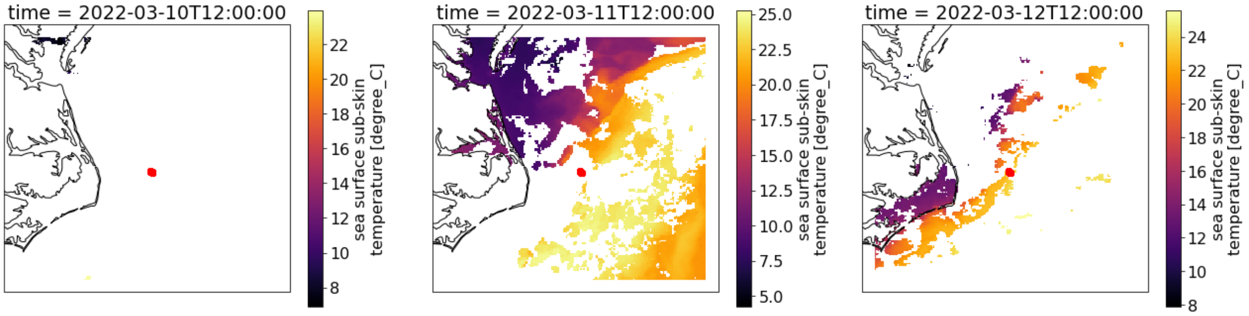


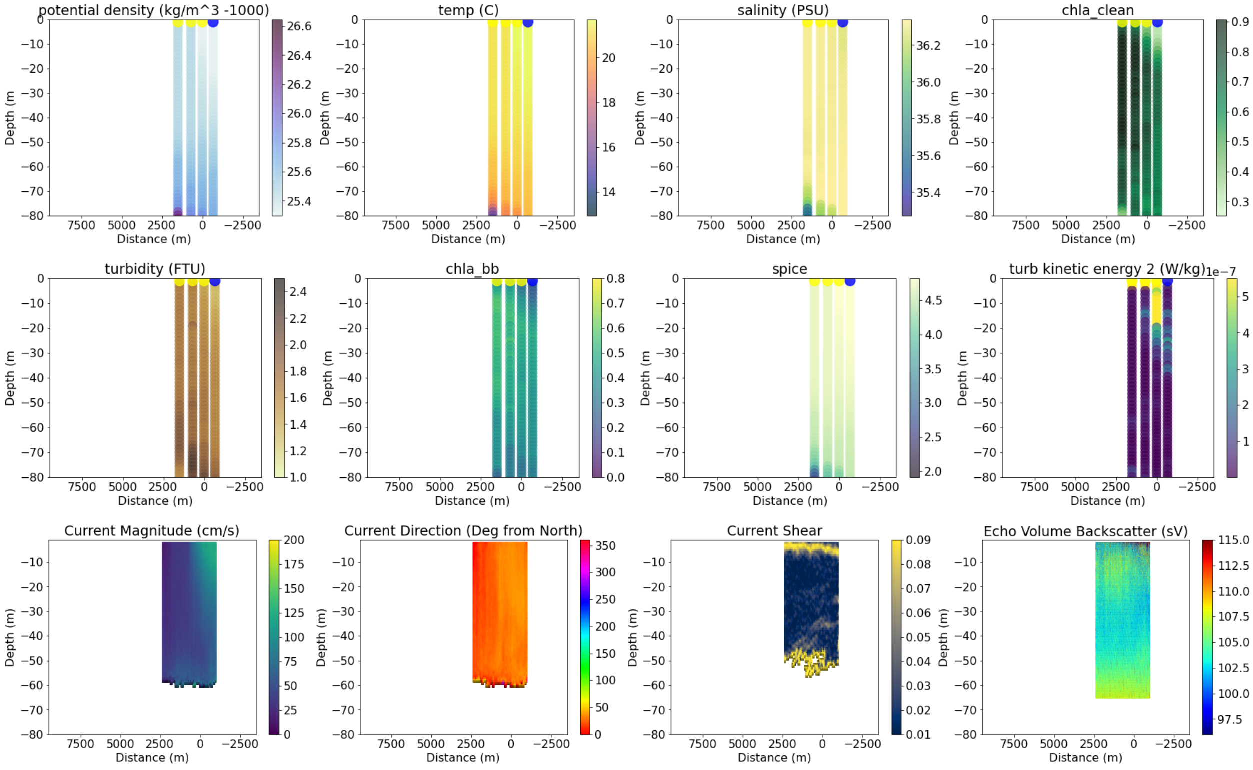


```
-----------------------------------------
58
data/all_adcp/processed/101706_20220311T160718UTC_100s
Distance b/w GS and Coastal water is 126005.30407021676
```

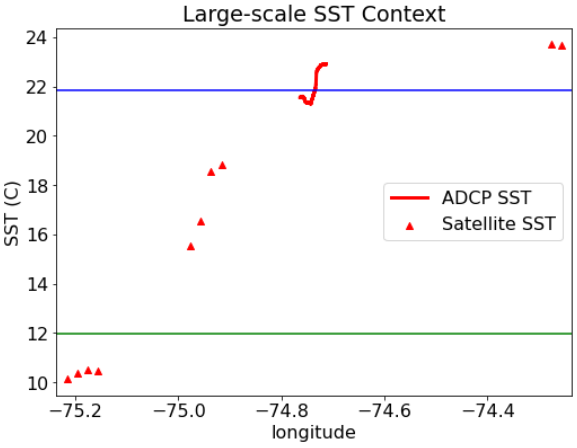


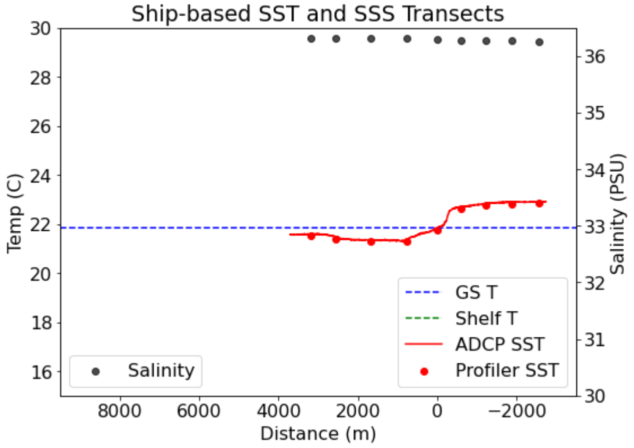

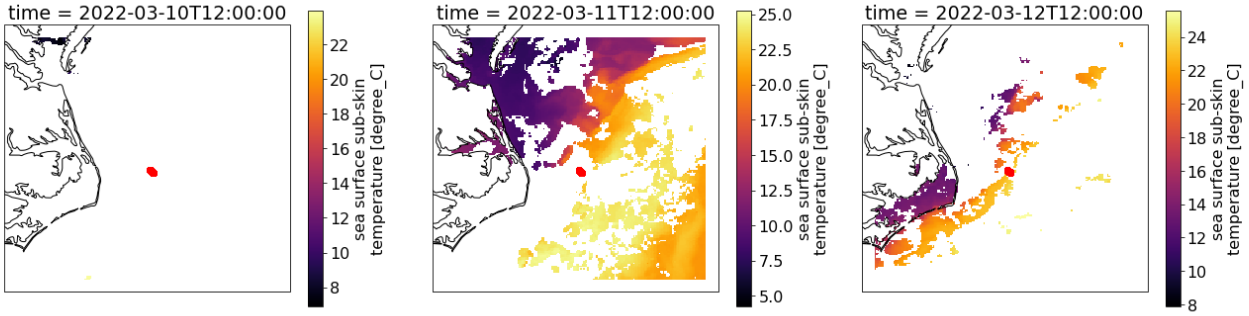


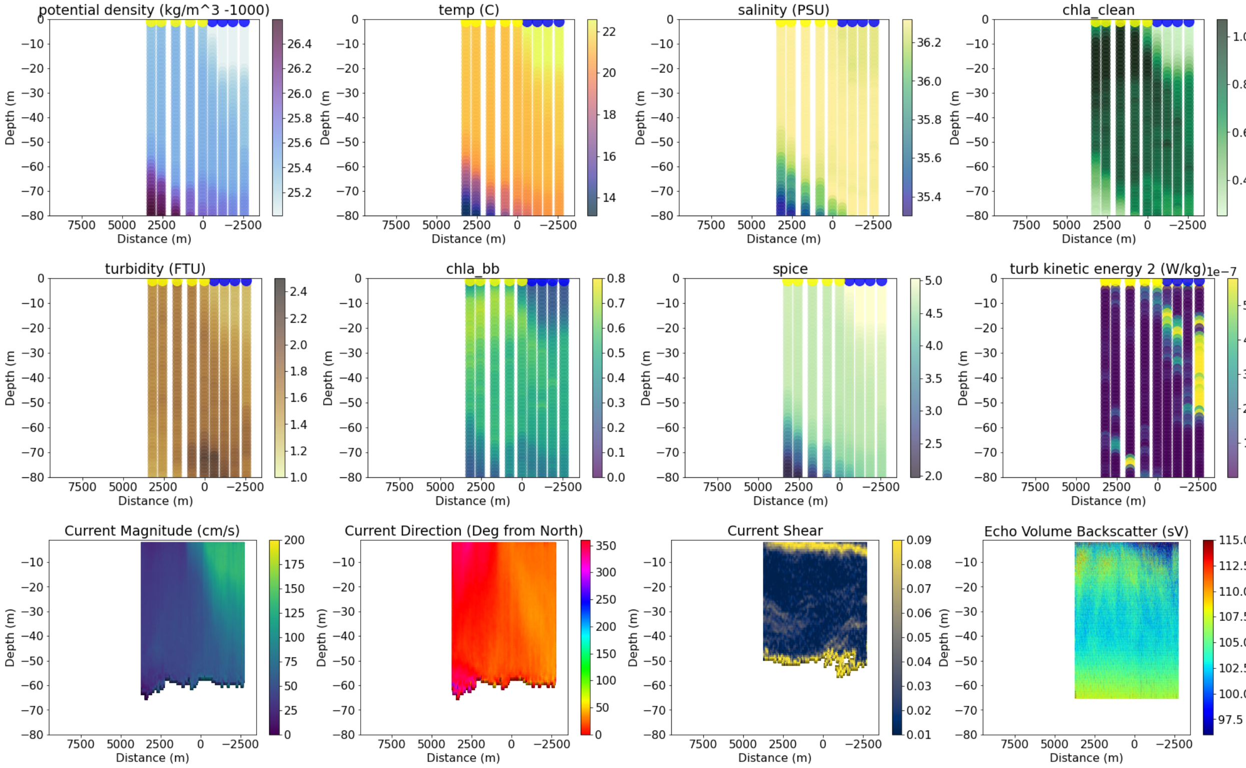


```
------------------------------------
59
data/all_adcp/processed/101706_20220403T114937UTC_100s
Distance b/w GS and Coastal water is 65744.1976264358
```

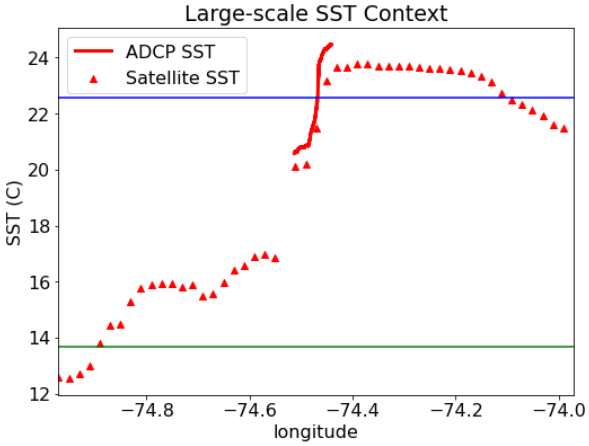


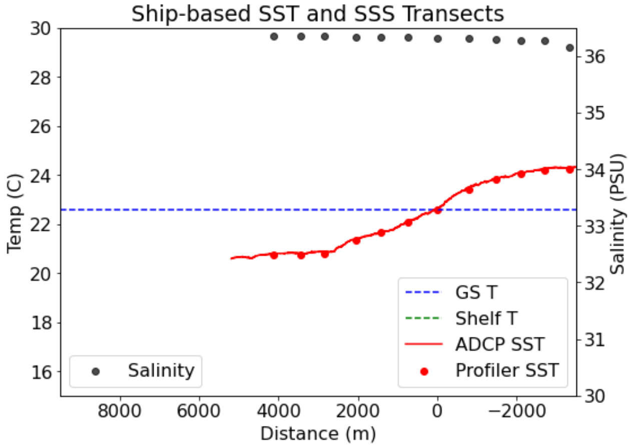

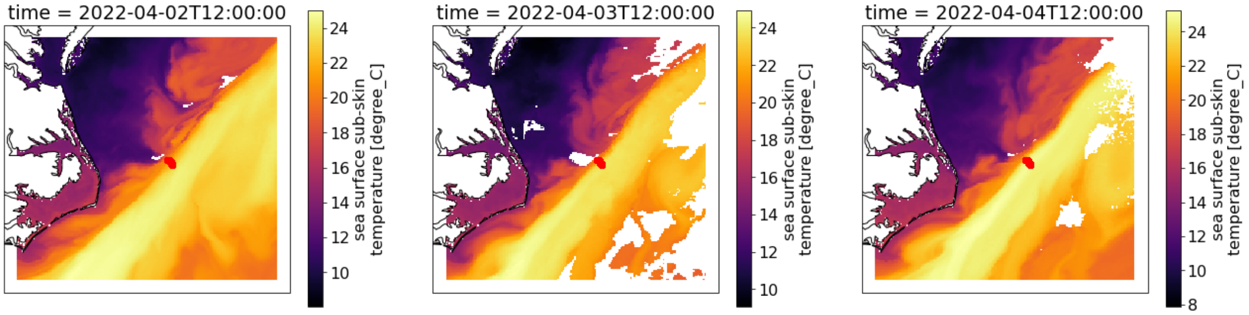


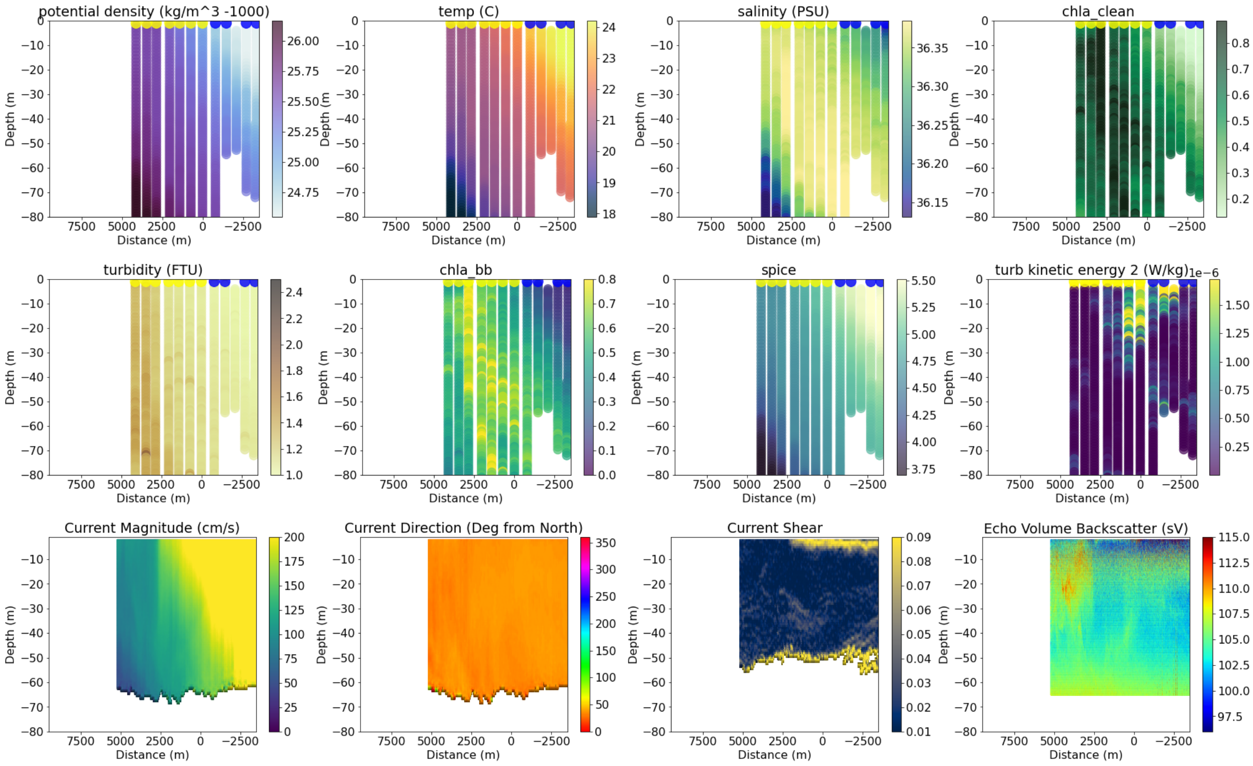


```
-----------------------------------
63
data/all_adcp/processed/101706_20220404T120954UTC_100s
Distance b/w GS and Coastal water is 25758.12928412933
```

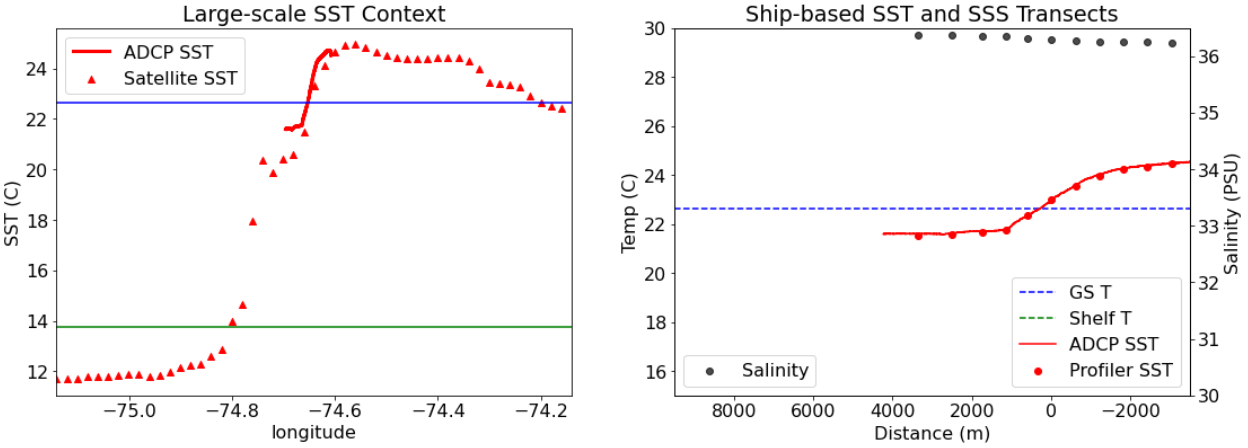

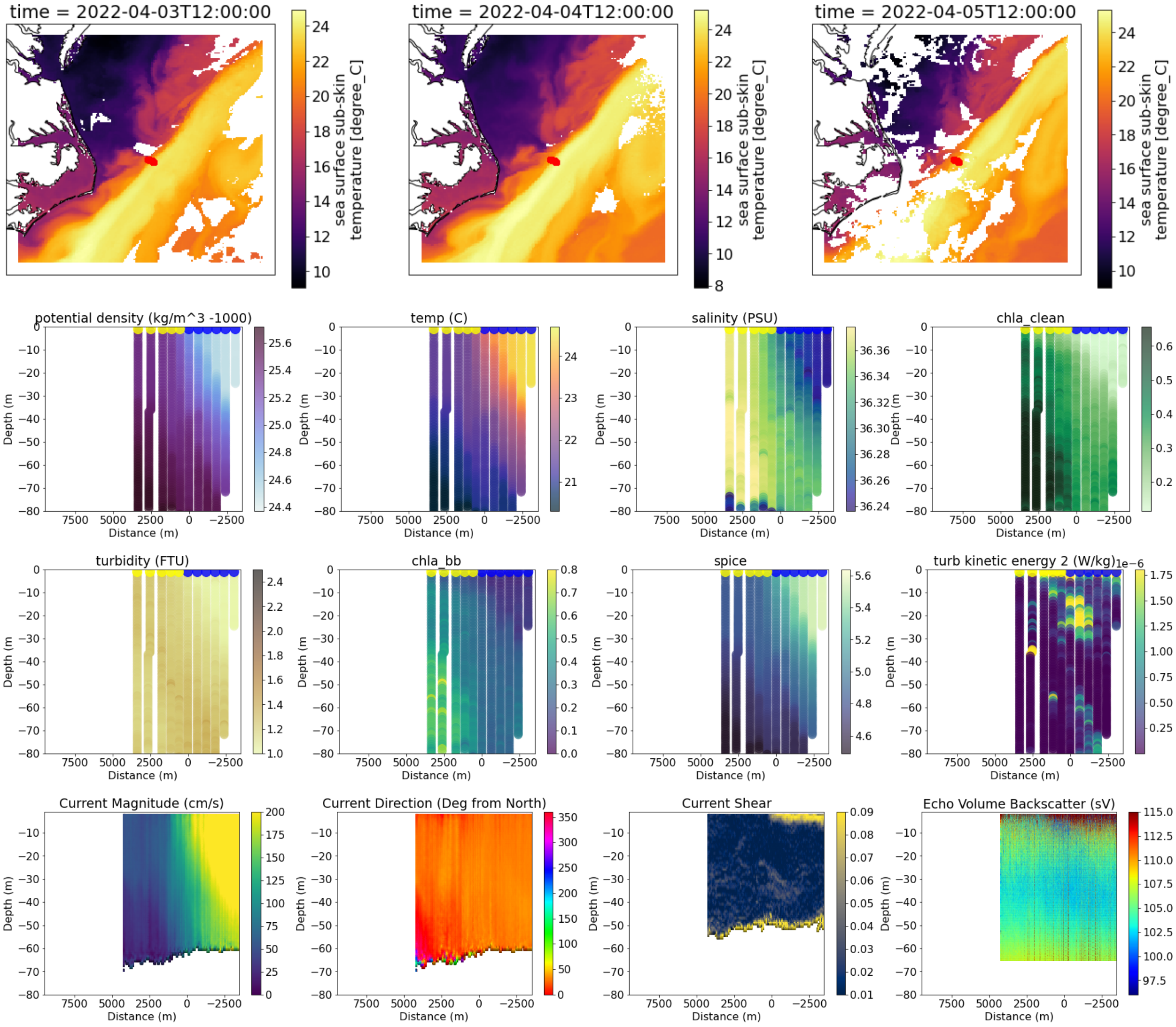


```
------------------------------------
64
data/all_adcp/processed/101706_20220404T145648UTC_100s
Distance b/w GS and Coastal water is 45752.21456440216
```

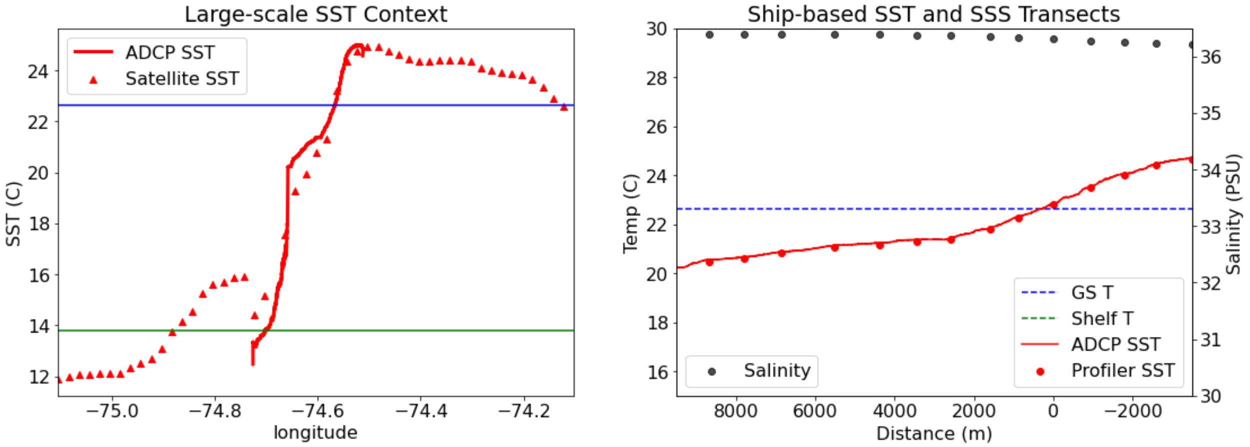

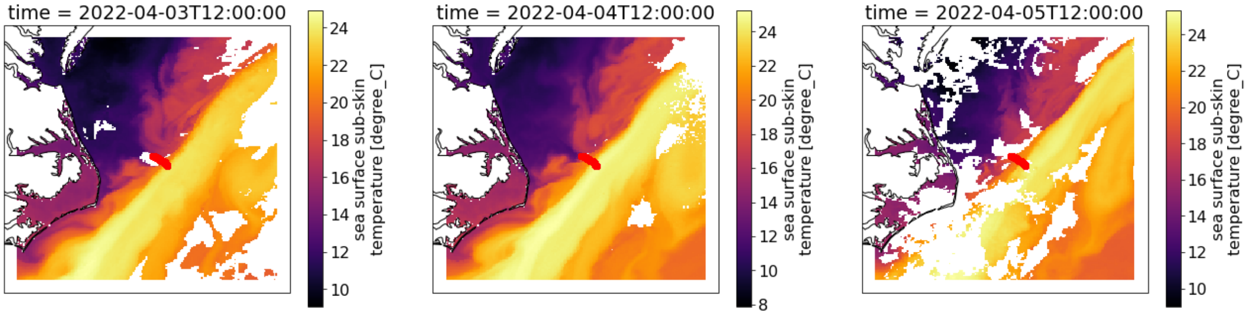


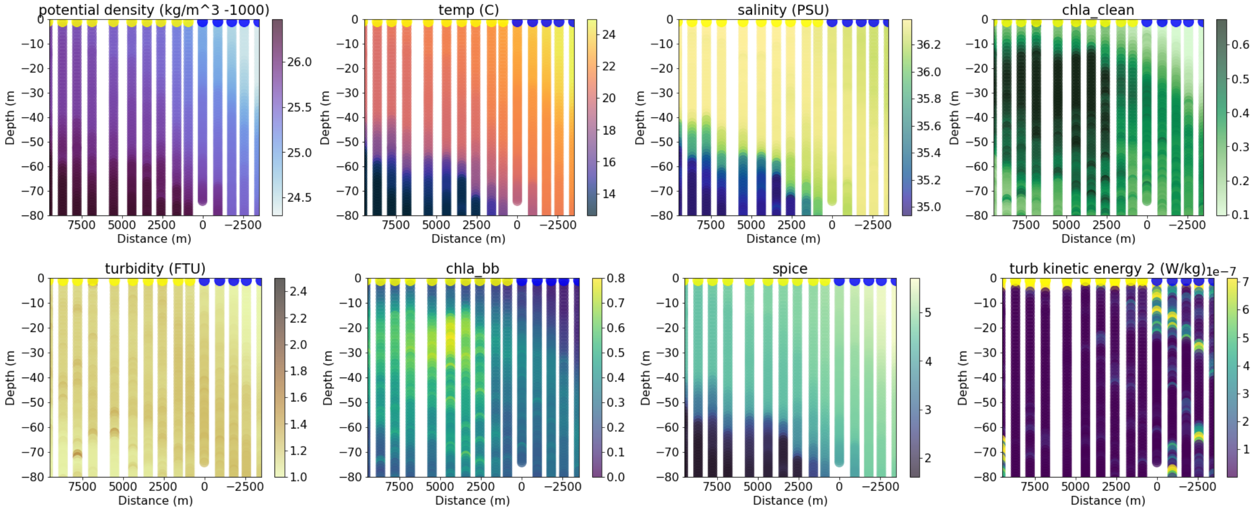


```
Incompatible X, Y inputs to pcolormesh; see help(pcolormesh)
65
data/all_adcp/processed/101706_20220404T194713UTC_100s
Distance b/w GS and Coastal water is 60002.13618931359
```

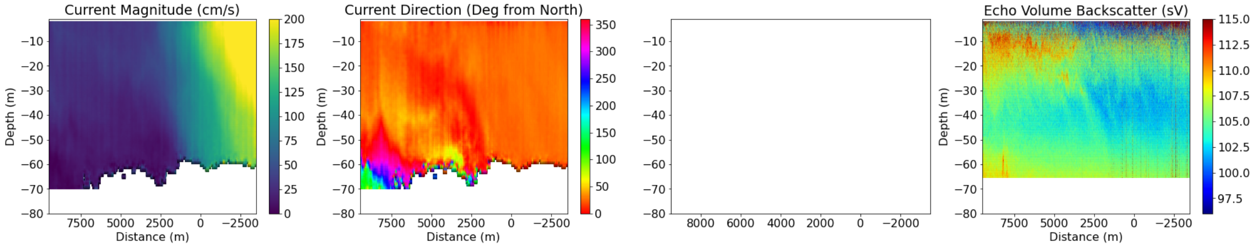


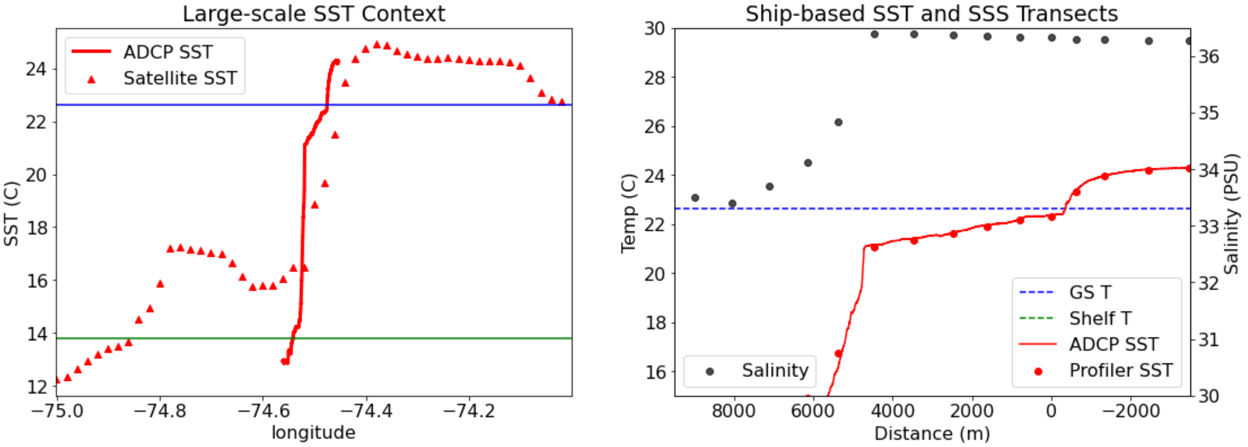

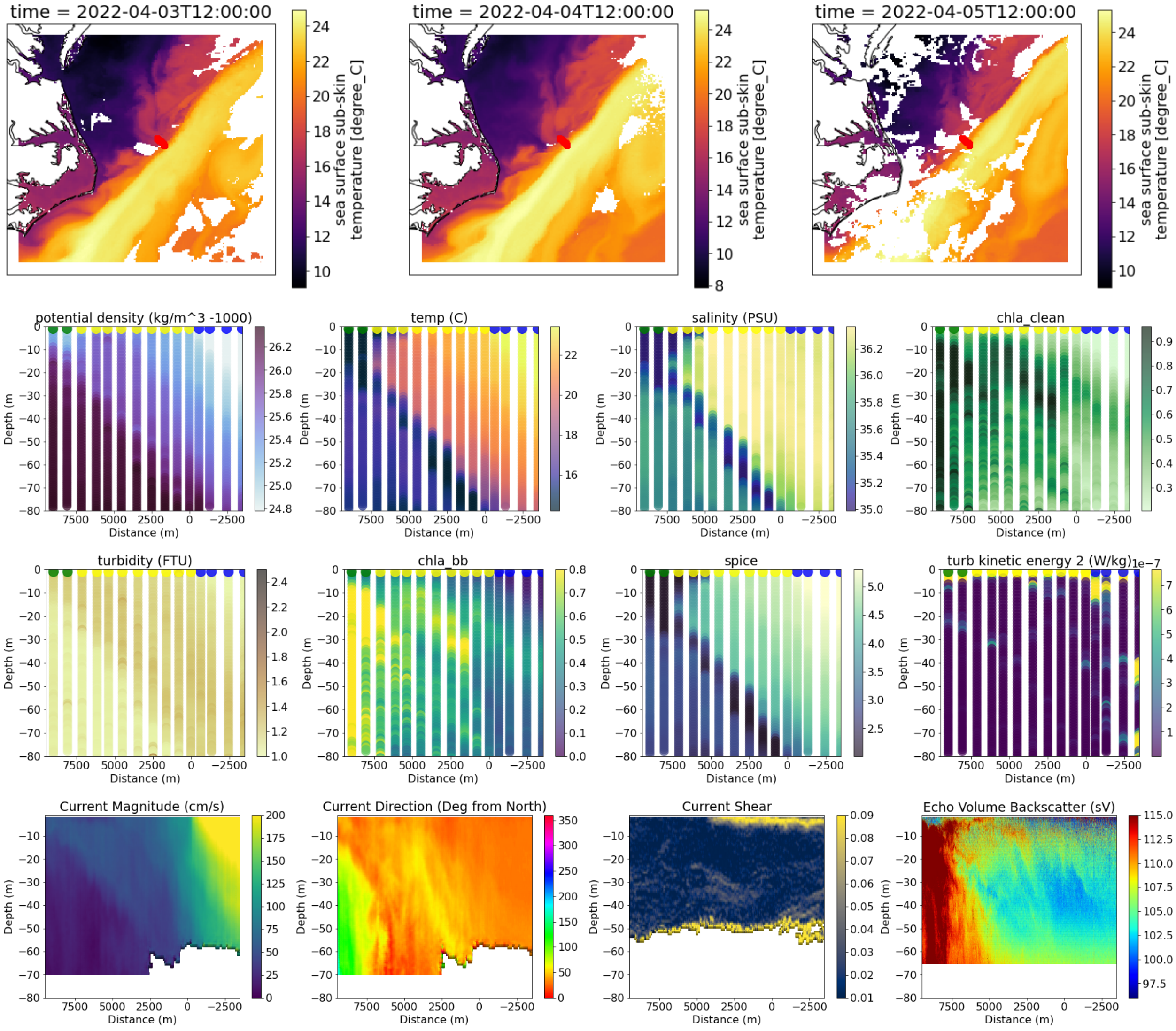

```
----------------------------------------
66
data/all_adcp/processed/101706_20220405T120045UTC_100s
Distance b/w GS and Coastal water is 40078.59380967104
```

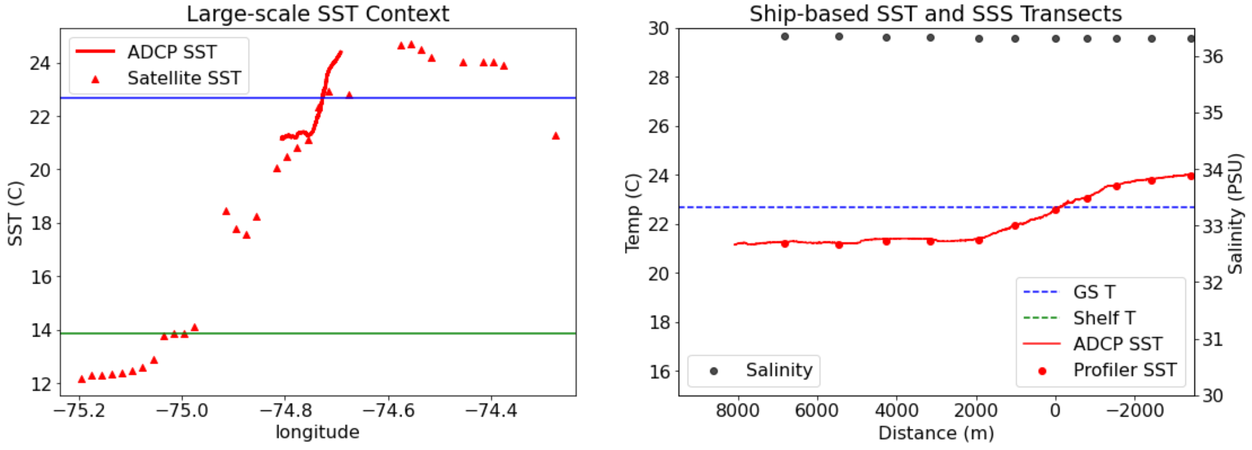

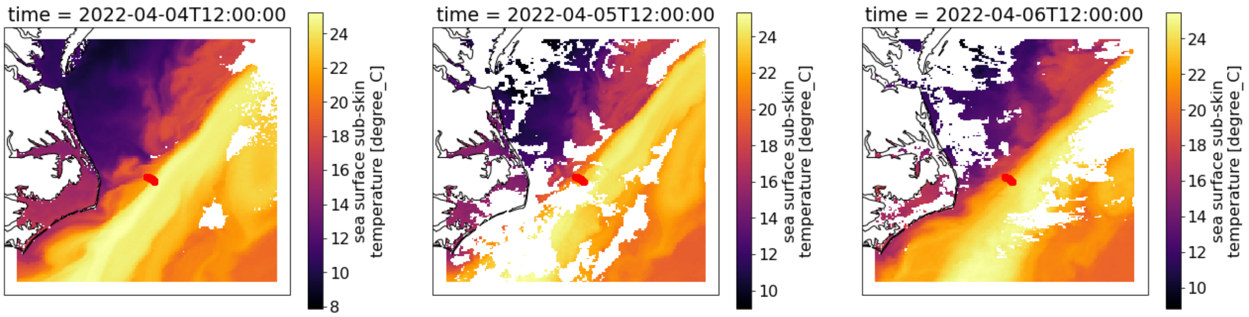


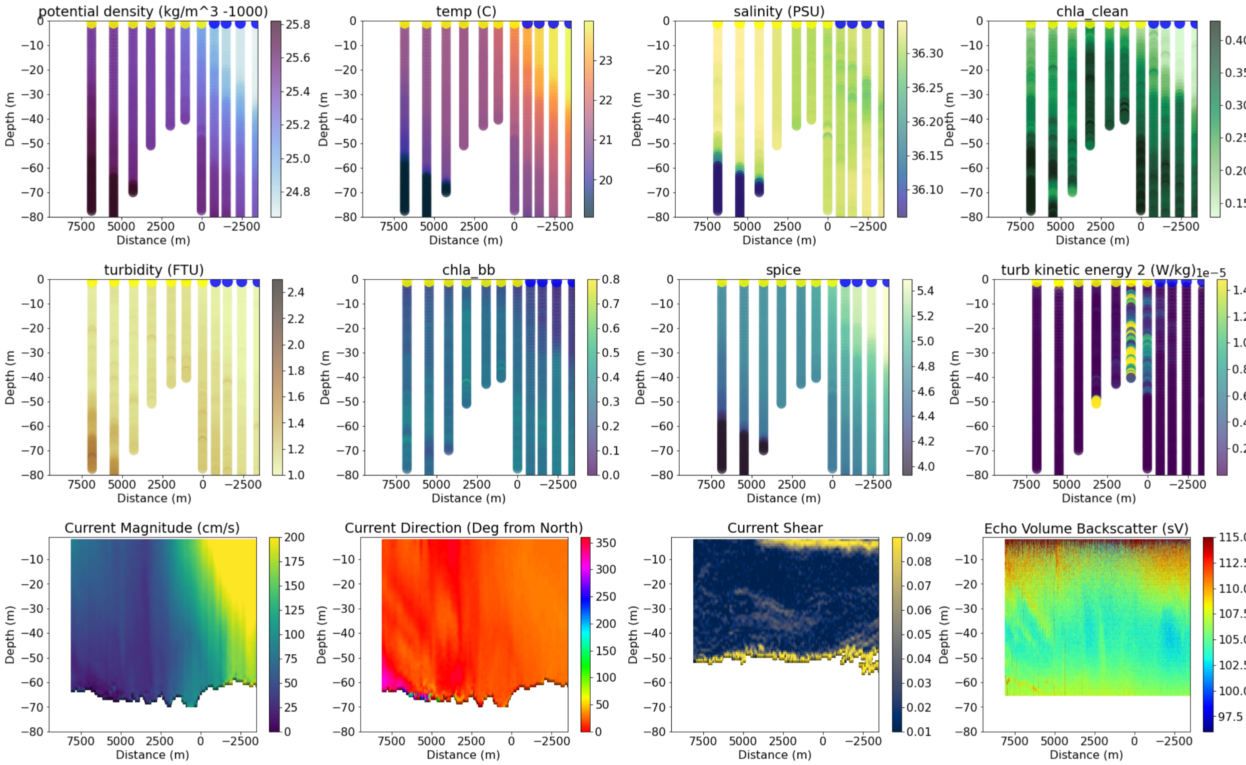


```
-------------------------------------
67
data/all_adcp/processed/101706_20220405T150549UTC_100s
Distance b/w GS and Coastal water is 51489.29022644355
```

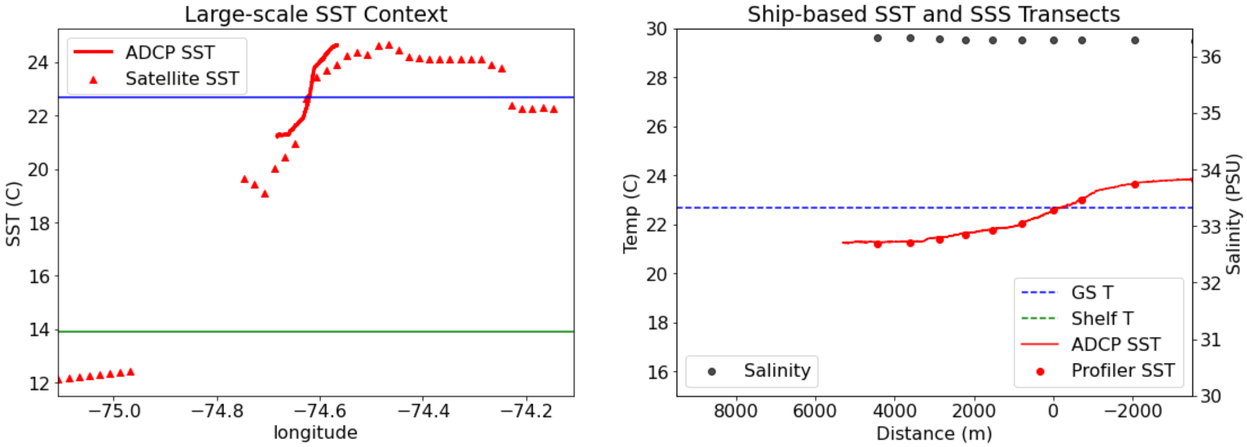

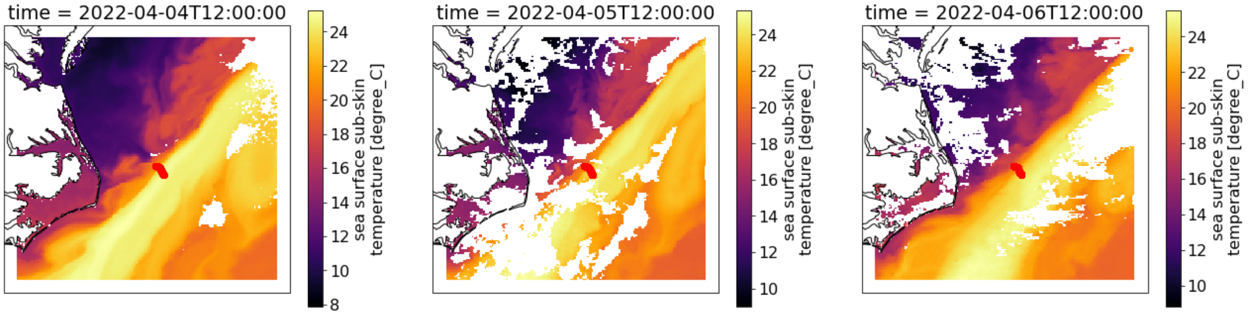


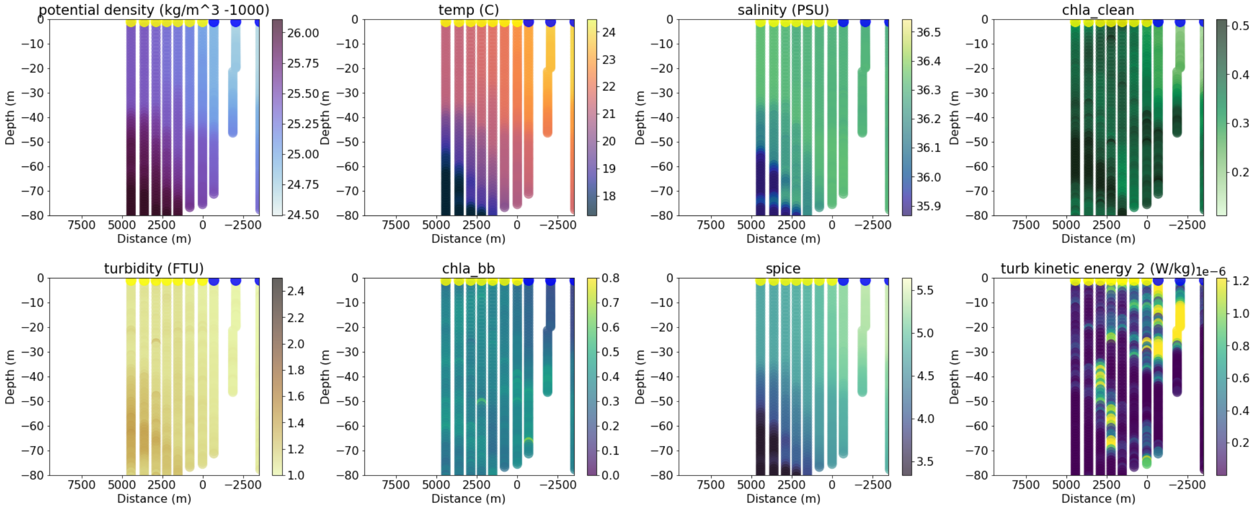


```
Incompatible X, Y inputs to pcolormesh; see help(pcolormesh)
68
data/all_adcp/processed/101706_20220405T184629UTC_100s
Distance b/w GS and Coastal water is 57181.3328640114
```

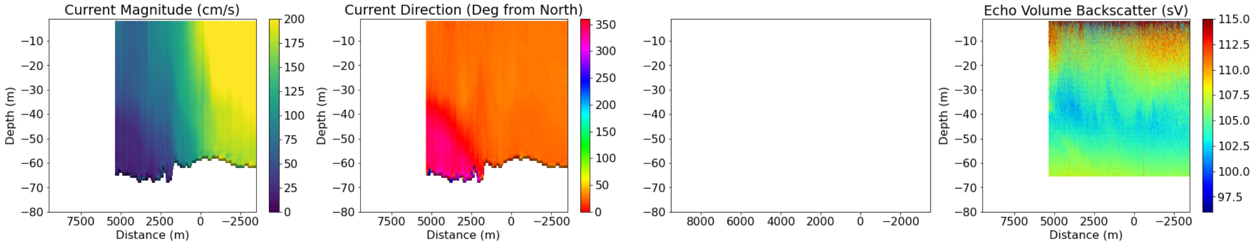


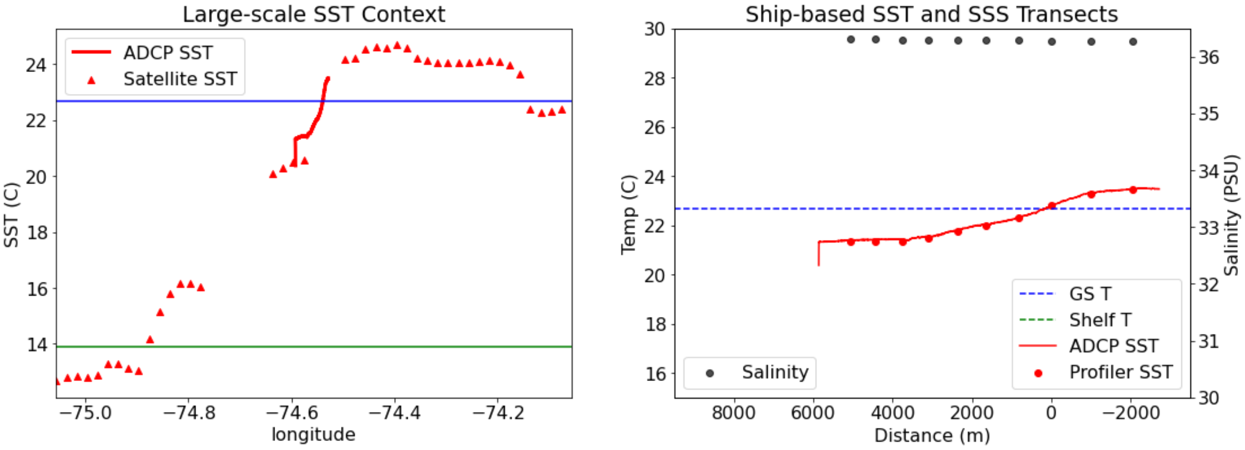

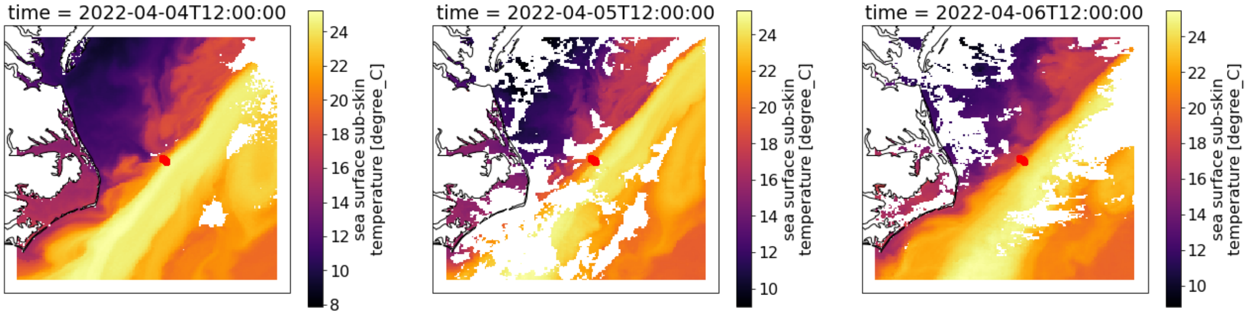
time = 2022-04-04T12:00:00
time = 2022-04-05T12:00:00
time = 2022-04-06T12:00:00
sea surface sub-skin temperature [degree_C]

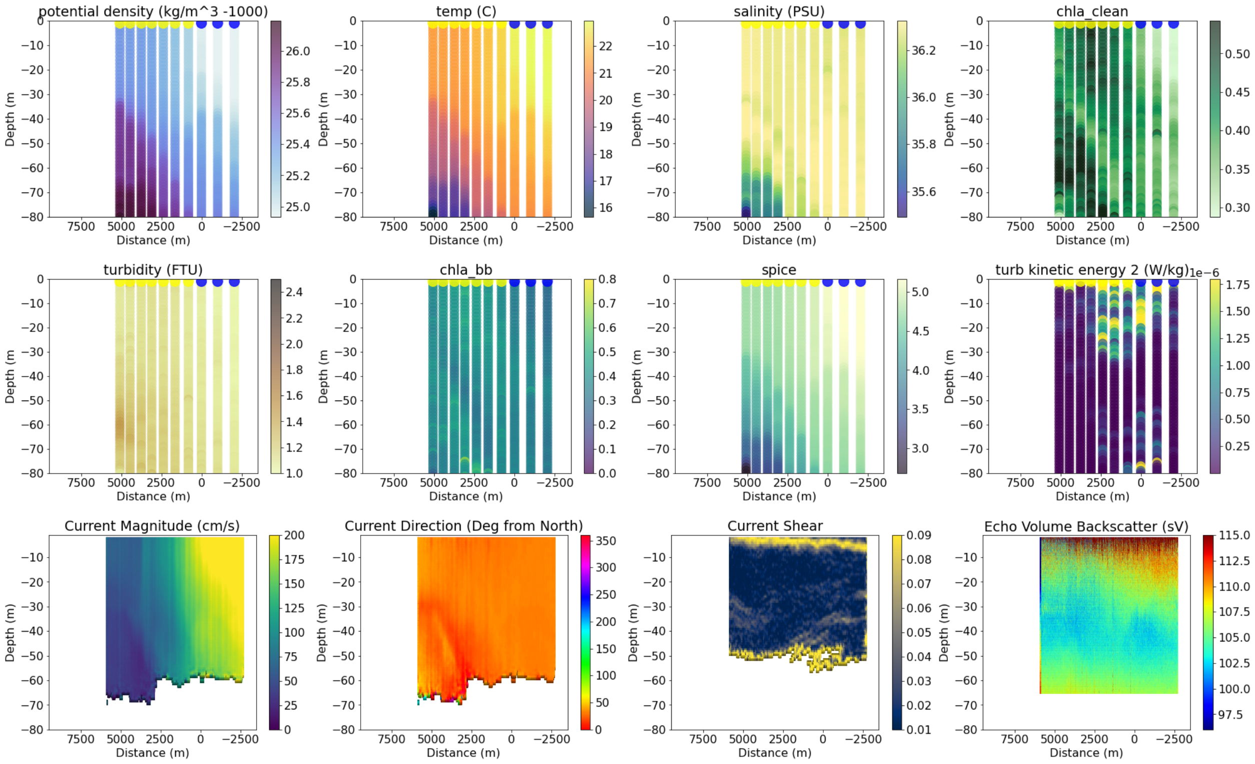
potential density (kg/m^3 -1000)
temp (C)
salinity (PSU)
chla_clean
turbidity (FTU)
chla_bb
spice
turb kinetic energy 2 (W/kg)
1e−6
Current Magnitude (cm/s)
Current Direction (Deg from North)
Current Shear
Echo Volume Backscatter (sV)
Depth (m)
Distance (m)

```
----------------------------------------
```
